# Genomic analyses capture the human-induced demographic collapse and recovery in a wide-ranging cervid

**DOI:** 10.1101/2023.07.19.549627

**Authors:** Camille Kessler, Aaron B.A. Shafer

## Abstract

The glacial cycles of the Quaternary heavily impacted species through successions of population contractions and expansions. Similarly, populations have been intensely shaped by human pressures such as unregulated hunting and land use changes. White-tailed and mule deer survived in different refugia through the Last Glacial Maximum, and their populations were severely reduced after the European colonisation. Here, we analysed 73 re-sequenced deer genomes from across their North American range to understand the consequences of climatic and anthropogenic pressures on deer demographic and adaptive history. We found a strong signal of glacial-induced vicariance and demographic decline; notably, there is a severe decline in white-tailed deer effective population size (N_e_) at the end of the Last Glacial Maximum. We found robust evidence for colonial overharvest in the form of a recent and dramatic drop in N_e_ in all analysed populations. Historical census size and restocking data show a clear parallel to historical N_e_ estimates, and temporal N_e_/N_c_ ratio shows patterns of conservation concern for mule deer. Signatures of selection highlight genes related to temperature, including a cold receptor previously highlighted in woolly mammoth. We also detected immune-genes that we surmise reflect the changing land-use patterns in North America. Our study provides a detailed picture of anthropogenic and climatic-induced decline in deer diversity, and clues to understanding the conservation concerns of mule deer and the successful demographic recovery of white-tailed deer.

## Introduction

The glacial cycles of the Quaternary with movement of ice sheets, corresponding sea level fluctuations, and global environmental changes, heavily impacted worldwide biota (Batchelor et al. 2019; Allen et al. 2020; Gowan et al. 2021). In the northern hemisphere, species adapted to the temperate climate were forced to adapt in cold periods, move or retreat to refugia, their population decreasing or going extinct in the process; warming episodes allowed for population expansion, colonisations, and adaptive divergence (Pamilo and Savolainen 2004; Lorenzen et al. 2011; Martchenko and Shafer 2023). The peopling of North America ∼15,000 years ago (kya; Willerslev and Meltzer 2021) followed by human expansion was also a driver of regional ecological shifts (Gajewski et al. 2019; Fulton and Yansa 2021; Commerford et al. 2022), although there is no consensus on whether this was responsible for large mammal declines (Stuart 2015; Meltzer 2020; Stewart et al. 2021).

The more recent Anthropocene has dramatically shaped demographic trajectories of wildlife through farming, hunting and logging (Ceballos et al. 2015; IPCC 2022). Human pressures increased during colonial eras when settlers profoundly altered the land, leading to declines in endemic populations (Woinarski et al. 2015; Lindo et al. 2016; Tavares et al. 2019; Brain and Prosser 2022; Stevens et al. 2022). Indeed, colonial impact has been associated with the extinction of several species such as Australian rodents, and the blue antelope (*Hippotragus leucophaeus;* Roycroft et al. 2021; Hempel et al. 2022). Beyond the demographic impacts, settler populations might also have altered selection pressures, with recent examples of selective harvest including brown bears (*Ursus arctos*) and moose (*Alces alces*; Allendorf and Hard 2009; Kvalnes et al. 2016; Van De Walle et al. 2018).

Severe population contractions, or bottlenecks, increase genetic drift and inbreeding risk, and while highly deleterious mutations can be purged, mildly deleterious mutations tend to accumulate (Frankham 2005; Charlesworth 2009), as seen in alpine ibex (*Capra ibex*) and woolly mammoth (*Mammuthus primigenius*; Palkopoulou et al. 2015; Grossen et al. 2020). Population size fluctuations also leave discernible genome-wide patterns: a declining population will exhibit a positive Tajima’s
 D and a relatively flat site frequency spectrum (SFS), whereas an expanding population will present a negative Tajima’s D and a SFS skewed towards low-frequency or rare alleles (Hahn 2018). Depending on the extent of the contraction, both in magnitude and duration, populations might also present with a low effective population size (N_e_), particularly if the bottleneck was strong and the recovery long (Martchenko and Shafer 2023). In contrast, a population that remained steady over time, or one that recovered quickly from a bottleneck, will usually exhibit a larger N_e_ (Charlesworth 2009). Genomic data can be used to model changes in historical and, due to recent analytical advances, contemporary trends of N_e_ (Li and Durbin 2011; Schiffels and Durbin 2014; Santiago et al. 2020; Schiffels and Wang 2020), and when compared to census size (N_c_), both historical and contemporary estimates of N_e_/N_c_ reflect adaptive potential and extinction risk (Palstra and Ruzzante 2008; Wilder et al. 2023).

The species of the North American genus *Odocoileus* (deer) were heavily impacted by the glacial cycles of the Pleistocene and European colonisation. Divergence estimates suggest the speciation between white-tailed (*O. virginianus*; WTD) and mule deer (*O. hemionus*; MD) took place between 500 kya and 4.3 million years (Mya; <u>Baccus et al. 1983; Kessler et al. 2023)</u>, the genus became among the most successful Cervids during glacial cycles (Hewitt 2011). Both species survived in different refugia during the Last Glacial Maximum (LGM, 26 - 19 kya (Clark et al. 2009); Ellsworth et al. 1994; Greenslade 1998; Latch et al. 2009; Latch et al. 2014) and share mitochondrial haplotypes due to incomplete lineage sorting (Klicka et al. 2023). While WTD & MD hybridise (eg. Carr and Hughes 1993; Russell et al. 2021), the hybridisation appears to be a result of secondary contact and did not impact speciation (Kessler et al. 2023).

Deer abundance is heavily monetised through hunting-related activities in North America (Cambronne 2013), and both species have been, and continue to be, important to indigenous communities as a source of food, clothing and culture (Adams and Hamilton 2011; Peres and Altman 2018). In the late Holocene, WTD were likely the most abundant large-mammal prey in eastern North America, and while some Indigenous Peoples depended on them for subsistence, harvest appeared to minimally impact WTD numbers (Wolverton et al. 2008; Weitzel 2021). In contrast, over-hunting and forestry practices post European-colonisation resulted in heavily depleted and even extirpated WTD populations, leading to strict hunting regulations followed by restocking efforts across the USA in the 20^th^ century (McDonald et al. 2004), with stocking efforts leaving discernible admixture patterns in WTD genome (Chafin et al. 2021). Mule deer abundance is, and was, lower than WTD; their populations also declined post-colonisation and plunged in the early 1900’s (Jensen et al. 2023). Stocking efforts were less pervasive for MD, and the species stabilised due to effective management strategies (Gruell 1986; Clements and Young 1997; Gill 1999; Bergman et al. 2015; Jensen et al. 2023). Previous historical demographic analysis for both species have generated both contrasting and coarse demographic reconstructions (Combe et al. 2021; Lamb et al. 2021).

Here, we aim to understand the consequences of climatic and anthropogenic pressures on WTD & MD populations through explicit demographic modelling and selection analyses. We hypothesise that the population genetic histories of North American deer were heavily impacted by the glacial cycles of the Pleistocene and by European colonisation, not the initial peopling of North America. Specifically, we predict that the LGM had a strong negative impact in the form of demographic contraction and population subdivision, and that the impact of European colonisation, notably overhunting and change in land use, would leave detectable signatures on the genome in a neutral and adaptive context. To test these predictions, we used whole genome sequence data of MD and WTD from across their North American range to compute summary statistics, reconstruct and model demographic histories, and quantify genomic variation linked to environmental factors and selective forces.

## Materials and Methods

### Sampling & Sequencing

We sequenced the genomes of 45 deer collected across North America (Fig. 1A, Table S1) and added WGS data from 28 published deer genomes (PRJNA830519; Kessler et al., 2023). The complete dataset comprised 20 MD and 53 WTD samples, including four WTD individuals from the Florida Keys subspecies (*O. v. clavium*). We extracted DNA using the Qiagen DNeasy Blood and Tissue Kit following manufacturer’s instructions and checked concentration using Invitrogen Qubit assays. DNA was sent to The Centre for Applied Genomics in Toronto, Canada, for library preparation following the Illumina TruSeq PCR-free DNA Library Prep, and sequencing to generate 150bp paired end reads. All samples were sequenced to an average of 4x coverage on an Illumina HiSeqX, except for four samples which were re-sequenced to achieve higher coverage (>15x; Table S1, Fig. 1A). This 4x coverage was targeted as we generated genotype likelihoods that are designed for lower coverage data (Meisner and Albrechtsen 2018).

**Figure 1:**
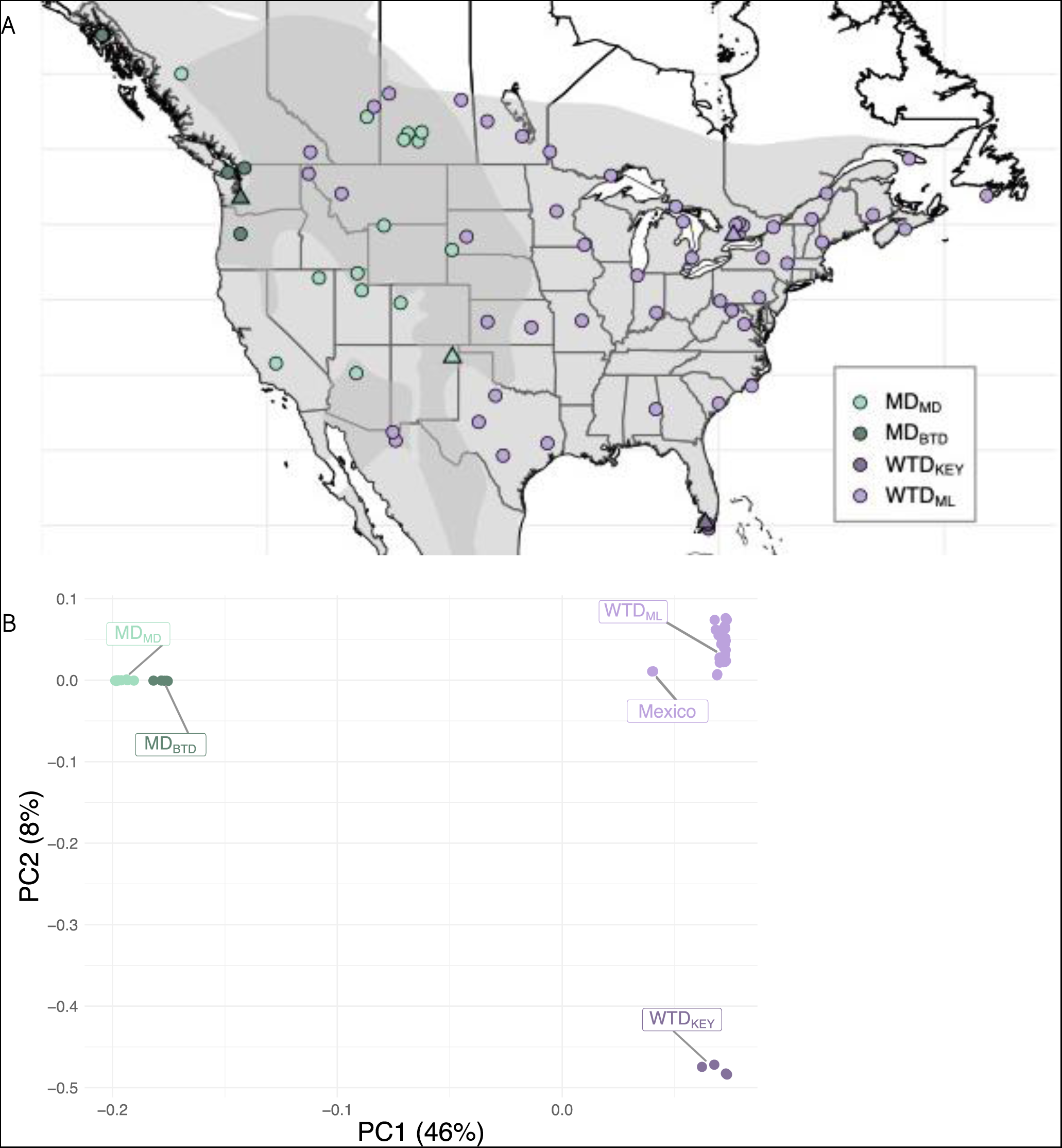
Sample information. (A) Sampling locations and population assignment, triangles represent high-coverage samples, shaded areas represent each species’ IUCN range with areas of sympatry in a daker shade (B) PCA of all individuals, based on allele frequencies. Colour coded by assigned population.

We obtained estimated census data for WTD and MD in the USA for the last 400 years from Webb (2018; complemented by Deer Friendly). This census data was generated using a process of environmental scanning that aggregated data from different sources such as state agencies, harvest records and historical sources for the most ancient timepoints. We also digitised restocking information for WTD in the USA from McDonald et al. (2004).

### Data processing

We analysed and trimmed raw reads with FastQC (v0.11.9; Andrews 2010) and Trimmomatic (v0.36; Bolger et al. 2014), respectively. Using bwa-mem and default settings (v0.7.17; Li and Durbin 2009), we aligned raw reads to the WTD genome (GCA_014726795.1) and sorted the reads using SAMtools sort (v1.10; Li et al. 2009). We identified and removed duplicates in Picard MarkDuplicates (v2.23.2; Broad Institute, GitHub Repository. 2019) and Sambamba view (v0.7.0; Tarasov et al. 2015). We used GATK RealignerTargetCreator and IndelRealigner (v4.1.7.0; McKenna et al. 2010) to carry out a local re-alignment and used Sambamba flagstat, mosdepth (v0.3.1; Pedersen and Quinlan 2018) and MultiQC (v1.10; Ewels et al. 2016) for quality checks.

For samples sequenced on multiple lanes, we specified the read groups before removing duplicates and merged the sequencing files using Picard AddOrReplaceReadGroups and MergeSamFiles, respectively. To have a consistent coverage among samples for non-SMC demographic analyses, we down-sampled data to the average coverage of all other samples (4x) using Picard DownsampleSam. We called the genotypes (-doGeno 4) and estimated genotype likelihood using the GATK model (-gl 2) in ANGSD (v0.918; Korneliussen et al. 2014). We retained SNPs with a minimum base quality and a minimum mapQ quality of 20 (- minQ 20 & minMapQ 20) and a minimum p-value of 1e-6 (-SNP_pval 1e-6).

### Population subdivisions and summary statistics

We used PCAngsd (v1.02; Meisner and Albrechtsen 2018) to investigate population clustering between and within species, this program fits best our data as it is designed explicitly for low-coverage data and genotype likelihoods. We explored the relationship between the spread on the PCA and the geographic distribution of our samples using a Pearson’s correlation test in R (v4.2.0; R Core Team 2021). We further generated an individual genetic distance matrix using the gene.dist() function from the ape R package (Paradis and Schliep 2019), with the dataset produced for the δaδi analysis (see below). Finally, to help decipher population structure in the MD samples, we further ran NGSadmix (Skotte et al. 2013), using K = 1 : 10 and 10 replicates, we identified the best K value using CLUMPAK (Evanno et al. 2005; Kopelman et al. 2015). We introduce the following convention to refer to populations supported by PCAngsd and NGSadmix and used in subsequent analyses: mainland WTD = WTD_ML_, Florida Keys WTD = WTD_KEY_, MD subspecies = MD_MD_, and MD black-tailed deer subspecies = MD_BTD_.

We computed the genome-wide Tajima’s D for all populations following the ANGSD pipeline (http://popgen.dk/angsd/index.php/Thetas,Tajima,Neutrality_tests). Briefly, we produced a folded SFS with angsd -doSaf 1 and realSFS, computed theta with realSFS saf2theta followed by Tajima’s D using thetaStat. Further, we converted the hardcalls generated in ANGSD (.geno file) into a .vcf file using a python script (genoToVCF.py; Martin 2021) and retained only biallelic sites (--max-alleles 2) in VCFtools (v0.1.16; Danecek et al. 2011). Using VCFtools we computed weighted F_st_ between the four populations, and nucleotide diversity (π) per population, all processed in windows of 50kbp. We used PLINK (v1.90b6.21; Chang et al. 2015) to estimate the inbreeding coefficient based on runs of homozygosity (F_ROH_) on the 160 scaffolds representing the N90 of the WTD’s genome assembly, using the following non-default criteria: 100kb window size containing a minimum of 50 SNPs, allowing up to three heterozygote sites and 10 missing calls per called ROH. We computed F_ROH_ per individual using the sum of ROH sizes divided by the sum of N90 scaffolds length.

### Historical and recent demographic inference

We inferred the historical demography of WTD_KEY_, WTD_ML_, MD_MD_ and MD_BTD_ by implementing the multiple sequentially Markovian coalescent (MSMC) model (Schiffels and Durbin 2014) that generates estimates of N_e_ over time. Here, we used four high-coverage samples, one per population (Fig. 1A, Table S1), and first generated a 35-mer mappability mask file using SNPable, as required for MSMC with several samples (Schiffels and Wang 2020). Because of the small sample size of high coverage genomes, we phased the data with WhatsHap (v1.3; Patterson et al. 2015; Martin et al. 2016) and filtered out scaffolds shorter than 500 Kbp. We ran MSMC2, including 100 bootstraps, using 1×2+25×1+1×2+1×3 as time segment pattern. To estimate the cross-coalescence rate (CCR) across populations, we carried out a pairwise comparison of the first haplotype from each sample using the -P flag, skipping any site with ambiguous phasing. Finally, to test and account for ancestral migration, we implemented MSMC-IM (Wang et al. 2020), assuming a generation time of 2 years (Deyoung et al. 2003) and a mutation rate of 1.23x10^-8^ mutations/site/generation (average from Table S16 in Chen et al. 2019). We complemented this inference with a stairway plot analysis (v2.1.2; Liu and Fu 2015; Liu and Fu 2020) to reconstruct historical population size changes. Using each population, we generated a folded site frequency spectra in easySFS (v0.0.1; Gutenkunst et al. 2009) and used the resulting SFS as input to stairway plot, along with the same mutation rate and generation time as used for MSMC2.

We used GONE (Santiago et al. 2020) to compute demographic history in the recent past from patterns of linkage disequilibrium (LD) for the four populations. We restricted this analysis to the 160 scaffolds representing the N90 of the WTD assembly. We ran GONE with 500 repetitions on 500 generations with 30,000 SNPs per chromosome, a minor allele frequency pruning of 0.01, a hc value of 0.04, a recombination rate of 0.5 for WTD populations and 0.35 for the MD populations (Kessler et al. 2023). We compared the census size (N_c_) point estimates from Webb (2018) to the temporal estimates of N_e_ resulting from GONE to obtain a contemporary N_e_/N_c_ ratio over time.

We further filtered the dataset in VCFtools to I) account for LD by removing sites that were within 10kbp of each other (--thin 10000), and II) remove all missing data (--max-missing 1). To model explicitly the demographic history of WTD and MD, we used δaδi (v 2.1.1; Gutenkunst et al. 2009) and dadi_pipeline (v3.1.7; Portik et al. 2017) and performed two different demographic analyses with different outgroups: 1) WTD_KEY_, WTD_ML_ and all MD, and 2) MD_MD_, MD_BTD_ and WTD_ML_. We selected relevant demographic models involving migration and asynchronous splitting (Table S2), optimised and fitted the models with four repetitions and default settings, the best model was selected based on the log-likelihood and its consistency across runs. This model was additionally run with 100 bootstrapped frequency spectrums to quantify uncertainty. We obtained ancestral N_e_/N_c_ ratio over time for both species using the ancestral N_e_ (NuA) from the δaδi analyses.

### Adaptive divergence and selection scans

We used a redundancy analysis (RDA) to detect genotype-environment association and loci under local adaptation across individuals, RDA is a multivariate regression method that allows to identify loci associated with a variety of predictors (Legendre and Legendre 2012). In our case, the predictors are 19 bioclimatic variables (Table S3) which we collected at a resolution of 2.5’ from WorldClim2 using the R package raster (v.3.6-3; Hijmans 2022; Fick and Hijmans 2017) and extracted the values corresponding to our samples’ coordinates. We performed three analyses: one inter-specific with all individuals, and two intra-specific for WTD & MD using 19 bioclimatic variables, latitude, longitude, and species as predictors. As highly correlated variables are problematic in RDAs, we removed those that showed a Pearson correlation > 0.7 in each analysis (Dormann et al. 2013). We carried out the RDA with the R package vegan (v.2.6-4; Oksanen et al. 2022) based on the datasets created for demographic modelling. In addition to full models including non-correlated predictors, we investigated conditional models with the condition set as either the bioclimatic variables or longitude.

To detect regions under selection in the four populations, we computed the integrated haplotype score (iHS) on the scaffolds representing the N90 of the WTD reference genome. This statistic is based on extended haplotype decay to detect selective sweeps by comparing the state of decay between ancestral and derived allele (Sabeti et al. 2002; Voight et al. 2006). The state of decay for all alleles should be similar in neutral regions and iHS should equal zero in the absence of selection. In the case of a selective sweep and as one allele will rise in frequency, haplotype decay will be slower and iHS values will become either positive or negative depending if the sweep takes place on the ancestral or derived allele, respectively (Voight et al. 2006). As this analysis included all samples, we started by phasing our data in SHAPEIT (v2.r904; O’Connell et al. 2014), then calculated and normalised iHS in vcflib (v1.0.3; Garrison et al. 2022) with the functions iHS and normalize-iHS and default settings. We identified peaks in the most extreme 1% iHS values in R using the function findPeaks (Grinberg 2019) and extracted gene IDs within 25Kbp upstream and downstream. The genes with available entry IDs were uploaded to Uniprot (Pundir et al. 2016) for ID mapping where we retrieved the gene names and functions. The gene list was manually curated, and each gene was assigned to one or more of the following categories based on its function: reproduction, immunity, physiology, metabolism, sensory perception, development, and other. We analysed all results and produced all the graphics in R.

## Results

### Population subdivisions and summary statistics

We called 144,634,190 SNPs across species from 73 individuals at a coverage of 4x on average. The PCA on allele frequencies of all individuals presents a clear species delimitation along PC1 which explained 46% of the variation, whereas PC2 separated the WTD_KEY_ individuals (Fig. 1B). WTD-specific PCA showed the delimitation between WTD_ML_ and WTD_KEY_ individuals (PC1); PC2 strongly correlated with longitude (Pearson correlation = -0.88, p-value = < 2.2e-16) and accounted for 7% of the variation (Fig. S1A). In the MD PCA, PC1 correlated to longitude (Pearson correlation = 0.83, p-value = 5.96e-06), and PC2 associated to latitude (Pearson correlation = 0.47, p-value = 0.037, Fig. S1B). The MD PCA showed one clear cluster of 13 samples; we clarified the population identity of the seven remaining individuals using NGSadmix that assigned two admixed individuals to the first cluster (MD_MD_) and identified a second population of five individuals (MD_BTD_; Fig. S1C). Individual genetic distances separated MD and WTD_KEY_ individuals from WTD (Fig. S1D). The individual genetic distance mirrored nucleotide diversity estimates which were highest in WTD_ML_ and similar in the other three populations (Table 1, Fig. S1D).

**Table 1:** Tajima’s D, nucleotide diversity and F_ROH_ for our four populations, F_ROH_ as percentage of the N90 scaffolds.

|  | **Tajima's D** | **Nucleotide**<br>**diversity ($\pi$)** | **$F_{ROH}$** |
| --- | --- | --- | --- |
| **WTD<sub>ML</sub>** | -0.76 | 0.0068 | 0.89 |
| **WTD<sub>KEY</sub>** | 0.70 | 0.0038 | 1.38 |
| **MD<sub>MD</sub>** | 0.81 | 0.0036 | 0.30 |
| **MD<sub>BTd</sub>** | 1.08 | 0.0037 | 0.28 |

Weighted F_st_ measures were high in between species comparisons (0.36 - 0.58); intra- specific comparisons ranged from 0.12 to 0.17 (Table S4). Tajima’s D values were negative only for the WTD_ML_ population (−0.76), the highest being 1.08 for MD_BTD_ (Table 1,Fig. S2A). F_ROH_ values were lowest in the MD populations (<= 0.3%), whereas WTD_KEY_ presented the highest F_ROH_ value (1.4%; Table 1; Fig. S2B).

### Demographic inferences

We inferred the demography of four high-coverage samples, one per population, using MSMC2. The trajectories of the four samples split at approximately 600 kya where the MD_MD_ population start declining whereas WTD populations keep increasing (Fig. 2A). Both MD population trajectories are very similar and show a general steady decline. WTD_KEY_ appear to decline ∼50 kya, whereas WTD_ML_ drastically increased to peak at an N_e_ of >1 million individuals, followed by a clear crash at the end of the LGM (Fig. 2A). Species split times inferred by the cross-coalescence rate of the MSMC-IM model is approximately 500 kya (Fig. S3A), the within-species split times are estimated at ∼ 25 kya between WTD_ML_ and WTD_KEY_, and ∼70 kya for MD_MD_ and MD_BTD_ (Fig. S3B). Effective population size over time as inferred by the stairway plot analysis showed similar historic trajectories where all populations have declines surrounding the LGM (Fig. S4). However, both MD_BTD_ and WTD_ML_ presented a large recovery post-LGM followed by a recent crash; this is consistent with model overfitting in a stairway plot that produces spurious and complex patterns (Lapierre et al. 2017).

**Figure 2:**
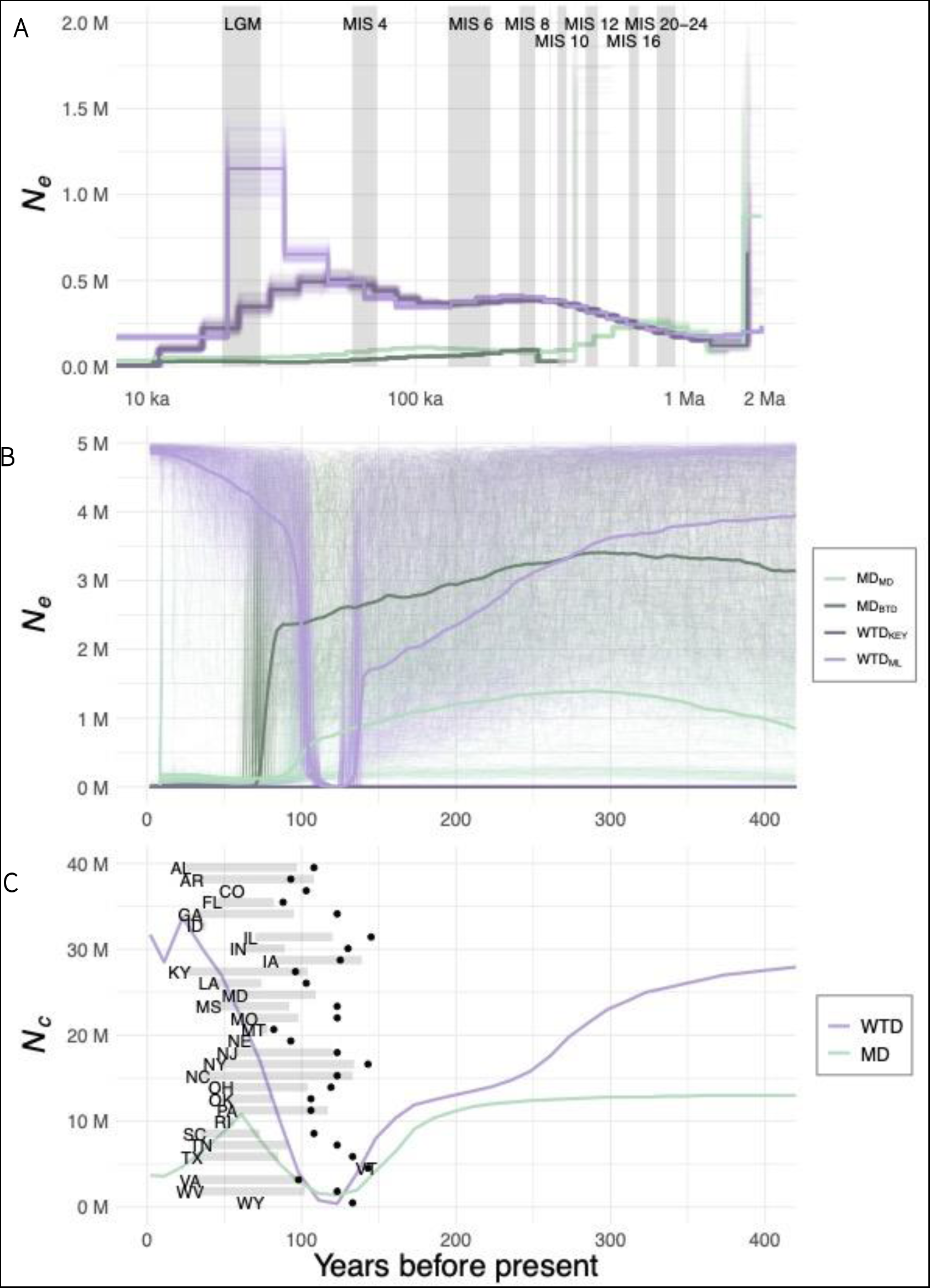
Demographic inference. (A) Changes in population size of four high coverage samples with bootstraps as estimated by MSMC2, shaded areas symbolise full glacial periods (Batchelor et al. 2019). (B) Recent changes in N_e_ computed by GONE on all samples with bootstraps. (C) Estimated WTD & MD population in the US from Webb (2018) with WTD restocking period for 30 US states (grey rectangles) and corresponding state’s year with lowest WTD population (black points) from McDonald et al. (2004).

The N_e_ of WTD_KEY_ appears very low and stable for the last ∼400 years, reaching its lowest at ∼350 individuals (Fig. 2B). Temporally, the WTD_ML_ population decreased with variable intensity that ended with a drastic decline to a N_e_ of ∼2000 near the end of the 19^th^ century. Following this severe drop, the WTD_ML_ population rapidly rebounded to reach a current effective size in the millions (Fig. 2B); this clearly correlates with census data and restocking efforts (Fig. 2C). Both MD populations exhibited a slow but steady increase until approximately 300 years ago, after which both trajectories show a gradual decline ending with a severe drop without recovery (Fig. 2B). MD N_e_ patterns are more decoupled from the census data as N_c_ here shows a crash in both MD_MD_ and MD_BTD_ followed by a small recovery (Fig. 2C). As such, the contemporary N_e_/N_c_ ratio is higher and more variable for MD (Fig. 3; mean = 0.49, var = 0.3 vs WTD mean = 0.18 & var = 0.01) with most values from the last century below 0.1. Both species have a low ancestral N_e_/N_c_ ratio with low variability, the averages are of 0.06 and 0.01 for WTD and MD, respectively. White tailed deer’s historical ratio (var = 0.03) is more variable than MD (var = 0.0001) and the contemporary WTD (Fig. 3).

**Figure 3:**
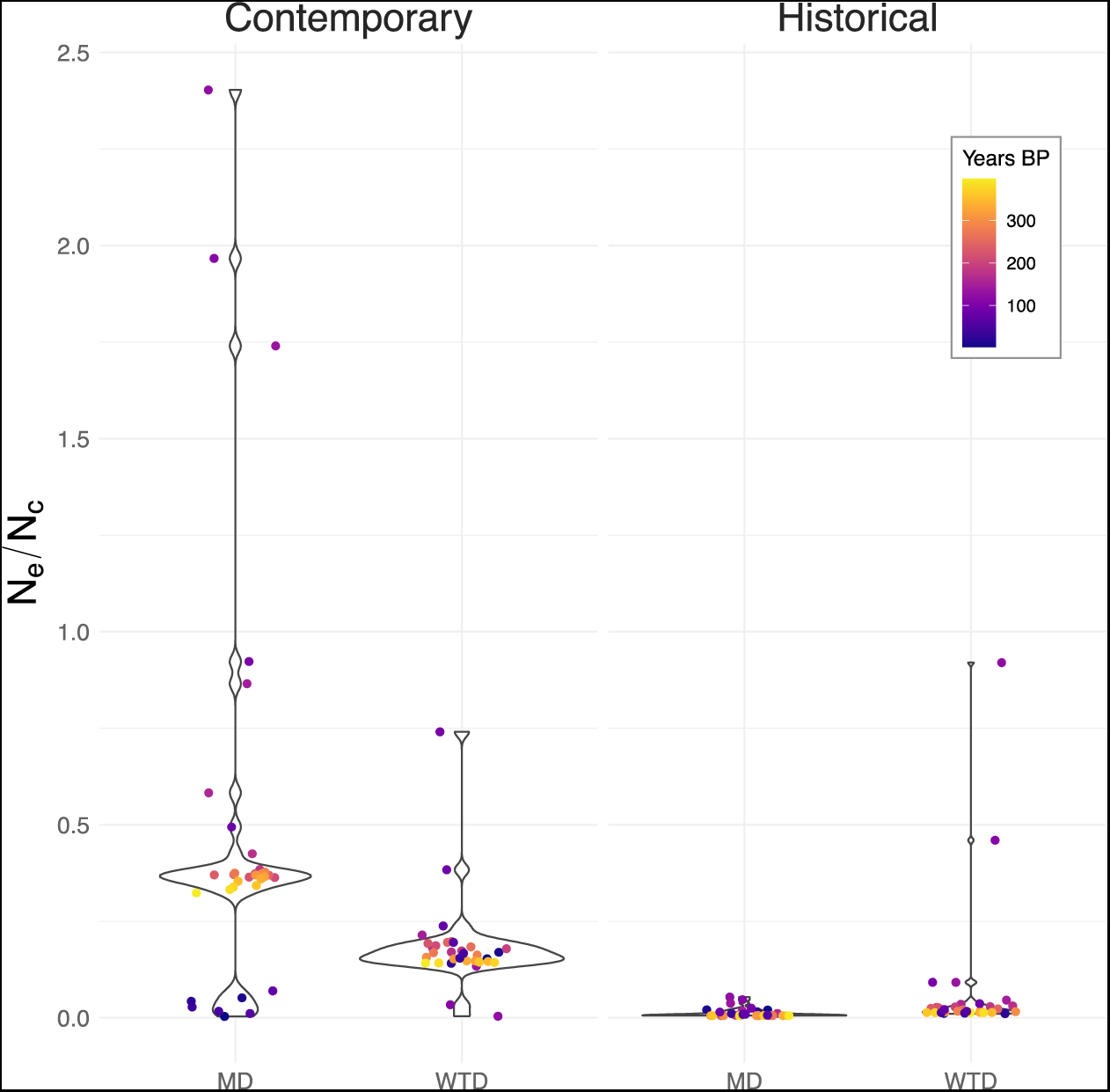
Contemporary and historical N_e_/N_c_ ratio over time, point colour indicates time before present, N_e_ of the historical ratio taken from NuA values in the δaδi analysis.

Finally, we modelled the demographic history of our four populations using the site frequency spectra. For our first comparison, we analysed the demographic history of the trio WTD_KEY_ – WTD_ML_ – MD, the best model supports adjacent secondary contact between populations, however migration rates remain low (m < 1e^-5^; Fig. 4A, Table S5). This model suggests a MD/WTD split ∼280 kya followed by a MD decline and a WTD expansion. After the split between WTD_KEY_ and WTD_ML,_ both populations declined with notably a demographic crash in WTD_KEY_; all of these predate the peopling of North America. In our second trio, we investigated the relationship between MD_MD_ – MD_BTD_ – WTD_ML_. In this case, the split WTD/MD is seen ∼390 kya, the best model suggests a secondary contact between WTD_ML_ – MD_MD_ that predates the split between the two MD populations, ∼13 kya, but all migration rates are extremely low (m <= 4e^-6^). This model also suggests population expansion for WTD_ML_ after the split from MD, but a decline for both MD_BTD_ and MD_MD_ before and after the peopling of North America (Fig. 4B, Table S5).

**Figure 4:**
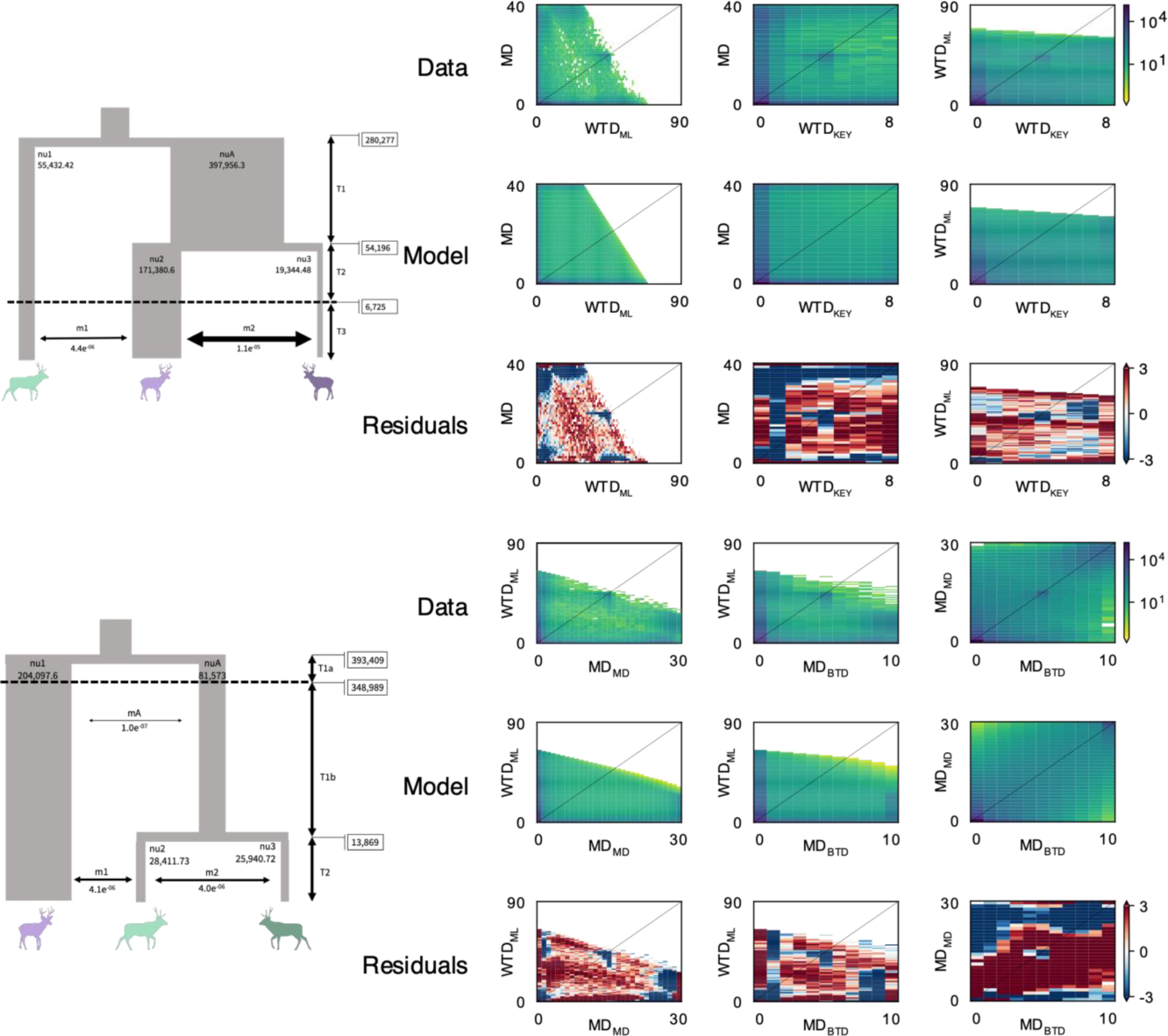
Three populations models in δaδI with best model schema on the left and 2D spectra on the right. (A) MD – WTD_ML_ – WTD_KEY_ comparisons, with MD in green, WTD_ML_ in light and WTD_KEY_ in dark purple. (B) WTD_ML_ – MD_MD_ – MD_BTD_ comparison with WTD_ML_ in light purple, MD_MD_ in light green and MD_BTD_ in dark green. Population sizes (nu) and migration rates (m) estimates to scale, time is not to scale.

### Adaptive divergence and selection

We conducted three redundancy analyses to identify genotypes linked to environmental variables and loci under local adaptation. The RDA with all individuals explained 16.5% of the variation, most of which was driven by the species variable (Table 2, Fig. S7A). The full model for the WTD explained 6.32% of the variation and included five significant predictors - BIO1 (annual mean temperature), BIO2 (mean diurnal range), BIO8 (mean temperature of wettest quarter), BIO15 (precipitation seasonality) and longitude (Table 2, Fig. S7B). There were 12,206 outlier SNPs, predominantly associated with annual mean temperature (Table 2 & S3, Fig. S7C). In the RDA on MD, we found only BIO2 and BIO16 (precipitation of wettest quarter) as significant predictors, and 7434 outlier SNPs, mostly associated with BIO16 (Table 2). Conditional models of bioclimatic variable explained more variation than those of longitude in both species (Table 2).

**Table 2:**
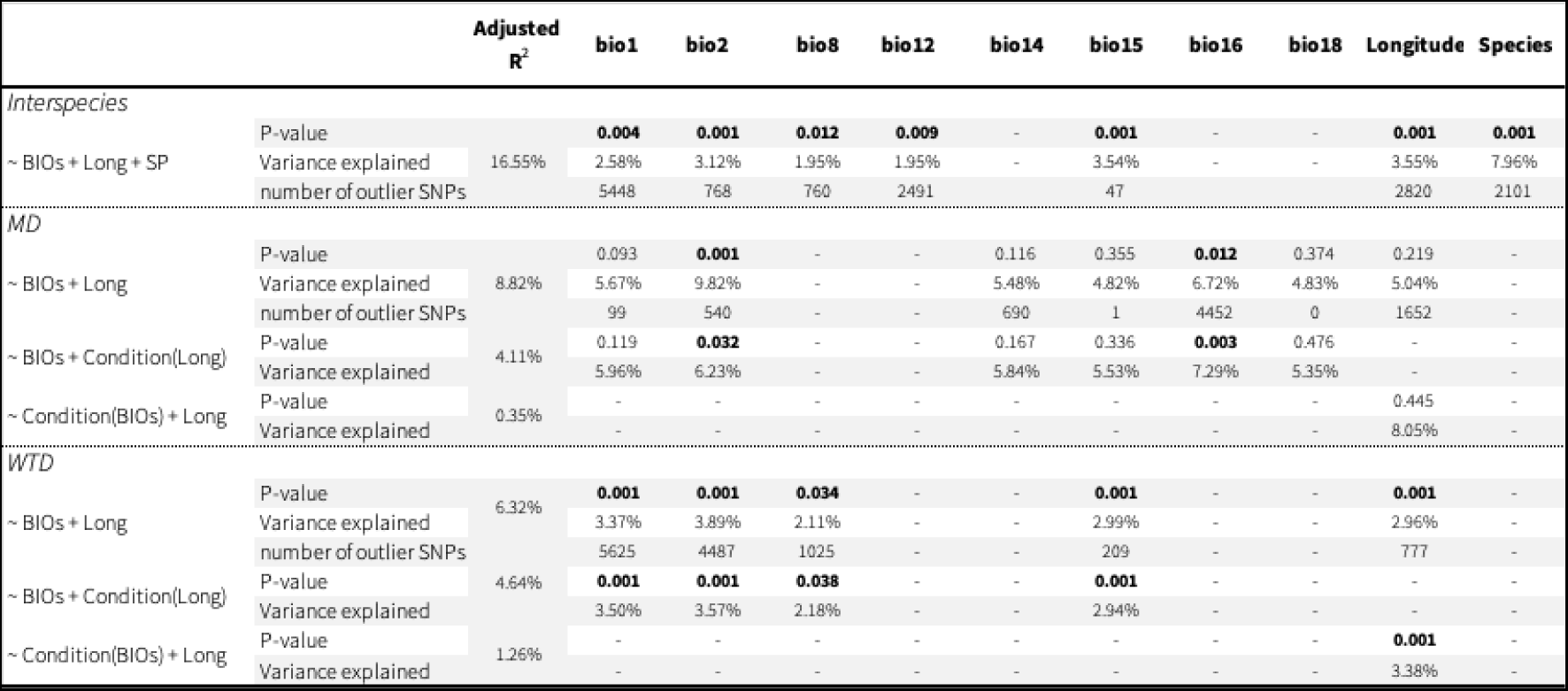
Anova of the RDA for full and conditional models as well as number of outlier SNPs for each predictor in the full model. Significant p-values are highlighted in bold.

We found a total of 443 genes within 25 Kbp upstream and downstream of an iHS peaks, including 249 with an existing Entry ID (Table S6). There were 121 genes around iHS peaks in the WTD_ML_ population, 45 of which we could assign an explicit category including 18 genes assigned to immunity, of which two related to the major histocompatibility complex (MHC): *HLA-DQA1* and *HLA-DQB*. Additionally, we found two genes linked to sensory perception: one odorant receptor (*OR5D18*) and one cold receptor (*TRPM8*, Table S6). We identified 23 genes surrounding iHS peaks in the WTD_KEY_, including one MHC gene: *HLA- DRB4*. For the MD_MD_ population, we could explicitly categorise 34 genes on a total of 95 neighbouring iHS peaks. They include four MHC genes: *HLA-DQA1*, *HLA-DQB1*, *HLA-DRB3* and *HLA-DRB4,* another gene related to the MHC: *HIVEP2*, three genes described as olfactory or odorant receptors (*OR51I1*, *OR5D18*, *TAAR5*), and one cold receptor (*GRIK2*). Finally, we identified 26 genes adjacent to iHS peaks in the MD_BTD_ population, including two related to immunity and three linked to reproduction.

## Discussion

The complex climatic cycles of the Pleistocene, concluding with the LGM, generally forced species to move from a wide range into smaller, often isolated refugia (Soltis et al. 2006; Shafer et al. 2010). The peopling of North America and the colonisation by Europeans several millennia later represented further pressures to many populations inhabiting the continent (McDonald et al. 2004; Lindo et al. 2016; Smith et al. 2021). Here, we reconstructed the demographic history and signals of selection of WTD & MD to improve our understanding of climatic and anthropogenic pressures on these two North American species. Through extensive genomic analysis we observed a remarkable overlap between climate, recent human drivers, and N_e_, and provide a temporal N_e_/N_c_ assessment supporting the demographic concerns facing MD (Bergman et al. 2015; Webb 2018). Further, our selection scans that reflect climate and disease pressures provide testable hypotheses for the impact of European colonisation.

### Strong historical climatic impact on deer populations

The glacial cycles of the Quaternary forced populations to move or adapt, resulting in population divergence (Colella et al. 2021; Ito et al. 2021), decline (Ersmark et al. 2019; Baca 2020; Dussex et al. 2020; Meiri et al. 2020) and even replacement (Loog et al. 2020). Our results suggest a clear impact of climate on WTD during the late Quaternary as we observed an intense N_e_ drop for WTD_ML_ as well as a population split time that coincides with the LGM (Fig. 2A, Fig. S3A), and support WTD populations being separated into different refugia during this period (Ellsworth et al. 1994; Moscarella et al. 2003; Combe et al. 2021; Wright et al. 2022). The Key deer showed a steady decline that predated the LGM, with a clear separation from WTD_ML_ ∼50 kya (Fig. 2A, Fig. 4A, Table S5). All analyses suggest both populations started diverging before the isolation of the Florida Keys from the continent ∼8,000 years ago (Ellsworth et al. 1994; Villanova et al. 2017). This isolation led to a clear loss of genomic diversity (Table 1, Fig. S2) and accumulation of deleterious alleles in the endangered Key deer (Cars et al. 2023; U.S. Fish & Wildlife Service 1967). Overall, the collective demographic methods used here pinpoint the LGM as a driver of subdivision and historical declines in WTD. The LGM has also been implicated in the MD and BTD (*O. h. columbianus*) subdivision, as the two subspecies were separated in different refugia during that time (Latch et al. 2009; Latch et al. 2014; Wright et al. 2022). Our results are equivocal regarding the split time: the MSMC analysis suggest a steady decline of both populations and a MD/BTD split estimated at approximately 70 kya by the CCR (Fig. 2A, S2B); δaδi analysis, in contrast, places the split after the LGM and concomitant with a decline of ∼30% in both populations (Fig. 4B, Table S5). While we present here the best model for our data, both comparisons still present with relatively large residuals (Fig. 4) suggesting the true story might be more complicated than depicted here. Additional modelling with different approaches might help render a more precise history. Nevertheless, both analyses support changes pre-peopling of North America and are consistent pre-split: the N_e_ of the common ancestor of MD_MD_ and MD_BTD_ appeared low but stable through time (Fig. 2A). As MD are more prone to population fluctuations (Forrester and Wittmer 2013), and N_e_ is most strongly influenced by long periods of low N_c_ (Peart et al. 2020), the overall stable historical N_e_ is consistent with population genetic theory.

Hahn (2018) suggested to focus on big picture, not point estimates, for demographic inferences, and while MSMC reconstructions differ in magnitude based on data treatment, overall time and shape remains constant (Schiffels and Wang 2020). While the general historical patterns of demographic change observed here support previous assessments (Lamb et al. 2021), including the LGM impact in WTD, they are in contradiction with Combe et al. (2021). Combe et al’s (2021) stairway plot analysis of ∼25k SNPs suggests a single population contraction ∼100 kya followed by an expansion for WTD, and population plateau at high number for over 200 kya before a drastic decline for MD. These times are inflated due to their generation time of five years versus two used here <u>(see also Deyoung et al. 2003)</u>. Simulations, have shown that a dataset of this size is cannot reliably resolve changes in N_e_ (Shafer et al. 2015), thus our explicit modelling and genome-wide data set has significantly more power and precision with respect to historical demographic inference. Further stairway plots are prone to over-fitting and erroneous reconstructions (Lapierre et al. 2017). For example, the explosive LGM recovery suggested by our WTD_ML_ stairway plot is at odds with our all other results and patterns seen in comparable species (e.g. <u>Dussex et al. 2020; Taylor et al. 2021)</u>.

None of the deer populations analysed here showed any signal of recovery after the LGM and ice sheets recession. This is particularly striking given the massive loss of large mammals on the continent (Elias and Schreve 2007; Stuart 2015; Meltzer 2020), presumably opening new niches, reducing competition and predation. One explanation could be that the primary cause of population declines in deer were the environmental shifts caused by the deglaciation dynamics (Mottl et al. 2021; Hanberry 2023) and the changes in ecosystem function after the megafaunal extinctions (Malhi et al. 2016; Malhi et al. 2022). The lack of full recovery is consistent with population dynamics of other temperate ungulate species (De Jong et al. 2020; Dussex et al. 2020; Taylor et al. 2021; Robin et al. 2022). Nevertheless, the limits of detection of MSMC2 and GONE (10 kya – 0.4 kya) do leave a gap in the Holocene for which it is difficult to reconstruct demographic history; however, δaδi should pick up a large Holocene change <u>(e.g. Dedato et al. 2022)</u>, or the pattern seen in the stairway plot analysis, were they to have actually occurred.

Hybridisation between WTD and MD in areas of sympatry is well documented (eg. Cronin 1991; Derr 1991; Carr and Hughes 1993), but it was suggested that introgression had no impact on species divergence and rather that gene flow is a result of secondary contact (Kessler et al. 2023). Likewise, <u>Klicka et al. (2023)</u> suggested incomplete lineage sorting of mitochondria, not historical hybridisation, explains shared haplotypes between species. This is consistent with our analyses as both δaδi comparisons suggest a secondary contact with low migration rates (Fig. 4, Table S5), and while some ancient migration is detected, it is extremely low (Fig. 4B, Table S5, Fig. S3C). Our results therefore support the finding of negligible ancient gene flow between species. In fact, our analyses suggest a species split time under 500 kya (Fig. 4, Table S5, Fig. S3), which are more recent than previous molecular clock evaluations ranging between 750 kya and 4.3 mya (Baccus et al. 1983; Douzery and Randi 1997; Combe et al. 2021; Wright et al. 2022). All of these estimates were based either on a restricted number of nuclear and mitochondrial loci, presenting lower power of analysis than our whole genome assessment.

### Human-induced collapse, varying selection pressures on white-tailed deer

Driven by overharvest and change in land use for agriculture and logging, the recent decline and near extirpation of WTD has been thoroughly documented in the USA (McDonald et al. 2004). The reconstructed demographic trajectory of the mainland WTD population mirror these historical sources and census size estimates (Fig. 2B-C); specifically, we capture the gradual decline followed by dramatic collapse at the end of the 19^th^ century. This depletion led authorities to implement protective laws and managers to supplement deer populations with translocations (see McDonald et al. 2004 for an extensive summary). Those measures were responsible for the quick rebound we observed for the WTD_ML_ population (Fig. 2B) as analyses of nearly extirpated populations suggest the WTD bottleneck had minimal impact on diversity thanks to the rapid population expansion following translocations (Deyoung et al. 2003; Budd et al. 2018). Together with the stable relationship of N_e_/N_c_ over time we observed (Fig. 3), these results suggest that WTD are highly resilient; however, despite the recovery and high diversity, assessment of deleterious alleles also showed a high genetic load for WTD (Cars et al. 2023; Wootton et al. 2023).

Given the low N_e_ and stocking efforts of WTD, it is probable that any sweep that took place before the demographic collapse was either lost or fixed, with the latter generally having reduced power in haplotype scans (Szpiech 2021). In contrast to ungulates experiencing reduced human intervention (e.g. Martchenko and Shafer 2023), longitude had a minimal impact in our models which is consistent with stocking efforts largely erasing geographic signals. Combined with evidence showing rapid contemporary evolutionary responses (Zamorano et al. 2023), we suggest that the outliers and sweeps detected in our analyses could be the result of selection during European colonisation that coincided with large changes to the landscape. In mainland WTD, we found a variety of immunity-related genes potentially under selection which would suggest selection pressures from pathogens (Table S6), some of which might have arisen recently from livestock (Campbell and VerCauteren 2011). The presence of selection at immune genes in WTD scans is consistent with the development of agriculture and large land use changes that have transpired in North America over the past two centuries. Nevertheless, as immune genes are often detected in genome scans and as hyper-variable regions of the genome such as the MHC are susceptible to misalignments (Manel et al. 2016), these results should be taken with caution. Recent selection at immune loci, and indeed the colonial impact, could be investigated by using historical samples: here, any sweep signal detected in the present analysis not seen in historical or ancient DNA would suggest a colonial selection event.

We also found selection patterns at the gene *TRPM8* conferring sensitivity to cold temperature (Table S6); mutations identified in woolly mammoth’s *TRPM8* were suggested to be linked to the species’ adaptation to cold climates (Lynch et al. 2015). Interestingly, in the RDA between species, as well as in the WTD RDA, most outlier SNPs are associated with mean annual temperature (BIO1) rather than species or longitude meaning that temperature is a major driver of differentiation. WTD are exposed to a wide range of temperatures on the continent, but also rapidly expanding northward (Dawe and Boutin 2016). As climate change clearly impacted historical and contemporary WTD ranges (Dawe and Boutin 2016), outliers and selection on genes like *TRPM8* are expected, and likely partly responsible for the success of WTD on the continent. Only few genes were identified as under selection for MD_BTD_ and WTD_KEY_, given the fewer number of samples from those populations the power of the selection scans analysis was clearly reduced (Klassmann and Gautier 2022).

### Delayed collapse but similar selection pressures on mule deer

Human Settlements on the MD historic range were estimated at less than 20 in 1800, and while harvest likely depressed populations in the surrounding areas, the rest of the range probably saw minimal impact (Jensen et al. 2023). Over time and with the increased settlements in western North America, MD populations strongly declined to reach their lowest at the end of the 19^th^ century (Gill 1999; Bergman et al. 2015; Jensen et al. 2023). This is consistent with our analysis which shows a sharp decrease in the MD_MD_ population during that period, though the MD_BTD_ population decline appeared later (Fig. 2B). These population reductions also appeared to happen after the WTD collapse, possibly reflecting the later western wave of settlers and more recent land use changes. As a migratory species, MD might respond more to human-induced habitat change, for example, the limitation in winter range habitat appears to be driving MD decline in Colorado (Gill 1999; Bergman et al. 2015). Collectively the delayed decline, the decoupling of N_e_ from N_c_ (Fig. 3), the positive Tajima’s D and the more muted temporal response of N_e_ are consistent with the species ecology and human intervention history. Despite this collapse and lack of recovery, both MD subspecies exhibit relatively high genetic diversity and low F_ROH_ (Table 1, Fig. S2B), all promising signals for a species that is often described as declining. Assessment of MD genetic load should help further inform the management challenges facing this species.

In MD_MD_, we found 14 immunity-related genes under selection (Table S6). Diseases affecting MD are often the same that infect WTD, including the risk of spilling over to and from livestock (Campbell and VerCauteren 2011). Chronic wasting disease is of particular concern for MD as it is considered as a major factor in the species decline in many regions (DeVivo et al. 2017), with the genomic and immune response clearly polygenic (Seabury et al. 2020). Regarding sensory perception, we found selection patterns at the cold receptor *GRIK2*. Cold adaptation might have been a determinant factor for MD as they favour higher elevation habitats (Anthony and Smith 1977; Brunjes et al. 2006), and are more tolerant to cold climates (Mautz et al. 1985). Supporting this was outliers associated with mean diurnal range (BIO2), which is linked to temperature (Table S63). Most outlier SNPs were associated with precipitation of the wettest quarter (BIO16, Table 2), likely reflective of MD_BTD_ inhabitation of temperate rainforests (Heffelfinger and Krausman 2023).

### Variable N_e_/N_c_ ratio through time

The N_e_/N_c_ ratio is informative on the genomic health and demographic histories of species, which is why it has often been used to assess populations of management and conservation concern (Frankham 1995; Palstra and Ruzzante 2008; Ferchaud et al. 2016). Indeed, a population exhibiting a low contemporary N_e_/N_c_ ratio is more subject to rapid loss in genetic diversity than populations with a higher ratio (Frankham 1995; Palstra and Ruzzante 2008). While, this ratio is highly variable between taxa, the generally accepted median of concern in wild populations is 0.1 (Frankham 1995; Waples et al. 2013). Our unique integrated approach allowed us to measure this ratio over time, with the seven most recent values for MD all below 0.1 (Fig. 3). The larger fluctuations in MD’s N_c_ in the last 100 years appear to have led to a decoupling of N_e_ and N_c_ and reflect the species higher demographic stochasticity. This recent reduced ratio could mean that the species is at risk of faster loss of genetic diversity and echoes other calls of concern (Gill 1999; Bergman et al. 2015). Interestingly, historical N_e_/N_c_ ratio presents low variability for both species and no immediate red flag (Wilder et al. 2023); this ratio is clearly driven by the high N_c_ of both species, and focused sampling of isolated populations might reveal a different pattern as seen in caribou (Dedato et al. 2022). Denser sampling of MD might also help identify key corridors facilitating gene flow (e.g. <u>Fusco et al. 2023)</u>.

## Conclusion

We observed strong signals of climate- and human-induced demographic declines with a remarkable overlap between both census and biogeographic data and historical inferences coming from multiple methods. The dramatic drop in N_e_ in the recent past of all deer populations studied here is robust evidence for a colonial impact through unregulated hunting and changes in land use. Even though WTD & MD populations were heavily depleted, both species showed signs of recovery, particularly white-tailed deer from restocking efforts. While diversity and demographic trends appear positive, there is also a high genetic load in WTD (Wootton et al. 2023), and a low contemporary N_e_/N_c_ ratio in MD. Selection surrounding a wide variety of genes, including some of biological importance in WTD & MD such as genes related to immunity and temperature appear evident. Historical sampling of *Odocoileus* will prove informative for disentangling human-induced selection and genetic load in these wide-ranging cervids.

## Acknowledgements

Camille Kessler was supported by an International Graduate Scholarship, an Ontario Graduate Scholarship, and a French American Charitable Trust Scholarship. This work was supported by NSERC Discovery Grant (Grant Number: RGPIN-2017-03934); ComputeCanada Resources for Research Groups (Grant Number: RRG gme-665-ab); Canadian Foundation for Innovation: John R. Evans Leaders Fund and the Ontario Early Researcher Award (Grant Number: #36905). For providing samples, we thank Alan Cain, Steeve Côté, Anh Dao, Marco Festa-Bianchet, Brad Fulk, Eric Hoffman, Levi Jaster, Daniel Koelsch, Emily Latch, Brent Patterson, Joe Nocera, Don Stewart & Russell Easy, Jonathan Shaw, David Walter and Jon Wheeler. We also thank Jose Alberto López-Alemán for commenting on the text.

We sampled deer from across North America, a stolen land which remains a home to many First Peoples and Trent University is located on the territory of the Michi Saagiig Anishnaabeg. As settlers, we are grateful to have had the opportunity to live and work on this land, and to benefit from it. We would like to show our respect to the First Peoples and thank them for their care, stewardship, and teachings. Miigwetch.

## Data Accessibility

Raw reads for the 45 deer additional individuals were deposited on the NCBI Sequence Read Archive (Accession number PRJNA830519). All scripts are available on GitLab (https://gitlab.com/WiDGeT_TrentU/graduate_theses/-/tree/master/Kessler/CH_02).

## Author contributions

CK and ABAS conceived the study, ABAS collected the samples, CK performed the molecular laboratory work and the bioinformatic analyses with contribution from ABAS for MSMC2, CK and ABAS wrote the manuscript.

