## Supplementary figures and tables for "Genomic analyses capture the human-induced demographic collapse and recovery in a wide-ranging cervid"

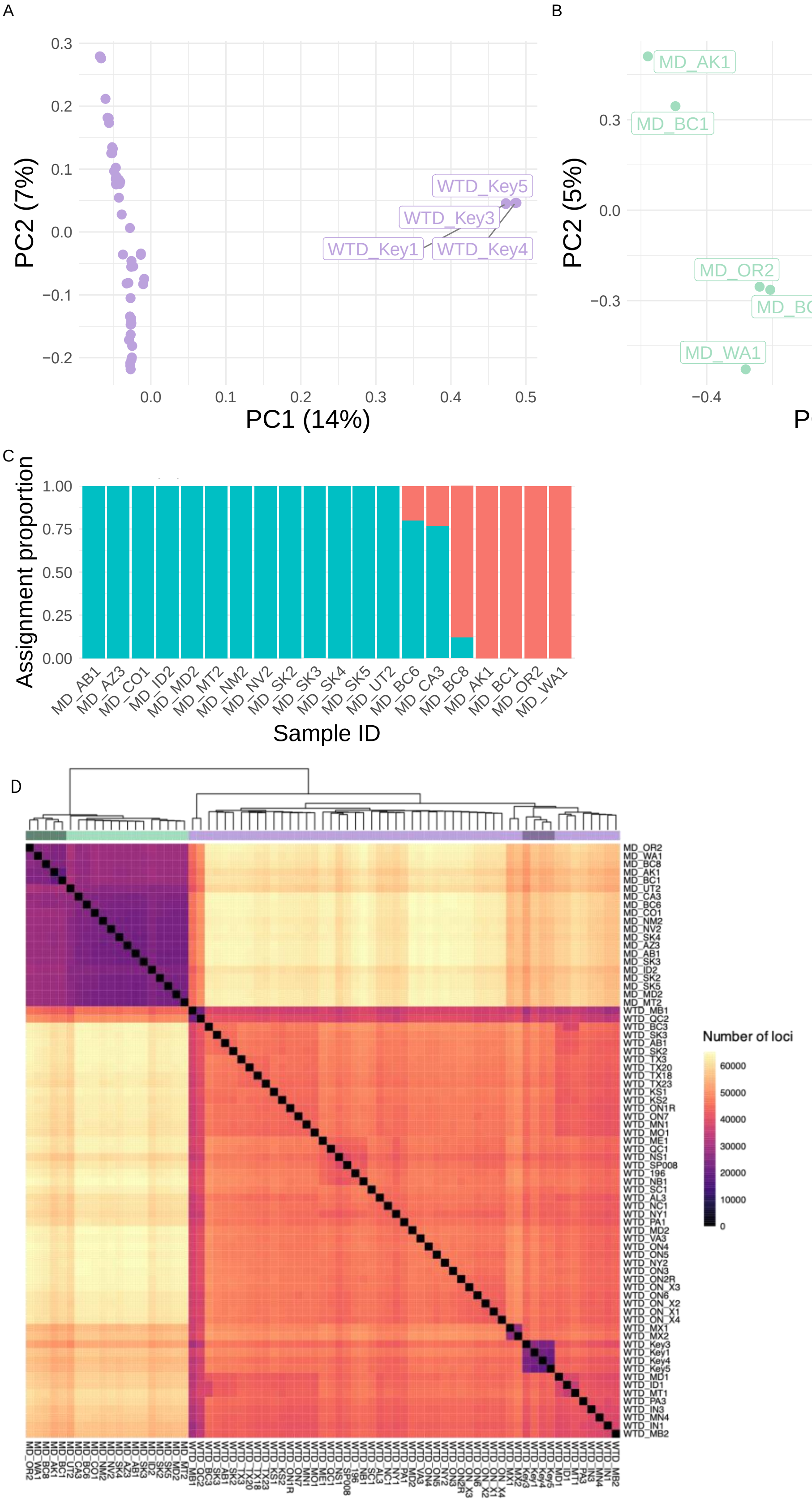


Figure S1: Population structure analyses. (A) PCA of WTD individuals. (B) PCA of MD individuals. (C) Admixture proportions of the MD individuals. (D) Genetic distance in the form of number of divergent loci between individuals, out of 249,380 snps.

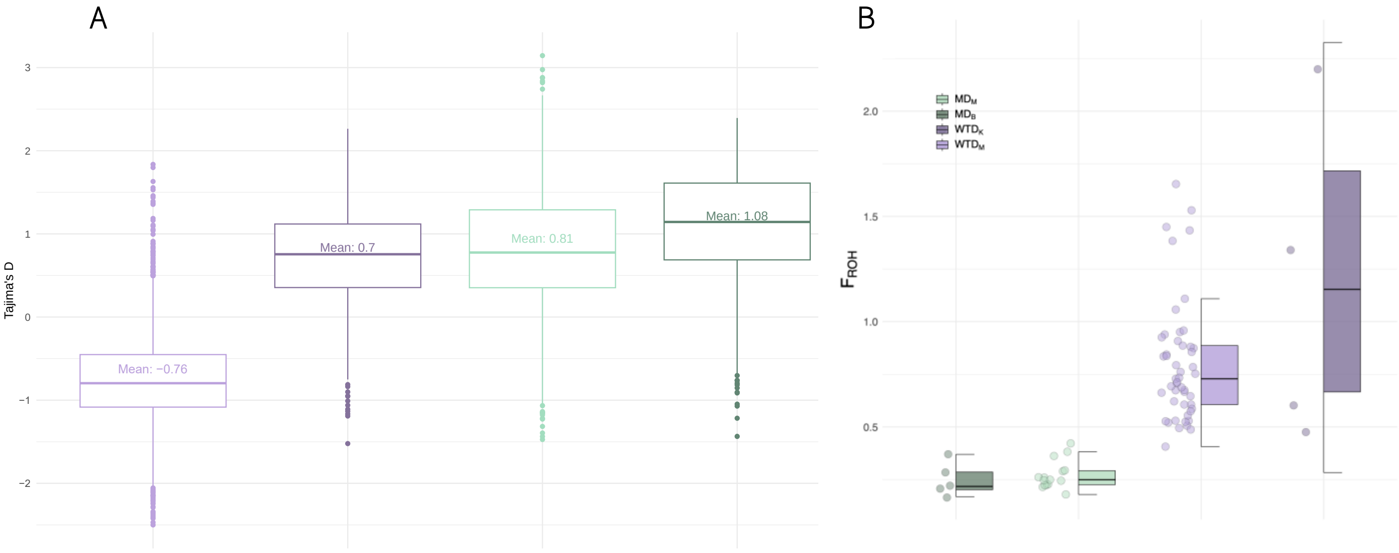


Figure S2: Summary statistics, (A) Genome-wide Tajima's D distribution per population, with average across scaffolds (B) $F_{ROH}$ distribution per individual per population. The y-axis $F_{ROH}$ are expressed as percentage of the genome.

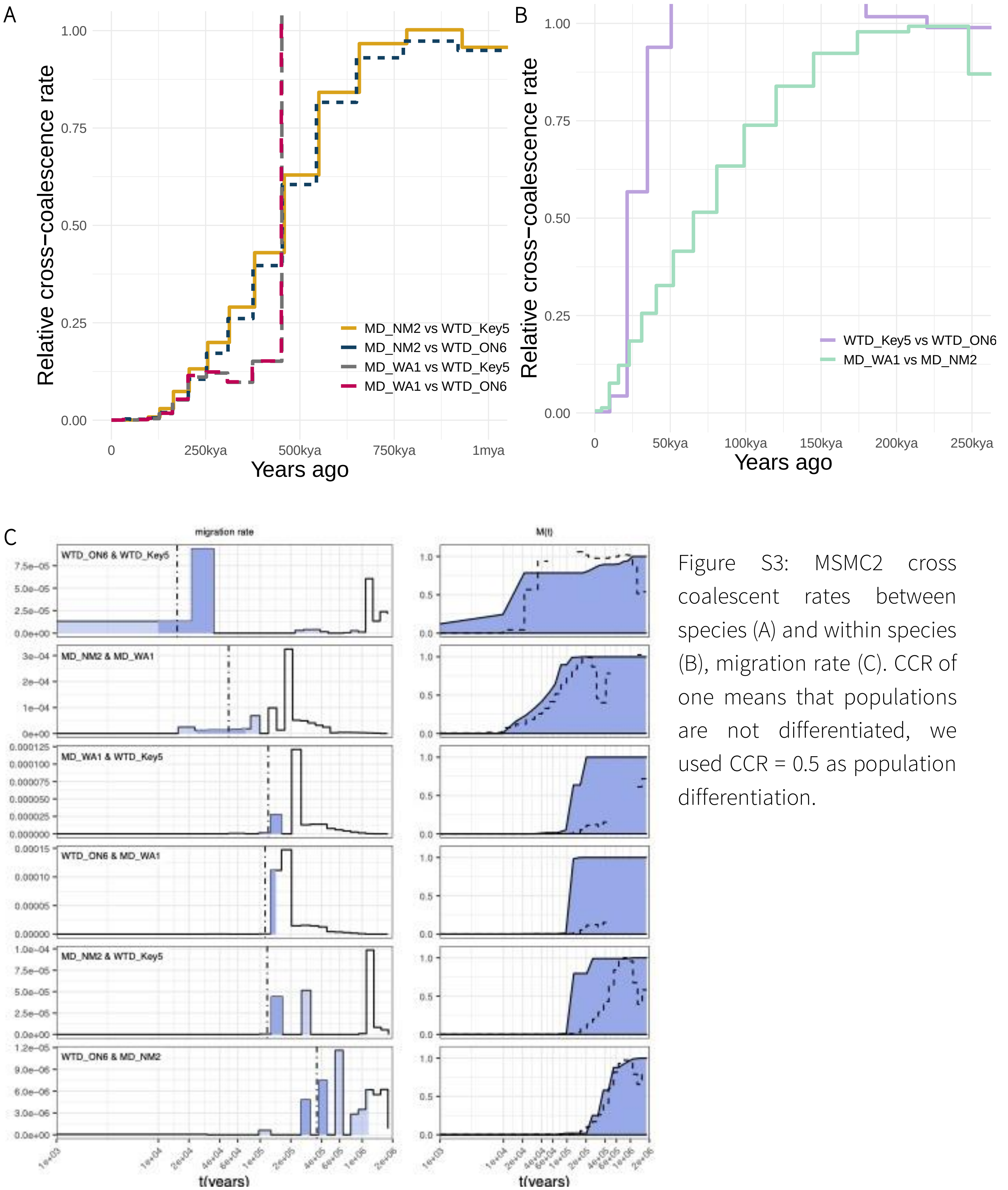


Figure S3: MSMC2 cross coalescent rates between species (A) and within species (B), migration rate (C). CCR of one means that populations are not differentiated, we used CCR = 0.5 as population differentiation.

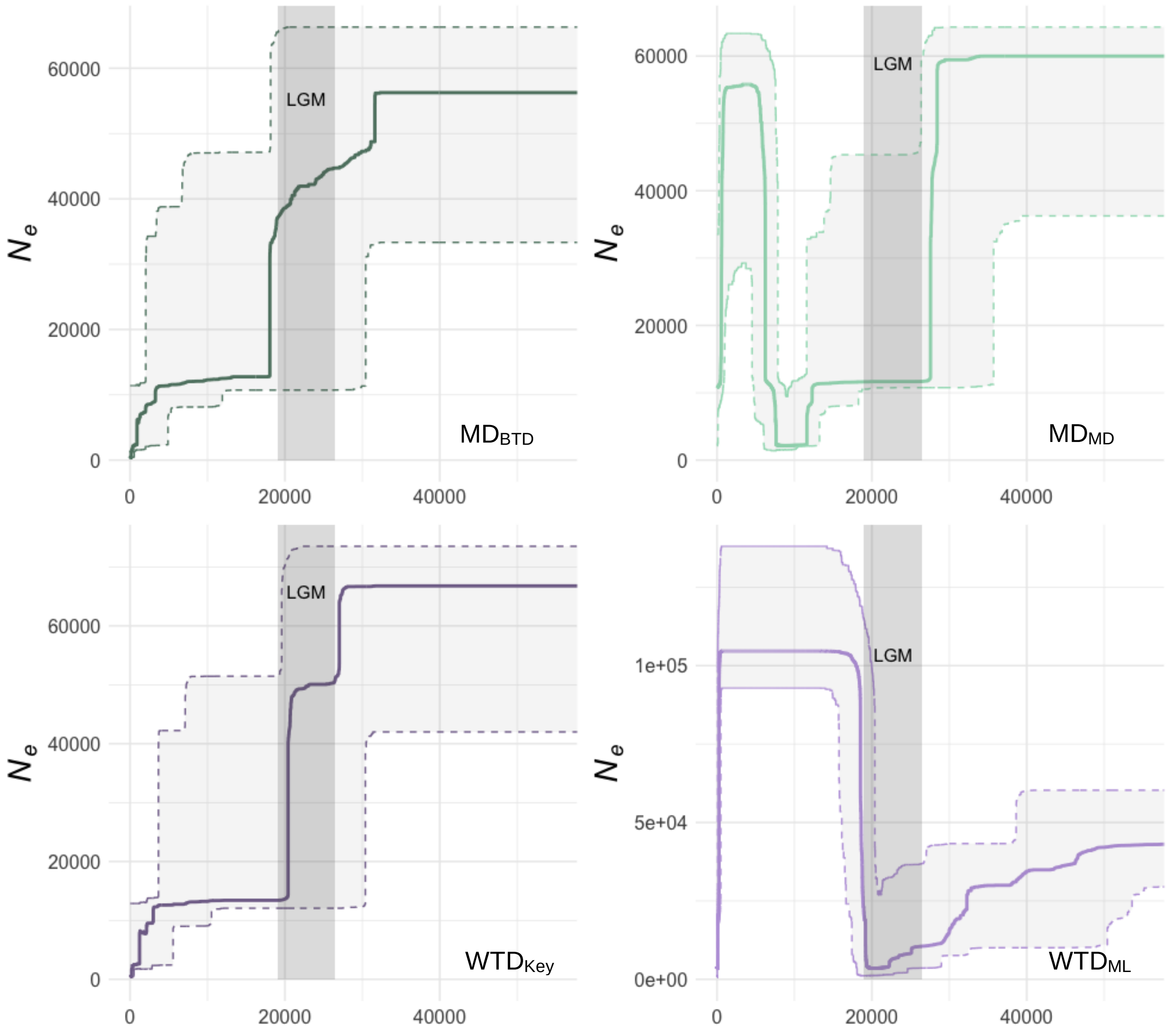


Figure S4: Stairway plot analysis for each population, grey rectangle represents the LGM; dashed lines represent 95% confidence interval.

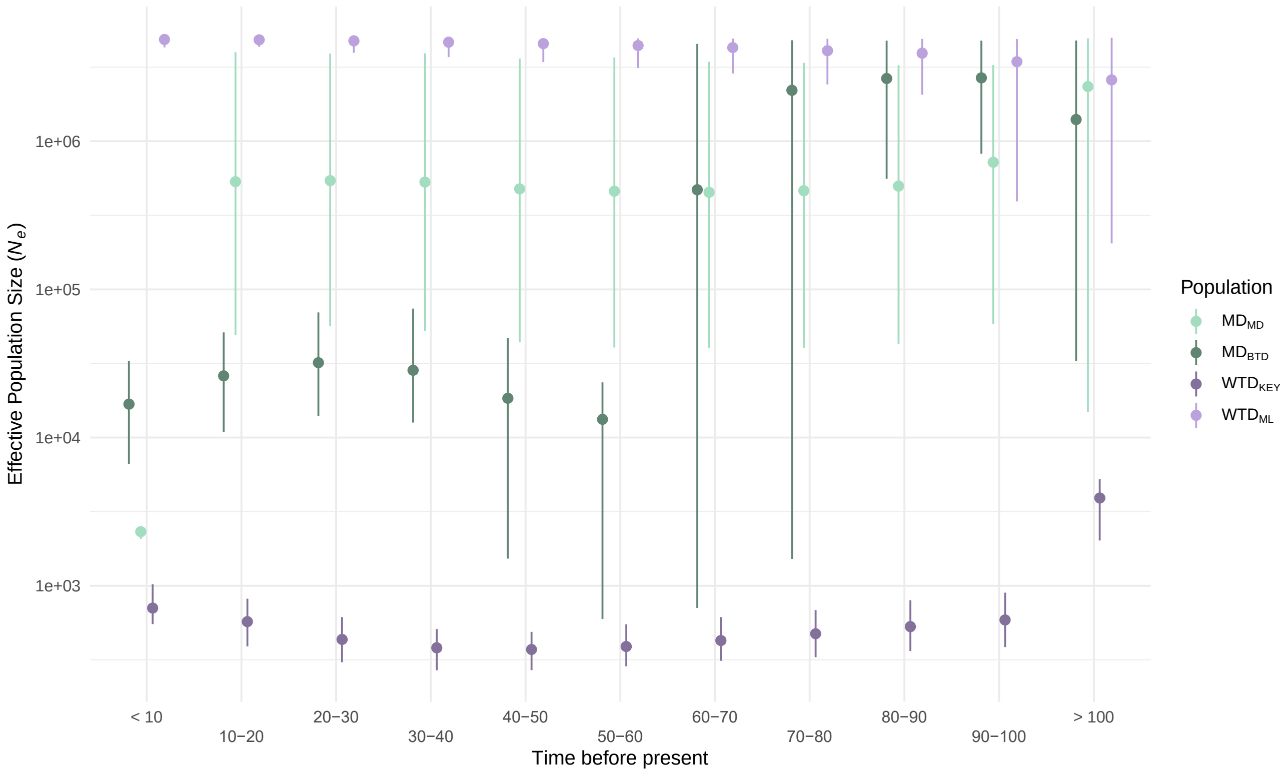


Figure S5: GONE uncertainty for the last 100 years (50 generations), y-axis in log10 scale, x-axis in time bins of 10 years, error bars represent 95% CI, colour represents population.

Figure S6: δaδi parameters uncertainty: Densities of parameters and best parameter value for the $WTD_{KEY}$, $WTD_{ML}$ and all MD comparison (A), and the $MD_{MD}$, $MD_{BTD}$ and $WTD_{ML}$ comparison (B)

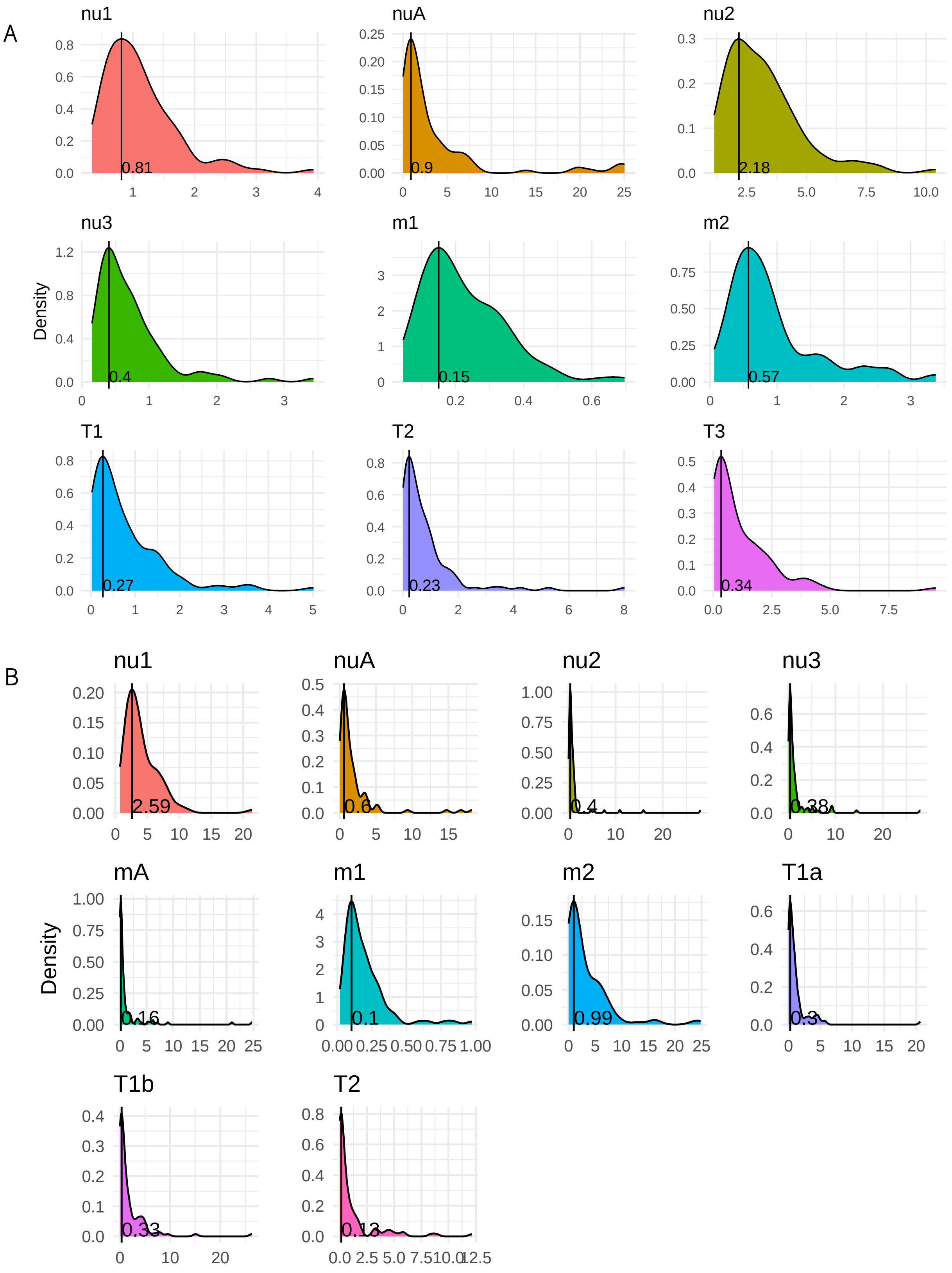

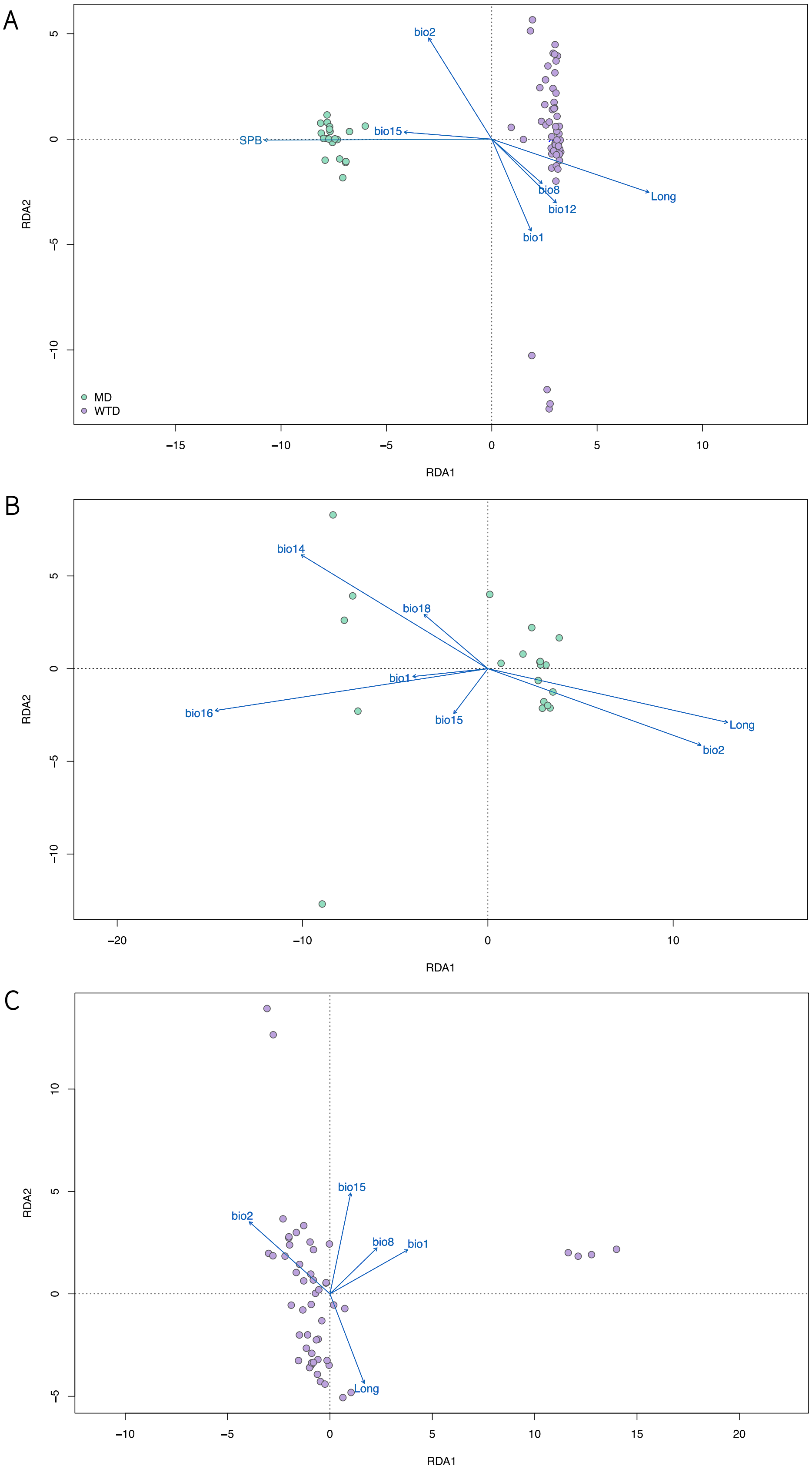


Figure S7: RDA between species (A), within MD (B) and within WTD (C).

Table S1: Sample information; The four re-sequenced samples are highlighted in bold, assigned population taken from the PCA and NGSAdmix analyses with corresponding colour code, sample coordinates were not taken at sampling site but estimated from the approximate sampling location.

| Sample ID | Species | Location | Assigned population | Sex | Latitude (approx) | Longitude (approx) |
|---|---|---|---|---|---|---|
| MD_BC1 | *O_hemionus* | British Columbia | MD$_{BTD}$ | F | 48.69 | -123.32 |
| **MD_WA1** | **O_hemionus** | **Washington** | **MD$_{BTD}$** | **M** | **46.88** | **-122.40** |
| MD_AB1 | O_hemionus | Alberta | MD$_{MD}$ | F | 52.62 | -110.75 |
| MD_AK1 | *O_hemionus* | Alaska | MD$_{BTD}$ | NA | 57.68 | -134.48 |
| MD_AZ3 | *O_hemionus* | Arizona | MD$_{MD}$ | NA | 35.19 | -111.65 |
| MD_BC6 | *O_hemionus* | British Columbia | MD$_{MD}$ | M | 54.78 | -127.16 |
| MD_BC8 | *O_hemionus* | British Columbia | MD$_{BTD}$ | M | 49.15 | -121.95 |
| MD_CA3 | *O_hemionus* | California | MD$_{MD}$ | NA | 35.37 | -119.01 |
| MD_CO1 | *O_hemionus* | Colorado | MD$_{MD}$ | F | 39.48 | -108.09 |
| MD_ID2 | *O_hemionus* | Idaho | MD$_{MD}$ | F | 42.05 | -111.90 |
| MD_MT2 | *O_hemionus* | Montana | MD$_{MD}$ | M | 45.42 | -109.73 |
| **MD_NM2** | **O_hemionus** | **New Mexico** | **MD$_{MD}$** | **M** | **36.18** | **-103.59** |
| MD_NV2 | *O_hemionus* | Nevada | MD$_{MD}$ | M | 41.30 | -115.13 |
| MD_OR2 | *O_hemionus* | Oregon | MD$_{BTD}$ | NA | 44.41 | -122.53 |
| MD_SD2 | *O_hemionus* | South Dakota | MD$_{MD}$ | F | 43.22 | -103.45 |
| MD_SK2 | *O_hemionus* | Saskatchewan | MD$_{MD}$ | NA | 51.02 | -106.43 |
| MD_SK3 | *O_hemionus* | Saskatchewan | MD$_{MD}$ | NA | 51.02 | -106.43 |
| MD_SK4 | *O_hemionus* | Saskatchewan | MD$_{MD}$ | NA | 50.74 | -107.81 |
| MD_SK5 | *O_hemionus* | Saskatchewan | MD$_{MD}$ | NA | 50.74 | -107.81 |
| MD_UT2 | *O_hemionus* | Utah | MD$_{MD}$ | M | 40.97 | -111.69 |
| WTD_AB1 | *O_virginianus* | Alberta | WTD$_{ML}$ | F | 52.78 | -111.01 |
| WTD_AL3 | *O_virginianus* | Alabama | WTD$_{ML}$ | NA | 32.38 | -85.68 |
| WTD_BC3 | *O_virginianus* | British Columbia | WTD$_{ML}$ | M | 49.51 | -115.76 |
| WTD_IN1 | *O_virginianus* | Indiana | WTD$_{ML}$ | NA | 41.25 | -87.22 |
| WTD_IN3 | *O_virginianus* | Indiana | WTD$_{ML}$ | NA | 39.25 | -85.04 |
| WTD_Key1 | *O_virginianus* | Florida | WTD$_{KEY}$ | NA | 24.69 | -81.32 |
| WTD_Key3 | *O_virginianus* | Florida | WTD$_{KEY}$ | NA | 24.69 | -81.32 |
| WTD_Key4 | *O_virginianus* | Florida | WTD$_{KEY}$ | NA | 24.69 | -81.32 |
| **WTD_Key5** | **O_virginianus** | **Florida** | **WTD$_{KEY}$** | **NA** | **24.69** | **-81.32** |
| WTD_KS1 | *O_virginianus* | Kansas | WTD$_{ML}$ | F | 38.69 | -100.81 |
| WTD_KS2 | *O_virginianus* | Kansas | WTD$_{ML}$ | M | 38.43 | -96.18 |
| WTD_MB1 | *O_virginianus* | Manitoba | WTD$_{ML}$ | M | 51.52 | -100.39 |
| WTD_MB2 | *O_virginianus* | Manitoba | WTD$_{ML}$ | M | 50.61 | -97.56 |
| WTD_MD2 | *O_virginianus* | Maryland | WTD$_{ML}$ | F | 39.62 | -78.61 |
| WTD_ME1 | *O_virginianus* | Maine | WTD$_{ML}$ | F | 43.66 | -70.63 |
| WTD_MN1 | *O_virginianus* | Minnesota | WTD$_{ML}$ | M | 46.35 | -94.20 |
| WTD_MN4 | *O_virginianus* | Minnesota | WTD$_{ML}$ | M | 43.67 | -92.08 |
| WTD_MO1 | *O_virginianus* | Missouri | WTD$_{ML}$ | M | 38.38 | -91.90 |

| | | | | | | | |
|---|---|---|---|---|---|---|---|
| WTD_MT1 | *O_virginianus* | Montana | $WTD_{ML}$ | | M | 47.52 | -113.71 |
| WTD_MX1 | *O_virginianus* | Mexico | $WTD_{ML}$ | | M | 30.72 | -108.73 |
| WTD_MX2 | *O_virginianus* | Mexico | $WTD_{ML}$ | | M | 30.72 | -108.73 |
| WTD_NB1 | *O_virginianus* | New Brunswick | $WTD_{ML}$ | | M | 45.68 | -66.69 |
| WTD_NC1 | *O_virginianus* | North Carolina | $WTD_{ML}$ | | M | 34.65 | -77.47 |
| WTD_NS1 | *O_virginianus* | Nova Scotia | $WTD_{ML}$ | | M | 45.05 | -63.16 |
| WTD_NY1 | *O_virginianus* | New York | $WTD_{ML}$ | | F | 42.64 | -73.74 |
| WTD_NY2 | *O_virginianus* | New York | $WTD_{ML}$ | | F | 43.02 | -76.17 |
| WTD_ON1R | *O_virginianus* | Ontario | $WTD_{ML}$ | | M | 48.24 | -89.12 |
| WTD_ON2R | *O_virginianus* | Ontario | $WTD_{ML}$ | | M | 45.12 | -75 |
| WTD_ON3 | *O_virginianus* | Ontario | $WTD_{ML}$ | | F | 42.48 | -82 |
| WTD_ON4 | *O_virginianus* | Ontario | $WTD_{ML}$ | | M | 44.92 | -83.4 |
| WTD_ON5 | *O_virginianus* | Ontario | $WTD_{ML}$ | | M | 46.36 | -84 |
| **WTD_ON6** | **O_virginianus** | **Ontario** | $\mathbf{WTD_{ML}}$ | | **M** | **44.53** | **-78.05** |
| WTD_ON7 | *O_virginianus* | Ontario | $WTD_{ML}$ | | M | 49.76 | -94.48 |
| WTD_ONX1 | *O_virginianus* | Ontario | $WTD_{ML}$ | | F | 44.58 | -78.07 |
| WTD_ONX2 | *O_virginianus* | Ontario | $WTD_{ML}$ | | F | 44.58 | -78.07 |
| WTD_ONX3 | *O_virginianus* | Ontario | $WTD_{ML}$ | | F | 44.58 | -78.25 |
| WTD_ONX4 | *O_virginianus* | Ontario | $WTD_{ML}$ | | M | 44.62 | -78.13 |
| WTD_PA1 | *O_virginianus* | Pennsylvania | $WTD_{ML}$ | | M | 39.96 | -76.72 |
| WTD_PA3 | *O_virginianus* | Pennsylvania | $WTD_{ML}$ | | F | 39.89 | -80.08 |
| WTD_QC1 | *O_virginianus* | Quebec | $WTD_{ML}$ | | M | 46.9 | -70.43 |
| WTD_QC2 | *O_virginianus* | Quebec | $WTD_{ML}$ | | F | 45.40 | -71.89 |
| WTD_SC1 | *O_virginianus* | South Carolina | $WTD_{ML}$ | | F | 33.41 | -80.41 |
| WTD_SD1 | *O_virginianus* | South Dakota | $WTD_{ML}$ | | M | 44.44 | -102.68 |
| WTD_SK2 | *O_virginianus* | Saskatchewan | $WTD_{ML}$ | | M | 52.86 | -102.36 |
| WTD_SK3 | *O_virginianus* | Saskatchewan | $WTD_{ML}$ | | M | 53.63 | -109.2 |
| WTD_SP008 | *O_virginianus* | St-Pierre et Miquelon | $WTD_{ML}$ | | F | 46.86 | -56.24 |
| WTD_TX18 | *O_virginianus* | Texas | $WTD_{ML}$ | | F | 30.71 | -94.81 |
| WTD_TX20 | *O_virginianus* | Texas | $WTD_{ML}$ | | F | 33.18 | -99.27 |
| WTD_TX23 | *O_virginianus* | Texas | $WTD_{ML}$ | | M | 31.78 | -101.07 |
| WTD_TX3 | *O_virginianus* | Texas | $WTD_{ML}$ | | F | 29.29 | -99.01 |
| WTD_VA3 | *O_virginianus* | Virginia | $WTD_{ML}$ | | F | 38.13 | -78.16 |
| WTD_196 | *O_virginianus* | Anticosti Island | $WTD_{ML}$ | | NA | 49.47 | -63.09 |
| WTD_ID1 | *O_virginianus* | Idaho | $WTD_{ML}$ | | F | 48.68 | -116.36 |

Table S2: List of 3D models run in dadi_pipeline (Portik et al., 2017) for two comparisons

| Comparison | Model | Reference |
|---|---|---|
| $WTD_{KEY}$ - $WTK_{ML}$ - MD | split_nomig | Portik et al., 2017 |
| | split_symmig_adjacent | |
| | refugia_adj_1 | |
| | refugia_adj_2 | |
| | refugia_adj_3 | |
| | ancmig_adj_1 | |
| | ancmig_adj_2 | |
| | ancmig_adj_3 | |
| | ancmig_2_size | Barratt et al., 2018 |
| | split_sym_mig_adjacent_var1 | Firneno et al., 2020 |
| | split_sym_mig_adjacent_var2 | |
| $MD_{BTD}$ - $MD_{MD}$ - $WTD_{ML}$ | split_nomig | Portik et al., 2017 |
| | split_symmig_all | |
| | split_symmig_adjacent | |
| | refugia_adj_1 | |
| | refugia_adj_2 | |
| | refugia_adj_3 | |
| | refugia_adj_2_var_sym | Firneno et al., 2020 |
| | refugia_adj_3_var_sym | |
| | refugia_adj_2_var_uni | |
| | refugia_adj_3_var_uni | |
| | split_sym_mig_adjacent_var1 | |
| | split_uni_mig_adjacent_var1 | |

#### Table S3: Bioclimatic variables from WorldClim2

| Code | Bioclimatic variable |
|---|---|
| BIO1 | Annual Mean Temperature |
| BIO2 | Mean Diurnal Range (Mean of monthly (max temp - min temp)) |
| BIO3 | Isothermality (BIO2/BIO7) (×100) |
| BIO4 | Temperature Seasonality (standard deviation ×100) |
| BIO5 | Max Temperature of Warmest Month |
| BIO6 | Min Temperature of Coldest Month |
| BIO7 | Temperature Annual Range (BIO5-BIO6) |
| BIO8 | Mean Temperature of Wettest Quarter |
| BIO9 | Mean Temperature of Driest Quarter |
| BIO10 | Mean Temperature of Warmest Quarter |
| BIO11 | Mean Temperature of Coldest Quarter |
| BIO12 | Annual Precipitation |
| BIO13 | Precipitation of Wettest Month |
| BIO14 | Precipitation of Driest Month |
| BIO15 | Precipitation Seasonality (Coefficient of Variation) |
| BIO16 | Precipitation of Wettest Quarter |
| BIO17 | Precipitation of Driest Quarter |
| BIO18 | Precipitation of Warmest Quarter |
| BIO19 | Precipitation of Coldest Quarter |

#### Table S4: Weighted $F_{st}$ values between populations

| | $WTD_{ML}$ | $WTD_{KEY}$ | $MD_{MD}$ |
|---|---|---|---|
| $WTD_{KEY}$ | 0.17 | | |
| $MD_{MD}$ | 0.4 | 0.58 | |
| $MD_{BTD}$ | 0.36 | 0.56 | 0.12 |

#### Table S5: Best model parameter values from δaδi per comparison and corresponding 95% CI computed on 100 bootstraps. Converted parameters values found by solving for $\Theta = 4N_{ref}\mu L$.

| | | nu | | | | m | | | T | | | | |
|---|---|---|---|---|---|---|---|---|---|---|---|---|---|
| | **Theta** | **nu1** | **nu2** | **nu3** | **nuA** | **m1** | **m2** | **mA** | **T1** | **T1a** | **T1b** | **T2** | **T3** |
| $WTD_{KEY}$ & $WTD_{ML}$ & MD | | | | | | | | | | | | | |
| Parameters | 21'150.17 | 0.56 | 1.73 | 0.20 | 4.02 | 0.86 | 2.15 | - | 0.57 | - | - | 0.12 | 0.02 |
| 2.5% CI | | 0.44 | 1.34 | 0.18 | 0.06 | 0.07 | 0.11 | - | 0.03 | - | - | 0.02 | 0.06 |
| 97.5% CI | | 2.61 | 7.47 | 2.04 | 24.23 | 0.49 | 2.70 | - | 3.27 | - | - | 3.95 | 4.10 |
| Converted parameters | | 55'432 | 171'381 | 19'344 | 397'956 | 4.4E-06 | 1.1E-05 | - | 226'081 | - | - | 47'471 | 6'725 |
| $WTD_{ML}$ & $MD_{BTD}$ & $MD_{MD}$ | | | | | | | | | | | | | |
| Parameters | 20'887.17 | 2.09 | 0.29 | 0.27 | 0.84 | 0.79 | 0.79 | 0.02 | - | 0.11 | 0.86 | 0.04 | - |
| 2.5% CI | | 1.19 | 0.15 | 0.10 | 0.08 | 0.05 | 0.10 | 0.03 | - | 0.03 | 0.05 | 0.02 | - |
| 97.5% CI | | 9.82 | 9.34 | 9.28 | 12.18 | 0.72 | 16.21 | 17.37 | - | 5.07 | 8.83 | 7.19 | - |
| Converted parameters | | 204'098 | 28'412 | 25'941 | 81'573 | 4.1E-06 | 4.0E-06 | 1.0E-07 | - | 44'420 | 335'120 | 13'869 | - |

### Table S6: 249 genes within 25kbp of iHS peaks.

| Entry | Gene Name | Function (uniprot.org) | Category | Pop |
|---|---|---|---|---|
| O14497 | *ARID1A BAF250 BAF250A C1orf4 OSA1 SMARCF1* | FUNCTION: Involved in transcriptional activation and repression of select genes by chromatin remodeling (alteration of DNA-nucleosome topology). Component of SWI/SNF chromatin remodeling complexes that carry out key enzymatic activities, changing chromatin structure by altering DNA-histone contacts within a nucleosome in an ATP-dependent manner. Binds DNA non-specifically. Belongs to the neural progenitors-specific chromatin remodeling complex (npBAF complex) and the neuron-specific chromatin remodeling complex (nBAF complex). During neural development a switch from a stem/progenitor to a postmitotic chromatin remodeling mechanism occurs as neurons exit the cell cycle and become committed to their adult state. The transition from proliferating neural stem/progenitor cells to postmitotic neurons requires a switch in subunit composition of the npBAF and nBAF complexes. As neural progenitors exit mitosis and differentiate into neurons, npBAF complexes which contain ACTL6A/BAF53A and PHF10/BAF45A, are exchanged for homologous alternative ACTL6B/BAF53B and DPF1/BAF45B or DPF3/BAF45C subunits in neuron-specific complexes (nBAF). The npBAF complex is essential for the self-renewal/proliferative capacity of the multipotent neural stem cells. The nBAF complex along with CREST plays a role regulating the activity of genes essential for dendrite growth (By similarity). {ECO:0000250\|UniProtKB:A2BH40, ECO:0000303\|PubMed:12672490, ECO:0000303\|PubMed:22952240, ECO:0000303\|PubMed:26601204}. | *Development* | $WTD_{ML}$ |
| O60229 | *KALRN DUET DUO HAPIP TRAD* | FUNCTION: Promotes the exchange of GDP by GTP. Activates specific Rho GTPase family members, thereby inducing various signaling mechanisms that regulate neuronal shape, growth, and plasticity, through their effects on the actin cytoskeleton. Induces lamellipodia independent of its GEF activity. {ECO:0000269\|PubMed:10023074}. | *Development* | $WTD_{ML}$ |
| P12272 | *PTHLH PTHRP* | FUNCTION: Neuroendocrine peptide which is a critical regulator of cellular and organ growth, development, migration, differentiation and survival and of epithelial calcium ion transport. Regulates endochondral bone development and epithelial-mesenchymal interactions during the formation of the mammary glands and teeth. Required for skeletal homeostasis. Promotes mammary mesenchyme differentiation and bud outgrowth by modulating mesenchymal cell responsiveness to BMPs. Up-regulates BMPR1A expression in the mammary mesenchyme and this increases the sensitivity of these cells to BMPs and allows them to respond to BMP4 in a paracrine and/or autocrine fashion. BMP4 signaling in the mesenchyme, in turn, triggers epithelial outgrowth and augments MSX2 expression, which causes the mammary mesenchyme to inhibit hair follicle formation within the nipple sheath (By similarity). Promotes colon cancer cell migration and invasion in an integrin alpha-6/beta-1-dependent manner through activation of Rac1. {ECO:0000250, ECO:0000269\|PubMed:20637541}.; FUNCTION: Osteostatin is a potent inhibitor of osteoclastic bone resorption. {ECO:0000269\|PubMed:20637541}. | *Development* | $WTD_{ML}$ |
| Q19T08 | *ECSCR ECSM2* | FUNCTION: Regulates endothelial chemotaxis and tube formation. Has a role in angiogenesis and apoptosis via modulation of the actin cytoskeleton and facilitation of proteasomal degradation of the apoptosis inhibitors BIRC3/IAP1 and BIRC2/IAP2. {ECO:0000269\|PubMed:18556573, ECO:0000269\|PubMed:19416853}. | *Development* | $WTD_{ML}$ |
| Q4G0U5 | *CFAP221 PCDP1* | FUNCTION: May play a role in cilium morphogenesis. {ECO:0000250\|UniProtKB:A9Q751}. | *Development* | $WTD_{ML}$ |
| Q5HYA8 | *TMEM67 MKS3* | FUNCTION: Required for ciliary structure and function. Part of the tectonic-like complex which is required for tissue-specific ciliogenesis and may regulate ciliary membrane composition (By similarity). Involved in centrosome migration to the apical cell surface during early ciliogenesis. Involved in the regulation of cilia length and appropriate number through the control of centrosome duplication. Is a key regulator of stereociliary bundle orientation (By similarity). Required for epithelial cell branching morphology. Essential for endoplasmic reticulum-associated degradation (ERAD) of surfactant protein C (SFTPC). Involved in the negative regulation of canonical Wnt signaling, and activation of the non-canonical cascade stimulated by WNT5A (PubMed:26035863). In non-canonical Wnt signaling, it may act as ROR2 coreceptor (By similarity). {ECO:0000250\|UniProtKB:Q8BR76, ECO:0000269\|PubMed:17185389, ECO:0000269\|PubMed:19515853, ECO:0000269\|PubMed:19596800, ECO:0000269\|PubMed:19815549, ECO:0000269\|PubMed:26035863}. | *Development* | $MD_{BTD}$ |
| Q5VUG0 | *SFMBT2 KIAA1617* | FUNCTION: Transcriptional repressor of HOXB13 gene. {ECO:0000269\|PubMed:23385818}. | *Development* | $WTD_{KEY}$ |
| Q68J44 | *DUSP29 DUPD1 DUSP27* | FUNCTION: Dual specificity phosphatase able to dephosphorylate phosphotyrosine, phosphoserine and phosphothreonine residues within the same substrate, with a preference for phosphotyrosine as a substrate (PubMed:17498703). Involved in the modulation of intracellular signaling cascades. In skeletal muscle regulates systemic glucose homeostasis by activating, AMPK, an energy sensor protein kinase (By similarity). Affects MAP kinase signaling though modulation of the MAPK1/2 cascade in skeletal muscle promoting muscle cell differentiation, development and atrophy (By similarity). {ECO:0000250\|UniProtKB:Q8BK84, ECO:0000269\|PubMed:17498703}. | *Development* | $WTD_{KEY}$ |
| Q6PI78 | *TMEM65* | FUNCTION: May play an important role in cardiac development and function. May regulate cardiac conduction and the function of the gap junction protein GJA1. May contribute to the stability and proper localization of GJA1 to cardiac intercalated disk thereby regulating gap junction communication (By similarity). May also play a role in the regulation of mitochondrial respiration and mitochondrial DNA copy number maintenance (PubMed:28295037). {ECO:0000250\|UniProtKB:Q4VAE3, ECO:0000269\|PubMed:28295037}. | *Development* | $WTD_{KEY}$ |
| Q7L9L4 | *MOB1B MOB4A MOBKL1 A* | FUNCTION: Activator of LATS1/2 in the Hippo signaling pathway which plays a pivotal role in organ size control and tumor suppression by restricting proliferation and promoting apoptosis. The core of this pathway is composed of a kinase cascade wherein STK3/MST2 and STK4/MST1, in complex with its regulatory protein SAV1, phosphorylates and activates LATS1/2 in complex with its regulatory protein MOB1, which in turn phosphorylates and inactivates YAP1 oncoprotein and WWTR1/TAZ. Phosphorylation of YAP1 by LATS1/2 inhibits its translocation into the nucleus to regulate cellular genes important for cell proliferation, cell death, and cell migration. Stimulates the kinase activity of STK38L. {ECO:0000269\|PubMed:15067004, ECO:0000269\|PubMed:19739119}. | *Development* | $WTD_{KEY}$ |
| Q8IZT6 | *ASPM MCPH5* | FUNCTION: Involved in mitotic spindle regulation and coordination of mitotic processes. The function in regulating microtubule dynamics at spindle poles including spindle orientation, astral microtubule density and poleward microtubule flux seems to depend on the association with the katanin complex formed by KATNA1 and KATNB1. Enhances the microtubule lattice severing activity of KATNA1 by recruiting the katanin complex to microtubules. Can block microtubule minus-end growth and reversely this function can be enhanced by the katanin complex (PubMed:28436967). May have a preferential role in regulating neurogenesis. {ECO:0000269\|PubMed:12355089, ECO:0000269\|PubMed:15972725, ECO:0000269\|PubMed:28436967}. | *Development* | $MD_{MD}$ |
| Q8WVF2 | *UCMA C10orf49* | FUNCTION: May be involved in the negative control of osteogenic differentiation of osteochondrogenic precursor cells in peripheral zones of fetal cartilage and at the cartilage-bone interface. {ECO:0000250}. | *Development* | $WTD_{ML}$ |

| | | | | |
|---|---|---|---|---|
| Q8WYB5 | *KAT6B KIAA0383 MORF MOZ2 MYST4* | FUNCTION: Histone acetyltransferase which may be involved in both positive and negative regulation of transcription. Required for RUNX2-dependent transcriptional activation. May be involved in cerebral cortex development. Component of the MOZ/MORF complex which has a histone H3 acetyltransferase activity. {ECO:0000269\|PubMed:10497217, ECO:0000269\|PubMed:11965546, ECO:0000269\|PubMed:16387653}. | *Development* | $WTD_{KEY}$ |
| Q9H799 | *CPLANE1 C5orf42 JBTS17* | FUNCTION: Involved in ciliogenesis (PubMed:25877302). Involved in the establishment of cell polarity required for directional cell migration. Proposed to act in association with the CPLANE (ciliogenesis and planar polarity effectors) complex. Involved in recruitment of peripheral IFT-A proteins to basal bodies (By similarity). {ECO:0000250\|UniProtKB:Q8CE72, ECO:0000305\|PubMed:25877302}. | *Development* | $WTD_{ML}$ |
| Q9UH90 | *FBXO40 FBX40 KIAA1195* | FUNCTION: Probable substrate-recognition component of the SCF (SKP1-CUL1-F-box protein)-type E3 ubiquitin ligase complex that may function in myogenesis. {ECO:0000250}. | *Development* | $MD_{MD}$ |
| Q9UMX5 | *NENF CIR2 SPUF* | FUNCTION: Acts as a neurotrophic factor in postnatal mature neurons enhancing neuronal survival (PubMed:31536960). Promotes cell proliferation and neurogenesis in undifferentiated neural progenitor cells at the embryonic stage and inhibits differentiation of astrocytes (By similarity). Its neurotrophic activity is exerted via MAPK1/ERK2, MAPK3/ERK1 and AKT1/AKT pathways (By similarity). Neurotrophic activity is enhanced by binding to heme (By similarity). Acts also as an anorexigenic neurotrophic factor that contributes to energy balance (By similarity). {ECO:0000250\|UniProtKB:Q9CQ45, ECO:0000269\|PubMed:31536960}. | *Development* | $MD_{MD}$ |
| P98164 | *LRP2* | FUNCTION: Multiligand endocytic receptor (By similarity). Acts together with CUBN to mediate endocytosis of high-density lipoproteins (By similarity). Mediates receptor-mediated uptake of polybasic drugs such as aprotinin, aminoglycosides and polymyxin B (By similarity). In the kidney, mediates the tubular uptake and clearance of leptin (By similarity). Also mediates transport of leptin across the blood-brain barrier through endocytosis at the choroid plexus epithelium (By similarity). Endocytosis of leptin in neuronal cells is required for hypothalamic leptin signaling and leptin-mediated regulation of feeding and body weight (By similarity). Mediates endocytosis and subsequent lysosomal degradation of CST3 in kidney proximal tubule cells (By similarity). Mediates renal uptake of 25-hydroxyvitamin D3 in complex with the vitamin D3 transporter GC/DBP (By similarity). Mediates renal uptake of metallothionein-bound heavy metals (PubMed:15126248). Together with CUBN, mediates renal reabsorption of myoglobin (By similarity). Mediates renal uptake and subsequent lysosomal degradation of APOM (By similarity). Plays a role in kidney selenium homeostasis by mediating renal endocytosis of selenoprotein SEPP1 (By similarity). Mediates renal uptake of the antiapoptotic protein BIRC5/survivin which may be important for functional integrity of the kidney (PubMed:23825075). Mediates renal uptake of matrix metalloproteinase MMP2 in complex with metalloproteinase inhibitor TIMP1 (By similarity). Mediates endocytosis of Sonic hedgehog protein N-product (ShhN), the active product of SHH (By similarity). Also mediates ShhN transcytosis (By similarity). In the embryonic neuroepithelium, mediates endocytic uptake and degradation of BMP4, is required for correct SHH localization in the ventral neural tube and plays a role in patterning of the ventral telencephalon (By similarity). Required at the onset of neurulation to sequester SHH on the apical surface of neuroepithelial cells of the rostral diencephalon ventral midline and to control PTCH1-dependent uptake and intracellular trafficking of SHH (By similarity). During neurulation, required in neuroepithelial cells for uptake of folate bound to the folate receptor FOLR1 which is necessary for neural tube closure (By similarity). In the adult brain, negatively regulates BMP signaling in the subependymal zone which enables neurogenesis to proceed (By similarity). In astrocytes, mediates endocytosis of ALB which is required for the synthesis of the neurotrophic factor oleic acid (By similarity). Involved in neurite branching (By similarity). During optic nerve development, required for SHH-mediated migration and proliferation of oligodendrocyte precursor cells (By similarity). Mediates endocytic uptake and clearance of SHH in the retinal margin which protects retinal progenitor cells from mitogenic stimuli and keeps them quiescent (By similarity). Plays a role in reproductive organ development by mediating uptake in reproductive tissues of androgen and estrogen bound to the sex hormone binding protein SHBG (By similarity). Mediates endocytosis of angiotensin-2 (By similarity). Also mediates endocytosis of angiotensis 1-7 (By similarity). Binds to the complex composed of beta-amyloid protein 40 and CLU/APOJ and mediates its endocytosis and lysosomal degradation (By similarity). Required for embryonic heart development (By similarity). Required for normal hearing, possibly through interaction with estrogen in the inner ear (By similarity). {ECO:0000250\|UniProtKB:A2ARV4, ECO:0000250\|UniProtKB:C0HL13, ECO:0000250\|UniProtKB:P98158, ECO:0000269\|PubMed:15126248, ECO:0000269\|PubMed:23825075}. | *Development, physiology* | $MD_{MD}$ |
| Q13113 | *PDZK1IP 1 MAP17* | FUNCTION: May play an important role in tumor biology. | *Development, physiology* | $WTD_{ML}$ |
| Q9BSE2 | *TMEM79 MATT* | FUNCTION: Contributes to the epidermal integrity and skin barrier function. Plays a role in the lamellar granule (LG) secretory system and in the stratum corneum (SC) epithelial cell formation (By similarity). {ECO:0000250}. | *Development, physiology* | $MD_{MD}$ |
| O95760 | *IL33 C9orf26 IL1F11 NFHEV* | FUNCTION: Cytokine that binds to and signals through the IL1RL1/ST2 receptor which in turn activates NF-kappa-B and MAPK signaling pathways in target cells (PubMed:16286016). Involved in the maturation of Th2 cells inducing the secretion of T-helper type 2-associated cytokines. Also involved in activation of mast cells, basophils, eosinophils and natural killer cells. Acts as a chemoattractant for Th2 cells, and may function as an 'alarmin', that amplifies immune responses during tissue injury (PubMed:17853410, PubMed:18836528). Induces rapid UCP2-dependent mitochondrial rewiring that attenuates the generation of reactive oxygen species and preserves the integrity of Krebs cycle required for persistent production of itaconate and subsequent GATA3-dependent differentiation of inflammation-resolving alternatively activated macrophages. {ECO:0000250\|UniProtKB:Q8BVZ5, ECO:0000269\|PubMed:16286016, ECO:0000269\|PubMed:17853410, ECO:0000269\|PubMed:18836528}.; FUNCTION: In quiescent endothelia the uncleaved form is constitutively and abundantly expressed, and acts as a chromatin-associated nuclear factor with transcriptional repressor properties, it may sequester nuclear NF-kappaB/RELA, lowering expression of its targets (PubMed:21734074). This form is rapidly lost upon angiogenic or pro-inflammatory activation (PubMed:18787100). {ECO:0000269\|PubMed:18787100, ECO:0000269\|PubMed:21734074}. | *Immunity* | $WTD_{ML}$ |

| | | | | |
|---|---|---|---|---|
| P01909 | *HLA-DQA1* | FUNCTION: Binds peptides derived from antigens that access the endocytic route of antigen presenting cells (APC) and presents them on the cell surface for recognition by the CD4 T-cells. The peptide binding cleft accommodates peptides of 10-30 residues. The peptides presented by MHC class II molecules are generated mostly by degradation of proteins that access the endocytic route, where they are processed by lysosomal proteases and other hydrolases. Exogenous antigens that have been endocytosed by the APC are thus readily available for presentation via MHC II molecules, and for this reason this antigen presentation pathway is usually referred to as exogenous. As membrane proteins on their way to degradation in lysosomes as part of their normal turn-over are also contained in the endosomal/lysosomal compartments, exogenous antigens must compete with those derived from endogenous components. Autophagy is also a source of endogenous peptides, autophagosomes constitutively fuse with MHC class II loading compartments. In addition to APCs, other cells of the gastrointestinal tract, such as epithelial cells, express MHC class II molecules and CD74 and act as APCs, which is an unusual trait of the GI tract. To produce a MHC class II molecule that presents an antigen, three MHC class II molecules (heterodimers of an alpha and a beta chain) associate with a CD74 trimer in the ER to form a heterononamer. Soon after the entry of this complex into the endosomal/lysosomal system where antigen processing occurs, CD74 undergoes a sequential degradation by various proteases, including CTSS and CTSL, leaving a small fragment termed CLIP (class-II-associated invariant chain peptide). The removal of CLIP is facilitated by HLA-DM via direct binding to the alpha-beta-CLIP complex so that CLIP is released. HLA-DM stabilizes MHC class II molecules until primary high affinity antigenic peptides are bound. The MHC II molecule bound to a peptide is then transported to the cell membrane surface. In B-cells, the interaction between HLA-DM and MHC class II molecules is regulated by HLA-DO. Primary dendritic cells (DCs) also to express HLA-DO. Lysosomal microenvironment has been implicated in the regulation of antigen loading into MHC II molecules, increased acidification produces increased proteolysis and efficient peptide loading. | *Immunity* | WTD$_{ML}$<br>MD$_{MD}$ |
| P01920 | *HLA-DQB1*<br>*HLA-DQB* | FUNCTION: Binds peptides derived from antigens that access the endocytic route of antigen presenting cells (APC) and presents them on the cell surface for recognition by the CD4 T-cells. The peptide binding cleft accommodates peptides of 10-30 residues. The peptides presented by MHC class II molecules are generated mostly by degradation of proteins that access the endocytic route, where they are processed by lysosomal proteases and other hydrolases. Exogenous antigens that have been endocytosed by the APC are thus readily available for presentation via MHC II molecules, and for this reason this antigen presentation pathway is usually referred to as exogenous. As membrane proteins on their way to degradation in lysosomes as part of their normal turn-over are also contained in the endosomal/lysosomal compartments, exogenous antigens must compete with those derived from endogenous components. Autophagy is also a source of endogenous peptides, autophagosomes constitutively fuse with MHC class II loading compartments. In addition to APCs, other cells of the gastrointestinal tract, such as epithelial cells, express MHC class II molecules and CD74 and act as APCs, which is an unusual trait of the GI tract. To produce a MHC class II molecule that presents an antigen, three MHC class II molecules (heterodimers of an alpha and a beta chain) associate with a CD74 trimer in the ER to form a heterononamer. Soon after the entry of this complex into the endosomal/lysosomal system where antigen processing occurs, CD74 undergoes a sequential degradation by various proteases, including CTSS and CTSL, leaving a small fragment termed CLIP (class-II-associated invariant chain peptide). The removal of CLIP is facilitated by HLA-DM via direct binding to the alpha-beta-CLIP complex so that CLIP is released. HLA-DM stabilizes MHC class II molecules until primary high affinity antigenic peptides are bound. The MHC II molecule bound to a peptide is then transported to the cell membrane surface. In B-cells, the interaction between HLA-DM and MHC class II molecules is regulated by HLA-DO. Primary dendritic cells (DCs) also to express HLA-DO. Lysosomal microenvironment has been implicated in the regulation of antigen loading into MHC II molecules, increased acidification produces increased proteolysis and efficient peptide loading. | *Immunity* | WTD$_{ML}$<br>MD$_{MD}$ |
| P05107 | *ITGB2*<br>*CD18*<br>*MFI7* | FUNCTION: Integrin ITGAL/ITGB2 is a receptor for ICAM1, ICAM2, ICAM3 and ICAM4. Integrin ITGAL/ITGB2 is also a receptor for the secreted form of ubiquitin-like protein ISG15; the interaction is mediated by ITGAL (PubMed:29100055). Integrins ITGAM/ITGB2 and ITGAX/ITGB2 are receptors for the iC3b fragment of the third complement component and for fibrinogen. Integrin ITGAX/ITGB2 recognizes the sequence G-P-R in fibrinogen alpha-chain. Integrin ITGAM/ITGB2 recognizes P1 and P2 peptides of fibrinogen gamma chain. Integrin ITGAM/ITGB2 is also a receptor for factor X. Integrin ITGAD/ITGB2 is a receptor for ICAM3 and VCAM1. Contributes to natural killer cell cytotoxicity (PubMed:15356110). Involved in leukocyte adhesion and transmigration of leukocytes including T-cells and neutrophils (PubMed:11812992, PubMed:28807980). Triggers neutrophil transmigration during lung injury through PTK2B/PYK2-mediated activation (PubMed:18587400). Integrin ITGAL/ITGB2 in association with ICAM3, contributes to apoptotic neutrophil phagocytosis by macrophages (PubMed:23775590). In association with alpha subunit ITGAM/CD11b, required for CD177-PRTN3-mediated activation of TNF primed neutrophils (PubMed:21193407). {ECO:0000269\|PubMed:11812992, ECO:0000269\|PubMed:15356110, ECO:0000269\|PubMed:18587400, ECO:0000269\|PubMed:21193407, ECO:0000269\|PubMed:23775590, ECO:0000269\|PubMed:28807980, ECO:0000269\|PubMed:29100055}. | *Immunity* | WTD$_{ML}$ |
| P13762 | *HLA-DRB4* | FUNCTION: Binds peptides derived from antigens that access the endocytic route of antigen presenting cells (APC) and presents them on the cell surface for recognition by the CD4 T-cells. The peptide binding cleft accommodates peptides of 10-30 residues. The peptides presented by MHC class II molecules are generated mostly by degradation of proteins that access the endocytic route, where they are processed by lysosomal proteases and other hydrolases. Exogenous antigens that have been endocytosed by the APC are thus readily available for presentation via MHC II molecules, and for this reason this antigen presentation pathway is usually referred to as exogenous. As membrane proteins on their way to degradation in lysosomes as part of their normal turn-over are also contained in the endosomal/lysosomal compartments, exogenous antigens must compete with those derived from endogenous components. Autophagy is also a source of endogenous peptides, autophagosomes constitutively fuse with MHC class II loading compartments. In addition to APCs, other cells of the gastrointestinal tract, such as epithelial cells, express MHC class II molecules and CD74 and act as APCs, which is an unusual trait of the GI tract. To produce a MHC class II molecule that presents an antigen, three MHC class II molecules (heterodimers of an alpha and a beta chain) associate with a CD74 trimer in the ER to form a heterononamer. Soon after the entry of this complex into the endosomal/lysosomal system where antigen processing occurs, CD74 undergoes a sequential degradation by various proteases, including CTSS and CTSL, leaving a small fragment termed CLIP (class-II-associated invariant chain peptide). The removal of CLIP is facilitated by HLA-DM via direct binding to the alpha-beta-CLIP complex so that CLIP is released. HLA-DM stabilizes MHC class II molecules until primary high affinity antigenic peptides are bound. The MHC II molecule bound to a peptide is then transported to the cell membrane surface. In B-cells, the interaction between HLA-DM and MHC class II molecules is regulated by HLA-DO. Primary dendritic cells (DCs) also to express HLA-DO. Lysosomal microenvironment has been implicated in the regulation of antigen loading into MHC II molecules, increased acidification produces increased proteolysis and efficient peptide loading. | *Immunity* | WTD$_{KEY}$<br>MD$_{MD}$ |
| P14317 | *HCLS1*<br>*HS1* | FUNCTION: Substrate of the antigen receptor-coupled tyrosine kinase. Plays a role in antigen receptor signaling for both clonal expansion and deletion in lymphoid cells. May also be involved in the regulation of gene expression. | *Immunity* | MD$_{MD}$ |

| | | | | |
|---|---|---|---|---|
| P20701 | *ITGAL*<br>*CD11A* | FUNCTION: Integrin ITGAL/ITGB2 is a receptor for ICAM1, ICAM2, ICAM3 and ICAM4. Integrin ITGAL/ITGB2 is a receptor for F11R (PubMed:11812992, PubMed:15528364). Integrin ITGAL/ITGB2 is a receptor for the secreted form of ubiquitin-like protein ISG15; the interaction is mediated by ITGAL (PubMed:29100055). Involved in a variety of immune phenomena including leukocyte-endothelial cell interaction, cytotoxic T-cell mediated killing, and antibody dependent killing by granulocytes and monocytes. Contributes to natural killer cell cytotoxicity (PubMed:15356110). Involved in leukocyte adhesion and transmigration of leukocytes including T-cells and neutrophils (PubMed:11812992). Required for generation of common lymphoid progenitor cells in bone marrow, indicating a role in lymphopoiesis (By similarity). Integrin ITGAL/ITGB2 in association with ICAM3, contributes to apoptotic neutrophil phagocytosis by macrophages (PubMed:23775590). {ECO:0000250\|UniProtKB:P24063, ECO:0000269\|PubMed:11812992, ECO:0000269\|PubMed:15356110, ECO:0000269\|PubMed:15528364, ECO:0000269\|PubMed:23775590, ECO:0000269\|PubMed:29100055}. | *Immunity* | $WTD_{ML}$ |
| P26022 | *PTX3*<br>*TNFAIP5*<br>*TSG14* | FUNCTION: Plays a role in the regulation of innate resistance to pathogens, inflammatory reactions, possibly clearance of self-components and female fertility. {ECO:0000305\|PubMed:12763682}. | *Immunity* | $WTD_{KEY}$ |
| P31629 | *HIVEP2* | FUNCTION: This protein specifically binds to the DNA sequence 5'-GGGACTTTCC-3' which is found in the enhancer elements of numerous viral promoters such as those of SV40, CMV, or HIV1. In addition, related sequences are found in the enhancer elements of a number of cellular promoters, including those of the class I MHC, interleukin-2 receptor, somatostatin receptor II, and interferon-beta genes. It may act in T-cell activation. | *Immunity* | $MD_{MD}$ |
| P31994 | *FCGR2B*<br>*CD32*<br>*FCG2*<br>*IGFR2* | FUNCTION: Receptor for the Fc region of complexed or aggregated immunoglobulins gamma. Low affinity receptor. Involved in a variety of effector and regulatory functions such as phagocytosis of immune complexes and modulation of antibody production by B-cells. Binding to this receptor results in down-modulation of previous state of cell activation triggered via antigen receptors on B-cells (BCR), T-cells (TCR) or via another Fc receptor. Isoform IIB1 fails to mediate endocytosis or phagocytosis. Isoform IIB2 does not trigger phagocytosis. | *Immunity* | $MD_{MD}$ |
| P32456 | *GBP2* | FUNCTION: Interferon (IFN)-inducible GTPase that plays important roles in innate immunity against a diverse range of bacterial, viral and protozoan pathogens (PubMed:31091448). Hydrolyzes GTP to GMP in 2 consecutive cleavage reactions, but the major reaction product is GDP (PubMed:8706832). Following infection, recruited to the pathogen-containing vacuoles or vacuole-escaped bacteria and acts as a positive regulator of inflammasome assembly by promoting the release of inflammasome ligands from bacteria (By similarity). Acts by promoting lysis of pathogen-containing vacuoles, releasing pathogens into the cytosol (By similarity). Following pathogen release in the cytosol, promotes recruitment of proteins that mediate bacterial cytolysis: this liberates ligands that are detected by inflammasomes, such as lipopolysaccharide (LPS) that activates the non-canonical CASP4/CASP11 inflammasome or double-stranded DNA (dsDNA) that activates the AIM2 inflammasome (By similarity). Confers protection to the protozoan pathogen Toxoplasma gondii (By similarity). Independently of its GTPase activity, acts as an inhibitor of various viruses infectivity, such as HIV-1, Zika and influenza A viruses, by inhibiting FURIN-mediated maturation of viral envelope proteins (PubMed:31091448). {ECO:0000250\|UniProtKB:Q9Z0E6, ECO:0000269\|PubMed:31091448, ECO:0000269\|PubMed:8706832}. | *Immunity* | $WTD_{ML}$ |
| P40305 | *IFI27* | FUNCTION: Probable adapter protein involved in different biological processes (PubMed:22427340, PubMed:27194766). Part of the signaling pathways that lead to apoptosis (PubMed:18330707, PubMed:27673746, PubMed:24970806). Involved in type-I interferon-induced apoptosis characterized by a rapid and robust release of cytochrome C from the mitochondria and activation of BAX and caspases 2, 3, 6, 8 and 9 (PubMed:18330707, PubMed:27673746). Also functions in TNFSF10-induced apoptosis (PubMed:24970806). May also have a function in the nucleus, where it may be involved in the interferon-induced negative regulation of the transcriptional activity of NR4A1, NR4A2 and NR4A3 through the enhancement of XPO1-mediated nuclear export of these nuclear receptors (PubMed:22427340). May thereby play a role in the vascular response to injury (By similarity). In the innate immune response, has an antiviral activity towards hepatitis C virus/HCV (PubMed:27194766, PubMed:27777077). May prevent the replication of the virus by recruiting both the hepatitis C virus non-structural protein 5A/NS5A and the ubiquitination machinery via SKP2, promoting the ubiquitin-mediated proteasomal degradation of NS5A (PubMed:27194766, PubMed:27777077). Promotes also virus-induced pyroptosis by activating CASP3 in the mitochondria after 'Lys-6'-linked ubiquitination by TRIM21 (PubMed:36426955). {ECO:0000250\|UniProtKB:Q8R412, ECO:0000269\|PubMed:18330707, ECO:0000269\|PubMed:22427340, ECO:0000269\|PubMed:24970806, ECO:0000269\|PubMed:27194766, ECO:0000269\|PubMed:27673746, ECO:0000269\|PubMed:27777077, ECO:0000269\|PubMed:36426955}. | *Immunity* | $WTD_{ML}$ |
| P40933 | *IL15* | FUNCTION: Cytokine that plays a major role in the development of inflammatory and protective immune responses to microbial invaders and parasites by modulating immune cells of both the innate and adaptive immune systems (PubMed:15123770). Stimulates the proliferation of natural killer cells, T-cells and B-cells and promotes the secretion of several cytokines (PubMed:8178155, PubMed:9326248). In monocytes, induces the production of IL8 and monocyte chemotactic protein 1/CCL2, two chemokines that attract neutrophils and monocytes respectively to sites of infection (PubMed:9326248). Unlike most cytokines, which are secreted in soluble form, IL15 is expressed in association with its high affinity IL15RA on the surface of IL15-producing cells and delivers signals to target cells that express IL2RB and IL2RG receptor subunits (PubMed:8026467, PubMed:23104097, PubMed:10233906). Binding to its receptor triggers the phosphorylation of JAK1 and JAK3 and the recruitment and subsequent phosphorylation of signal transducer and activator of transcription-3/STAT3 and STAT5 (PubMed:7568001). In mast cells, induces the rapid tyrosine phosphorylation of STAT6 and thereby controls mast cell survival and release of cytokines such as IL4 (By similarity). {ECO:0000250\|UniProtKB:P48346, ECO:0000269\|PubMed:10233906, ECO:0000269\|PubMed:15123770, ECO:0000269\|PubMed:23104097, ECO:0000269\|PubMed:7568001, ECO:0000269\|PubMed:8026467, ECO:0000269\|PubMed:8178155, ECO:0000269\|PubMed:9326248}. | *Immunity* | $MD_{MD}$ |

| | | | | |
|---|---|---|---|---|
| P79483 | *HLA-DRB3* | FUNCTION: A beta chain of antigen-presenting major histocompatibility complex class II (MHCII) molecule. In complex with the alpha chain HLA-DRA, displays antigenic peptides on professional antigen presenting cells (APCs) for recognition by alpha-beta T cell receptor (TCR) on HLA-DRB3-restricted CD4-positive T cells. This guides antigen-specific T-helper effector functions, both antibody-mediated immune response and macrophage activation, to ultimately eliminate the infectious agents and transformed cells. Typically presents extracellular peptide antigens of 10 to 30 amino acids that arise from proteolysis of endocytosed antigens in lysosomes (PubMed:2788702, PubMed:2463305, PubMed:16148104, PubMed:19531622, PubMed:20368442, PubMed:19830726, PubMed:23569328, PubMed:22929521, PubMed:30282837, PubMed:31020640, PubMed:31333679, PubMed:31308093). In the tumor microenvironment, presents antigenic peptides that are primarily generated in tumor-resident APCs likely via phagocytosis of apoptotic tumor cells or macropinocytosis of secreted tumor proteins (By similarity). Presents peptides derived from intracellular proteins that are trapped in autolysosomes after macroautophagy, a mechanism especially relevant for T cell selection in the thymus and central immune tolerance (By similarity). The selection of the immunodominant epitopes follows two processing modes: 'bind first, cut/trim later' for pathogen-derived antigenic peptides and 'cut first, bind later' for autoantigens/self-peptides. The anchor residue at position 1 of the peptide N-terminus, usually a large hydrophobic residue, is essential for high affinity interaction with MHCII molecules (By similarity). {ECO:0000250\|UniProtKB:P01911, ECO:0000269\|PubMed:16148104, ECO:0000269\|PubMed:19531622, ECO:0000269\|PubMed:19830726, ECO:0000269\|PubMed:20368442, ECO:0000269\|PubMed:22929521, ECO:0000269\|PubMed:23569328, ECO:0000269\|PubMed:2463305, ECO:0000269\|PubMed:2788702, ECO:0000269\|PubMed:30282837, ECO:0000269\|PubMed:31020640, ECO:0000269\|PubMed:31308093, ECO:0000269\|PubMed:31333679}.; FUNCTION: ALLELE DRB3*01:01: Exclusively presents several immunogenic epitopes derived from C. tetani neurotoxin tetX, playing a significant role in immune recognition and long-term protection (PubMed:19830726, PubMed:2788702, PubMed:2463305). Presents viral epitopes derived from HHV-6B U11, TRX2/U56 and U85 antigens to polyfunctional CD4-positive T cells with cytotoxic activity implicated in control of HHV-6B infection (PubMed:31020640). {ECO:0000269\|PubMed:19830726, ECO:0000269\|PubMed:2463305, ECO:0000269\|PubMed:2788702, ECO:0000269\|PubMed:31020640}.; FUNCTION: ALLELE DRB3*02:02 Exclusively presents several immunogenic epitopes derived from C. tetani neurotoxin tetX, playing a significant role in immune recognition and long-term protection (PubMed:19830726, PubMed:2788702). Upon EBV infection, presents to CD4-positive T cells latent antigen EBNA2 (PRSPTVFYNIPPMPLPPSQL) and lytic antigen BZLF1 (LTAYHVSTAPTGSWF) peptides, driving oligoclonal expansion and selection of virus-specific memory T cell subsets with cytotoxic potential to directly eliminate virus-infected B cells (PubMed:31308093, PubMed:23569328). Presents viral epitopes derived from HHV-6B U11, gB/U39 and gH/U48 antigens to polyfunctional CD4-positive T cells with cytotoxic activity implicated in control of HHV-6B infection (PubMed:31020640). Plays a minor role in CD4-positive T cell immune response against Dengue virus by presenting conserved peptides from capsid and non-structural NS3 proteins (PubMed:31333679). Displays peptides derived from IAV matrix protein M, implying a role in protection against IAV infection (PubMed:19830726). In the context of tumor immunesurveillance, may present to T-helper 1 cells an immunogenic epitope derived from tumor-associated antigen WT1 (KRYFKLSHLQMHSRKH), likely providing for effective antitumor immunity in a wide range of solid and hematological malignancies (PubMed:22929521). Presents to Vbeta2-positive T-helper 1 cells specifically an immunodominant peptide derived from tumor antigen CTAG1A/NY-ESO-1(PGVLLKEFTVSGNILTIRLTAADHR) and confers protective memory response (PubMed:19531622, PubMed:20368442). In metastatic epithelial tumors, presents to intratumoral CD4-positive T cells a TP53 neoantigen (HYNYMCNSSCMGSMNRRPILTIITL) carrying G245S hotspot driver mutation and may mediate tumor regression (PubMed:30282837). {ECO:0000269\|PubMed:19531622, ECO:0000269\|PubMed:19830726, ECO:0000269\|PubMed:20368442, ECO:0000269\|PubMed:22929521, ECO:0000269\|PubMed:23569328, ECO:0000269\|PubMed:2788702, ECO:0000269\|PubMed:30282837, ECO:0000269\|PubMed:31020640, ECO:0000269\|PubMed:31308093, ECO:0000269\|PubMed:31333679}.; FUNCTION: ALLELE DRB3*03:01: Presents a series of conserved peptides derived from the M. tuberculosis PPE family of proteins, in particular PPE29 and PPE33, known to be highly immunogenic (PubMed:32341563). Presents immunogenic epitopes derived from C. tetani neurotoxin tetX, playing a role in immune recognition and long-term protection (PubMed:2788702). Displays immunodominant viral peptides from HCV non-structural protein NS2, as part of a broad range T-helper response to resolve infection (PubMed:16148104). {ECO:0000269\|PubMed:16148104, ECO:0000269\|PubMed:2788702, ECO:0000269\|PubMed:32341563}. | *Immunity* | MD$_{MD}$ |
| Q01804 | *OTUD4 HIN-1 KIAA1046* | FUNCTION: Deubiquitinase which hydrolyzes the isopeptide bond between the ubiquitin C-terminus and the lysine epsilon-amino group of the target protein (PubMed:23827681, PubMed:25944111, PubMed:29395066). May negatively regulate inflammatory and pathogen recognition signaling in innate immune response. Upon phosphorylation at Ser-202 and Ser-204 residues, via IL-1 receptor and Toll-like receptor signaling pathway, specifically deubiquitinates 'Lys-63'-polyubiquitinated MYD88 adapter protein triggering down-regulation of NF-kappa-B-dependent transcription of inflammatory mediators (PubMed:29395066). Independently of the catalytic activity, acts as a scaffold for alternative deubiquitinases to assemble specific deubiquitinase-substrate complexes. Associates with USP7 and USP9X deubiquitinases to stabilize alkylation repair enzyme ALKBH3, thereby promoting the repair of alkylated DNA lesions (PubMed:25944111). {ECO:0000269\|PubMed:23827681, ECO:0000269\|PubMed:25944111, ECO:0000269\|PubMed:29395066}. | *Immunity* | MD$_{MD}$ |
| Q14684 | *RRP1B KIAA0179* | FUNCTION: Positively regulates DNA damage-induced apoptosis by acting as a transcriptional coactivator of proapoptotic target genes of the transcriptional activator E2F1 (PubMed:20040599). Likely to play a role in ribosome biogenesis by targeting serine/threonine protein phosphatase PP1 to the nucleolus (PubMed:20926688). Involved in regulation of mRNA splicing (By similarity). Inhibits SIPA1 GTPase activity (By similarity). Involved in regulating expression of extracellular matrix genes (By similarity). Associates with chromatin and may play a role in modulating chromatin structure (PubMed:19710015). {ECO:0000250\|UniProtKB:Q91YK2, ECO:0000269\|PubMed:19710015, ECO:0000269\|PubMed:20040599, ECO:0000269\|PubMed:20926688}.; FUNCTION: (Microbial infection) Following influenza A virus (IAV) infection, promotes viral mRNA transcription by facilitating the binding of IAV RNA-directed RNA polymerase to capped mRNA. {ECO:0000269\|PubMed:26311876}. | *Immunity* | MD$_{MD}$ |
| Q6P9F5 | *TRIM40 RNF35* | FUNCTION: E3 ubiquitin-protein ligase that plays a role in the limitation of the innate immune response (PubMed:21474709, PubMed:29117565). Mediates inhibition of the RLR signaling pathway by ubiquitinating RIGI and IFIH1 receptors, leading to their proteasomal degradation (PubMed:21474709). Promotes also the neddylation of IKBKG/NEMO, stabilizing NFKBIA, and thereby inhibiting of NF-kappa-B nuclear translocation and activation (PubMed:21474709). {ECO:0000269\|PubMed:21474709, ECO:0000269\|PubMed:29117565}. | *Immunity* | MD$_{MD}$ |
| Q6UX01 | *LMBR1L KIAA1174 LIMR UNQ458/ PRO783* | FUNCTION: Plays an essential role in lymphocyte development by negatively regulating the canonical Wnt signaling pathway (By similarity). In association with UBAC2 and E3 ubiquitin-protein ligase AMFR, promotes the ubiquitin-mediated degradation of CTNNB1 and Wnt receptors FZD6 and LRP6 (By similarity). LMBR1L stabilizes the beta-catenin destruction complex that is required for regulating CTNNB1 levels (By similarity). Acts as a LCN1 receptor and can mediate its endocytosis (PubMed:11287427, PubMed:12591932, PubMed:23964685). {ECO:0000250\|UniProtKB:Q9D1E5, ECO:0000269\|PubMed:11287427, ECO:0000269\|PubMed:12591932, ECO:0000269\|PubMed:23964685}. | *Immunity* | WTD$_{ML}$ |

| | | | | |
|---|---|---|---|---|
| Q7L513 | *FCRLA*<br>*FCRL*<br>*FCRL1*<br>*FCRLM1*<br>*FCRX*<br>*FREB*<br>*UNQ291/*<br>*PRO329* | FUNCTION: May be implicated in B-cell differentiation and lymphomagenesis. {ECO:0000269\|PubMed:11754007, ECO:0000269\|PubMed:11891275}. | *Immunity* | $MD_{MD}$ |
| Q86WV6 | *STING1*<br>*ERIS*<br>*MITA*<br>*STING*<br>*TMEM17*<br>*3* | FUNCTION: Facilitator of innate immune signaling that acts as a sensor of cytosolic DNA from bacteria and viruses and promotes the production of type I interferon (IFN-alpha and IFN-beta) (PubMed:18724357, PubMed:18818105, PubMed:19433799, PubMed:19776740, PubMed:23027953, PubMed:23910378, PubMed:23747010, PubMed:29973723, PubMed:30842659, PubMed:35045565, PubMed:27801882, PubMed:36808561). Innate immune response is triggered in response to non-CpG double-stranded DNA from viruses and bacteria delivered to the cytoplasm (PubMed:26300263). Acts by binding cyclic dinucleotides: recognizes and binds cyclic di-GMP (c-di-GMP), a second messenger produced by bacteria, and cyclic GMP-AMP (cGAMP), a messenger produced by CGAS in response to DNA virus in the cytosol (PubMed:21947006, PubMed:23258412, PubMed:23707065, PubMed:23722158, PubMed:26229117, PubMed:23910378, PubMed:23747010, PubMed:30842659). Upon binding of c-di-GMP or cGAMP, STING1 oligomerizes, translocates from the endoplasmic reticulum and is phosphorylated by TBK1 on the pLxIS motif, leading to recruitment and subsequent activation of the transcription factor IRF3 to induce expression of type I interferon and exert a potent anti-viral state (PubMed:22394562, PubMed:25636800, PubMed:29973723, PubMed:30842653, PubMed:35045565). In addition to promote the production of type I interferons, plays a direct role in autophagy (PubMed:30568238, PubMed:30842662). Following cGAMP-binding, STING1 buds from the endoplasmic reticulum into COPII vesicles, which then form the endoplasmic reticulum-Golgi intermediate compartment (ERGIC) (PubMed:30842662). The ERGIC serves as the membrane source for WIPI2 recruitment and LC3 lipidation, leading to formation of autophagosomes that target cytosolic DNA or DNA viruses for degradation by the lysosome (PubMed:30842662). The autophagy- and interferon-inducing activities can be uncoupled and autophagy induction is independent of TBK1 phosphorylation (PubMed:30568238, PubMed:30842662). Autophagy is also triggered upon infection by bacteria: following c-di-GMP-binding, which is produced by live Gram-positive bacteria, promotes reticulophagy (By similarity). Exhibits 2',3' phosphodiester linkage-specific ligand recognition: can bind both 2'-3' linked cGAMP (2'-3'-cGAMP) and 3'-3' linked cGAMP but is preferentially activated by 2'-3' linked cGAMP (PubMed:26300263, PubMed:23910378, PubMed:23747010). The preference for 2'-3'-cGAMP, compared to other linkage isomers is probably due to the ligand itself, whichs adopts an organized free-ligand conformation that resembles the STING1-bound conformation and pays low energy costs in changing into the active conformation (PubMed:26150511). May be involved in translocon function, the translocon possibly being able to influence the induction of type I interferons (PubMed:18724357). May be involved in transduction of apoptotic signals via its association with the major histocompatibility complex class II (MHC-II) (By similarity). {ECO:0000250\|UniProtKB:Q3TBT3, ECO:0000269\|PubMed:18724357, ECO:0000269\|PubMed:18818105, ECO:0000269\|PubMed:19433799, ECO:0000269\|PubMed:19776740, ECO:0000269\|PubMed:21947006, ECO:0000269\|PubMed:22394562, ECO:0000269\|PubMed:23027953, ECO:0000269\|PubMed:23258412, ECO:0000269\|PubMed:23707065, ECO:0000269\|PubMed:23722158, ECO:0000269\|PubMed:23747010, ECO:0000269\|PubMed:23910378, ECO:0000269\|PubMed:25636800, ECO:0000269\|PubMed:26150511, ECO:0000269\|PubMed:26229117, ECO:0000269\|PubMed:26300263, ECO:0000269\|PubMed:27801882, ECO:0000269\|PubMed:29973723, ECO:0000269\|PubMed:30568238, ECO:0000269\|PubMed:30842653, ECO:0000269\|PubMed:30842659, ECO:0000269\|PubMed:30842662, ECO:0000269\|PubMed:35045565, ECO:0000269\|PubMed:36808561}.; FUNCTION: (Microbial infection) Antiviral activity is antagonized by oncoproteins, such as papillomavirus (HPV) protein E7 and adenovirus early E1A protein (PubMed:26405230). Such oncoproteins prevent the ability to sense cytosolic DNA (PubMed:26405230). {ECO:0000269\|PubMed:26405230}. | *Immunity* | $WTD_{ML}$ |
| Q8IXQ6 | *PARP9*<br>*BAL BAL1* | FUNCTION: ADP-ribosyltransferase which, in association with E3 ligase DTX3L, plays a role in DNA damage repair and in immune responses including interferon-mediated antiviral defenses (PubMed:16809771, PubMed:23230272, PubMed:26479788, PubMed:27796300). Within the complex, enhances DTX3L E3 ligase activity which is further enhanced by PARP9 binding to poly(ADP-ribose) (PubMed:28525742). In association with DTX3L and in presence of E1 and E2 enzymes, mediates NAD(+)-dependent mono-ADP-ribosylation of ubiquitin which prevents ubiquitin conjugation to substrates such as histones (PubMed:28525742). During DNA repair, PARP1 recruits PARP9/BAL1-DTX3L complex to DNA damage sites via PARP9 binding to ribosylated PARP1 (PubMed:23230272). Subsequent PARP1-dependent PARP9/BAL1-DTX3L-mediated ubiquitination promotes the rapid and specific recruitment of 53BP1/TP53BP1, UIMC1/RAP80, and BRCA1 to DNA damage sites (PubMed:23230272, PubMed:28525742). In response to DNA damage, PARP9-DTX3L complex is required for efficient non-homologous end joining (NHEJ); the complex function is negatively modulated by PARP9 activity (PubMed:28525742). Dispensable for B-cell receptor (BCR) assembly through V(D)J recombination and class switch recombination (CSR) (By similarity). In macrophages, positively regulates pro-inflammatory cytokines production in response to IFNG stimulation by suppressing PARP14-mediated STAT1 ADP-ribosylation and thus promoting STAT1 phosphorylation (PubMed:27796300). Also suppresses PARP14-mediated STAT6 ADP-ribosylation (PubMed:27796300). {ECO:0000250\|UniProtKB:Q8CAS9, ECO:0000269\|PubMed:16809771, ECO:0000269\|PubMed:23230272, ECO:0000269\|PubMed:26479788, ECO:0000269\|PubMed:27796300, ECO:0000269\|PubMed:28525742}. | *Immunity* | $WTD_{ML}$ |

| | | | | |
|---|---|---|---|---|
| Q8NHX9 | *TPCN2 TPC2* | FUNCTION: Intracellular channel initially characterized as a non-selective Ca(2+)-permeable channel activated by NAADP (nicotinic acid adenine dinucleotide phosphate), it is also a highly-selective Na(+) channel activated directly by PI(3,5)P2 (phosphatidylinositol 3,5-bisphosphate) (PubMed:19387438, PubMed:19620632, PubMed:20880839, PubMed:30860481, PubMed:32167471, PubMed:31825310, PubMed:23063126, PubMed:24776928, PubMed:23394946, PubMed:24502975). Localizes to the lysosomal and late endosome membranes where it regulates organellar membrane excitability, membrane trafficking, and pH homeostasis. Is associated with a plethora of physiological processes, including mTOR-dependent nutrient sensing, skin pigmentation and autophagy (PubMed:32167471, PubMed:23394946, PubMed:18488028). Ion selectivity is not fixed but rather agonist-dependent and under defined ionic conditions, can be readily activated by both NAADP and PI(3,5)P2 (PubMed:31825310, PubMed:32167471, PubMed:24502975). As calcium channel, it increases the pH in the lysosomal lumen, as sodium channel, it promotes lysosomal exocytosis (PubMed:31825310, PubMed:32167471). Plays a crucial role in endolysosomal trafficking in the endolysosomal degradation pathway and is potentially involved in the homeostatic control of many macromolecules and cell metabolites (By similarity) (PubMed:18488028, PubMed:19387438, PubMed:19620632, PubMed:20880839, PubMed:23063126, PubMed:23394946, PubMed:24502975, PubMed:24776928, PubMed:31825310, PubMed:32167471, PubMed:32679067). Also expressed in melanosomes of pigmented cells where mediates a Ca(2+) channel and/or PI(3,5)P2-activated melanosomal Na(+) channel to acidify pH and inhibit tyrosinase activity required for melanogenesis and pigmentation (PubMed:27140606). Unlike the voltage-dependent TPCN1, TPCN2 is voltage independent and can be activated solely by PI(3,5)P2 binding. In contrast, PI(4,5)P2, PI(3,4)P2, PI(3)P and PI(5)P have no obvious effect on channel activation (PubMed:30860481). {ECO:0000250\|UniProtKB:Q8BWC0, ECO:0000269\|PubMed:18488028, ECO:0000269\|PubMed:19387438, ECO:0000269\|PubMed:19620632, ECO:0000269\|PubMed:20880839, ECO:0000269\|PubMed:23063126, ECO:0000269\|PubMed:23394946, ECO:0000269\|PubMed:24502975, ECO:0000269\|PubMed:24776928, ECO:0000269\|PubMed:27140606, ECO:0000269\|PubMed:30860481, ECO:0000269\|PubMed:31825310, ECO:0000269\|PubMed:32167471, ECO:0000269\|PubMed:32679067}.; FUNCTION: (Microbial infection) During Ebola virus (EBOV) infection, controls the movement of endosomes containing virus particles and is required by EBOV to escape from the endosomal network into the cell cytoplasm. {ECO:0000269\|PubMed:25722412}.; FUNCTION: (Microbial infection) Required for cell entry of coronaviruses SARS-CoV and SARS-CoV-2, as well as human coronavirus EMC (HCoV-EMC), by endocytosis. {ECO:0000269\|PubMed:32221306}. | *Immunity* | $WTD_{ML}$ |
| Q8TDB6 | *DTX3L BBAP* | FUNCTION: E3 ubiquitin-protein ligase which, in association with ADP-ribosyltransferase PARP9, plays a role in DNA damage repair and in interferon-mediated antiviral responses (PubMed:12670957, PubMed:19818714, PubMed:26479788, PubMed:23230272). Monoubiquitinates several histones, including histone H2A, H2B, H3 and H4 (PubMed:28525742). In response to DNA damage, mediates monoubiquitination of 'Lys-91' of histone H4 (H4K91ub1) (PubMed:19818714). The exact role of H4K91ub1 in DNA damage response is still unclear but it may function as a licensing signal for additional histone H4 post-translational modifications such as H4 'Lys-20' methylation (H4K20me) (PubMed:19818714). PARP1-dependent PARP9-DTX3L-mediated ubiquitination promotes the rapid and specific recruitment of 53BP1/TP53BP1, UIMC1/RAP80, and BRCA1 to DNA damage sites (PubMed:23230272). By monoubiquitinating histone H2B H2BC9/H2BJ and thereby promoting chromatin remodeling, positively regulates STAT1-dependent interferon-stimulated gene transcription and thus STAT1-mediated control of viral replication (PubMed:26479788). Independently of its catalytic activity, promotes the sorting of chemokine receptor CXCR4 from early endosome to lysosome following CXCL12 stimulation by reducing E3 ligase ITCH activity and thus ITCH-mediated ubiquitination of endosomal sorting complex required for transport ESCRT-0 components HGS and STAM (PubMed:24790097). In addition, required for the recruitment of HGS and STAM to early endosomes (PubMed:24790097). In association with PARP9, plays a role in antiviral responses by mediating 'Lys-48'-linked ubiquitination of encephalomyocarditis virus (EMCV) and human rhinovirus (HRV) C3 proteases and thus promoting their proteasomal-mediated degradation (PubMed:26479788). {ECO:0000269\|PubMed:12670957, ECO:0000269\|PubMed:19818714, ECO:0000269\|PubMed:23230272, ECO:0000269\|PubMed:24790097, ECO:0000269\|PubMed:26479788, ECO:0000269\|PubMed:28525742}. | *Immunity* | $WTD_{ML}$ |
| Q8WWU7 | *ITLN2 UNQ2789 /PRO7179* | FUNCTION: May play a role in the defense system against pathogens. {ECO:0000250}. | *Immunity* | $MD_{BTD}$ |
| Q96BN8 | *OTULIN FAM105B* | FUNCTION: Deubiquitinase that specifically removes linear ('Met-1'-linked) polyubiquitin chains to substrates and acts as a regulator of angiogenesis and innate immune response (PubMed:26997266, PubMed:23708998, PubMed:23746843, PubMed:23806334, PubMed:23827681, PubMed:27523608, PubMed:27559085, PubMed:24726323, PubMed:24726327, PubMed:28919039, PubMed:35170849, PubMed:35587511). Required during angiogenesis, craniofacial and neuronal development by regulating the canonical Wnt signaling together with the LUBAC complex (PubMed:23708998). Acts as a negative regulator of NF-kappa-B by regulating the activity of the LUBAC complex (PubMed:23746843, PubMed:23806334). OTULIN function is mainly restricted to homeostasis of the LUBAC complex: acts by removing 'Met-1'-linked autoubiquitination of the LUBAC complex, thereby preventing inactivation of the LUBAC complex (PubMed:26670046). Acts as a key negative regulator of inflammation by restricting spontaneous inflammation and maintaining immune homeostasis (PubMed:27523608). In myeloid cell, required to prevent unwarranted secretion of cytokines leading to inflammation and autoimmunity by restricting linear polyubiquitin formation (PubMed:27523608). Plays a role in innate immune response by restricting linear polyubiquitin formation on LUBAC complex in response to NOD2 stimulation, probably to limit NOD2-dependent pro-inflammatory signaling (PubMed:23806334). {ECO:0000269\|PubMed:23708998, ECO:0000269\|PubMed:23746843, ECO:0000269\|PubMed:23806334, ECO:0000269\|PubMed:23827681, ECO:0000269\|PubMed:24726323, ECO:0000269\|PubMed:24726327, ECO:0000269\|PubMed:26670046, ECO:0000269\|PubMed:26997266, ECO:0000269\|PubMed:27523608, ECO:0000269\|PubMed:27559085, ECO:0000269\|PubMed:28919039, ECO:0000269\|PubMed:35170849, ECO:0000269\|PubMed:35587511}. | *Immunity* | $WTD_{ML}$ |

| | | | | |
|---|---|---|---|---|
| Q96LT7 | *C9orf72 DENND9 DENNL72* | FUNCTION: Component of the C9orf72-SMCR8 complex, a complex that has guanine nucleotide exchange factor (GEF) activity and regulates autophagy (PubMed:27193190, PubMed:27103069, PubMed:27617292, PubMed:28195531, PubMed:32303654). In the complex, C9orf72 and SMCR8 probably constitute the catalytic subunits that promote the exchange of GDP to GTP, converting inactive GDP-bound RAB8A and RAB39B into their active GTP-bound form, thereby promoting autophagosome maturation (PubMed:27103069). The C9orf72-SMCR8 complex also acts as a regulator of autophagy initiation by interacting with the ULK1/ATG1 kinase complex and modulating its protein kinase activity (PubMed:27617292). As part of the C9orf72-SMCR8 complex, stimulates RAB8A and RAB11A GTPase activity in vitro (PubMed:32303654). Positively regulates initiation of autophagy by regulating the RAB1A-dependent trafficking of the ULK1/ATG1 kinase complex to the phagophore which leads to autophagosome formation (PubMed:27334615). Acts as a regulator of mTORC1 signaling by promoting phosphorylation of mTORC1 substrates (PubMed:27559131). Plays a role in endosomal trafficking (PubMed:24549040). May be involved in regulating the maturation of phagosomes to lysosomes (By similarity). Promotes the lysosomal localization and lysosome-mediated degradation of CARM1 which leads to inhibition of starvation-induced lipid metabolism (By similarity). Regulates actin dynamics in motor neurons by inhibiting the GTP-binding activity of ARF6, leading to ARF6 inactivation (PubMed:27723745). This reduces the activity of the LIMK1 and LIMK2 kinases which are responsible for phosphorylation and inactivation of cofilin, leading to CFL1/cofilin activation (PubMed:27723745). Positively regulates axon extension and axon growth cone size in spinal motor neurons (PubMed:27723745). Required for SMCR8 protein expression and localization at pre- and post-synaptic compartments in the forebrain, also regulates protein abundance of RAB3A and GRIA1/GLUR1 in post-synaptic compartments in the forebrain and hippocampus (By similarity). Plays a role within the hematopoietic system in restricting inflammation and the development of autoimmunity (By similarity). {ECO:0000250\|UniProtKB:Q6DFW0, ECO:0000269\|PubMed:24549040, ECO:0000269\|PubMed:27103069, ECO:0000269\|PubMed:27193190, ECO:0000269\|PubMed:27334615, ECO:0000269\|PubMed:27559131, ECO:0000269\|PubMed:27617292, ECO:0000269\|PubMed:27723745, ECO:0000269\|PubMed:28195531, ECO:0000269\|PubMed:32303654}.; FUNCTION: [Isoform 1]: Regulates stress granule assembly in response to cellular stress. {ECO:0000269\|PubMed:27037575}.; FUNCTION: [Isoform 2]: Does not play a role in regulation of stress granule assembly in response to cellular stress. {ECO:0000269\|PubMed:27037575}. | *Immunity* | $WTD_{ML}$ |
| Q96PP9 | *GBP4* | FUNCTION: Interferon (IFN)-inducible GTPase that plays important roles in innate immunity against a diverse range of bacterial, viral and protozoan pathogens (By similarity). Negatively regulates the antiviral response by inhibiting activation of IRF7 transcription factor (By similarity). {ECO:0000250\|UniProtKB:A4UUI3}. | *Immunity* | $WTD_{ML}$ |
| Q9C030 | *TRIM6 RNF89* | FUNCTION: E3 ubiquitin ligase that plays a crucial role in the activation of the IKBKE-dependent branch of the type I interferon signaling pathway (PubMed:24882218, PubMed:31694946). In concert with the ubiquitin-conjugating E2 enzyme UBE2K, synthesizes unanchored 'Lys-48'-linked polyubiquitin chains that promote the oligomerization and autophosphorylation of IKBKE leading to stimulation of an antiviral response (PubMed:24882218). Ubiquitinates also MYC and inhibits its transcription activation activity, maintaining the pluripotency of embryonic stem cells (By similarity). Promotes the association of unanchored 'Lys-48'-polyubiquitin chains with DHX16 leading to enhanced RIGI-mediated innate antiviral immune response (PubMed:35263596). {ECO:0000250\|UniProtKB:Q8BGE7, ECO:0000269\|PubMed:24882218, ECO:0000269\|PubMed:31694946, ECO:0000269\|PubMed:35263596}.; FUNCTION: (Microbial infection) Ubiquitinates ebolavirus protein VP35 leading to enhanced viral transcriptase activity. {ECO:0000269\|PubMed:28679761, ECO:0000269\|PubMed:35533195}. | *Immunity* | $MD_{BTD}$ |
| Q9UDY6 | *TRIM10 RFB30 RNF9* | FUNCTION: E3 ligase that plays an essential role in the differentiation and survival of terminal erythroid cells. May directly bind to PTEN and promote its ubiquitination, resulting in its proteasomal degradation and activation of hypertrophic signaling (By similarity). In addition, plays a role in immune response regulation by repressing the phosphorylation of STAT1 and STAT2 in the interferon/JAK/STAT signaling pathway independent of its E3 ligase activity. Mechanistically, interacts with the intracellular domain of IFNAR1 and thereby inhibits the association between TYK2 and IFNAR1 (PubMed:33811647). {ECO:0000250\|UniProtKB:Q9WUH5, ECO:0000269\|PubMed:33811647}. | *Immunity* | $MD_{MD}$ |
| O60911 | *CTSV CATL2 CTSL2 CTSU UNQ268/ PRO305* | FUNCTION: Cysteine protease. May have an important role in corneal physiology. {ECO:0000269\|PubMed:10029531, ECO:0000269\|PubMed:9727401}. | *Immunity, physiology* | $MD_{MD}$ |
| Q8IW41 | *MAPKAP K5 PRAK* | FUNCTION: Tumor suppressor serine/threonine-protein kinase involved in mTORC1 signaling and post-transcriptional regulation. Phosphorylates FOXO3, ERK3/MAPK6, ERK4/MAPK4, HSP27/HSPB1, p53/TP53 and RHEB. Acts as a tumor suppressor by mediating Ras-induced senescence and phosphorylating p53/TP53. Involved in post-transcriptional regulation of MYC by mediating phosphorylation of FOXO3: phosphorylation of FOXO3 leads to promote nuclear localization of FOXO3, enabling expression of miR-34b and miR-34c, 2 post-transcriptional regulators of MYC that bind to the 3'UTR of MYC transcript and prevent MYC translation. Acts as a negative regulator of mTORC1 signaling by mediating phosphorylation and inhibition of RHEB. Part of the atypical MAPK signaling via its interaction with ERK3/MAPK6 or ERK4/MAPK4: the precise role of the complex formed with ERK3/MAPK6 or ERK4/MAPK4 is still unclear, but the complex follows a complex set of phosphorylation events: upon interaction with atypical MAPK (ERK3/MAPK6 or ERK4/MAPK4), ERK3/MAPK6 (or ERK4/MAPK4) is phosphorylated and then mediates phosphorylation and activation of MAPKAPK5, which in turn phosphorylates ERK3/MAPK6 (or ERK4/MAPK4). Mediates phosphorylation of HSP27/HSPB1 in response to PKA/PRKACA stimulation, inducing F-actin rearrangement. {ECO:0000269\|PubMed:17254968, ECO:0000269\|PubMed:17728103, ECO:0000269\|PubMed:19166925, ECO:0000269\|PubMed:21329882, ECO:0000269\|PubMed:9628874}. | *Immunity, physiology* | $WTD_{ML}$ |
| Q9NQZ5 | *STARD7 GTT1* | FUNCTION: May play a protective role in mucosal tissues by preventing exaggerated allergic responses. {ECO:0000250\|UniProtKB:Q8R1R3}. | *Immunity, physiology* | $WTD_{ML}$ |
| Q16553 | *LY6E 9804 RIGE SCA2 TSA1* | FUNCTION: GPI-anchored cell surface protein that regulates T-lymphocytes proliferation, differentiation, and activation. Regulates the T-cell receptor (TCR) signaling by interacting with component CD3Z/CD247 at the plasma membrane, leading to CD3Z/CD247 phosphorylation modulation (By similarity). Restricts the entry of human coronaviruses, including SARS-CoV, MERS-CoV and SARS-CoV-2, by interfering with spike protein-mediated membrane fusion (PubMed:32641482). Also plays an essential role in placenta formation by acting as the main receptor for syncytin-A (SynA). Therefore, participates in the normal fusion of syncytiotrophoblast layer I (SynT-I) and in the proper morphogenesis of both fetal and maternal vasculatures within the placenta. May also act as a modulator of nicotinic acetylcholine receptors (nAChRs) activity (By similarity). {ECO:0000250\|UniProtKB:Q64253, ECO:0000269\|PubMed:32641482}.; FUNCTION: (Microbial infection) Promotes entry, likely through an enhanced virus-cell fusion process, of various viruses including HIV-1, West Nile virus, dengue virus and Zika virus (PubMed:28130445). In contrast, the paramyxovirus PIV5, which enters at the plasma membrane, does not require LY6E (PubMed:28130445, PubMed:29610346). Mechanistically, adopts a microtubule-like organization upon viral infection and enhances viral uncoating after endosomal escape (PubMed:28130445, PubMed:30190477). {ECO:0000269\|PubMed:28130445, ECO:0000269\|PubMed:29610346, ECO:0000269\|PubMed:30190477}. | *Immunity, reproduction, development* | $WTD_{ML}$ |

| | | | | |
|---|---|---|---|---|
| P06858 | *LPL LIPD* | FUNCTION: Key enzyme in triglyceride metabolism. Catalyzes the hydrolysis of triglycerides from circulating chylomicrons and very low density lipoproteins (VLDL), and thereby plays an important role in lipid clearance from the blood stream, lipid utilization and storage (PubMed:8675619, PubMed:11342582, PubMed:27578112). Although i t has both phospholipase and triglyceride lipase activities it is primarily a triglyceride lipase with low but detectable phospholipase activity (PubMed:7592706, PubMed:12032167). Mediates margination of triglyceride-rich lipoprotein particles in capillaries (PubMed:24726386). Recruited to its site of action on the luminal surface of vascular endothelium by binding to GPIHBP1 and cell surface heparan sulfate proteoglycans (PubMed:11342582, PubMed:27811232). {ECO:0000269\|PubMed:11342582, ECO:0000269\|PubMed:12032167, ECO:0000269\|PubMed:24726386, ECO:0000269\|PubMed:27578112, ECO:0000269\|PubMed:27811232, ECO:0000269\|PubMed:7592706, ECO:0000269\|PubMed:8675619}. | *Metabolism* | $WTD_{KEY}$ |
| P51857 | *AKR1D1 SRD5B1* | FUNCTION: Catalyzes the stereospecific NADPH-dependent reduction of the C4-C5 double bond of bile acid intermediates and steroid hormones carrying a delta(4)-3-one structure to yield an A/B cis-ring junction. This cis-configuration is crucial for bile acid biosynthesis and plays important roles in steroid metabolism. Capable of reducing a broad range of delta-(4)-3-ketosteroids from C18 (such as, 17beta-hydroxyestr-4-en-3-one) to C27 (such as, 7alpha-hydroxycholest-4-en-3-one). {ECO:0000269\|PubMed:11342103, ECO:0000269\|PubMed:18407998, ECO:0000269\|PubMed:20522910, ECO:0000269\|PubMed:21255593, ECO:0000269\|PubMed:7508385}. | *Metabolism* | $WTD_{ML}$ |
| Q02318 | *CYP27A1 CYP27* | FUNCTION: Cytochrome P450 monooxygenase that catalyzes regio- and stereospecific hydroxylation of cholesterol and its derivatives. Hydroxylates (with R stereochemistry) the terminal methyl group of cholesterol side-chain in a three step reaction to yield at first a C26 alcohol, then a C26 aldehyde and finally a C26 acid (PubMed:9660774, PubMed:12077124, PubMed:21411718, PubMed:28190002). Regulates cholesterol homeostasis by catalyzing the conversion of excess cholesterol to bile acids via both the 'neutral' (classic) and the 'acid' (alternative) pathways (PubMed:9660774, PubMed:1708392, PubMed:11412116, PubMed:2019602, PubMed:7915755, PubMed:9186905, PubMed:9790667). May also regulate cholesterol homeostasis via generation of active oxysterols, which act as ligands for NR1H2 and NR1H3 nuclear receptors, modulating the transcription of genes involved in lipid metabolism (PubMed:9660774, PubMed:12077124). Plays a role in cholestanol metabolism in the cerebellum. Similarly to cholesterol, hydroxylates cholestanol and may facilitate sterol diffusion through the blood-brain barrier to the systemic circulation for further degradation (PubMed:28190002). Also hydroxylates retinal 7-ketocholesterol, a noxious oxysterol with pro-inflammatory and pro-apoptotic effects, and may play a role in its elimination from the retinal pigment epithelium (PubMed:21411718). May play a redundant role in vitamin D biosynthesis. Catalyzes 25-hydroxylation of vitamin D3 that is required for its conversion to a functionally active form (PubMed:15465040). {ECO:0000269\|PubMed:11412116, ECO:0000269\|PubMed:12077124, ECO:0000269\|PubMed:15465040, ECO:0000269\|PubMed:1708392, ECO:0000269\|PubMed:2019602, ECO:0000269\|PubMed:21411718, ECO:0000269\|PubMed:28190002, ECO:0000269\|PubMed:7915755, ECO:0000269\|PubMed:9186905, ECO:0000269\|PubMed:9660774, ECO:0000269\|PubMed:9790667}. | *Metabolism* | $WTD_{ML}$ |
| Q49AA0 | *ZFP69 ZNF642* | FUNCTION: Putative transcription factor that appears to regulate lipid metabolism. {ECO:0000250\|UniProtKB:A2A761}. | *Metabolism* | $WTD_{ML}$ $MD_{MD}$ |
| Q8N118 | *CYP4X1 UNQ1929 /PRO440 4* | FUNCTION: A cytochrome P450 monooxygenase that selectively catalyzes the epoxidation of the last double bond of the arachidonoyl moiety of anandamide, potentially modulating endocannabinoid signaling. Has no hydroxylase activity toward various fatty acids, steroids and prostaglandins. Mechanistically, uses molecular oxygen inse rting one oxygen atom into a substrate, and reducing the second into a water molecule, with two electrons provided by NADPH via cytochrome P450 reductase (CPR; NADPH-ferrihemoprotein reductase). {ECO:0000269\|PubMed:18549450}. | *Metabolism* | $WTD_{ML}$ |
| Q9BV23 | *ABHD6* | FUNCTION: Lipase that preferentially hydrolysis medium-chain saturated monoacylglycerols including 2-arachidonoylglycerol (PubMed:22969151). Through 2-arachidonoylglycerol degradation may regulate endocannabinoid signaling pathways (By similarity). Also has a lysophosphatidyl lipase activity with a preference for lysophosphatidylglycerol among other lysophospholipids (By similarity). Also able to degrade bis(monoacylglycero)phosphate (BMP) and constitutes the major enzyme for BMP catabolism (PubMed:26491015). BMP, also known as lysobisphosphatidic acid, is enriched in late endosomes and lysosomes and plays a key role in the formation of intraluminal vesicles and in lipid sorting (PubMed:26491015). {ECO:0000250\|UniProtKB:Q8R2Y0, ECO:0000269\|PubMed:22969151, ECO:0000269\|PubMed:26491015}. | *Metabolism* | $WTD_{ML}$ |
| Q9H0X9 | *OSBPL5 KIAA1534 OBPH1 ORP5* | FUNCTION: Lipid transporter involved in lipid countertransport between the endoplasmic reticulum and the plasma membrane: specifically exchanges phosphatidylserine with phosphatidylinositol 4-phosphate (PI4P), delivering phosphatidylserine to the plasma membrane in exchange for PI4P, which is degraded by the SAC1/SACM1L phosphatase in the endoplasmic reticulum. Binds phosphatidylserine and PI4P in a mutually exclusive manner (PubMed:23934110, PubMed:26206935). May cooperate with NPC1 to mediate the exit of cholesterol from endosomes/lysosomes (PubMed:21220512). Binds 25-hydroxycholesterol and cholesterol (PubMed:17428193). {ECO:0000269\|PubMed:17428193, ECO:0000269\|PubMed:21220512, ECO:0000269\|PubMed:23934110, ECO:0000269\|PubMed:26206935}. | *Metabolism* | $MD_{MD}$ |
| O94933 | *SLITRK3 KIAA0848* | FUNCTION: Suppresses neurite outgrowth. {ECO:0000250}. | *Physiology* | $WTD_{ML}$ |
| P21439 | *ABCB4 MDR3 PGY3* | FUNCTION: [Isoform 1]: Energy-dependent phospholipid efflux translocator that acts as a positive regulator of biliary lipid secretion. Functions as a floppase that translocates specifically phosphatidylcholine (PC) from the inner to the outer leaflet of the canalicular membrane bilayer into the canaliculi of hepatocytes. Translocation of PC makes the biliary phospholipids available for extraction into the canaliculi lumen by bile salt mixed micelles and therefore protects the biliary tree from the detergent activity of bile salts (PubMed:7957936, PubMed:8898203, PubMed:9366571, PubMed:17523162, PubMed:23468132, PubMed:24806754, PubMed:24723470, PubMed:24594635, PubMed:21820390, PubMed:31873305). Plays a role in the recruitment of phosphatidylcholine (PC), phosphatidylethanolamine (PE) and sphingomyelin (SM) molecules to nonraft membranes and to further enrichment of SM and cholesterol in raft membranes in hepatocytes (PubMed:23468132). Required for proper phospholipid bile formation (By similarity). Indirectly involved in cholesterol efflux activity from hepatocytes into the canalicular lumen in the presence of bile salts in an ATP-dependent manner (PubMed:24045840). Promotes biliary phospholipid secretion as canaliculi-containing vesicles from the canalicular plasma membrane (PubMed:9366571, PubMed:28012258). In cooperation with ATP8B1, functions to protect hepatocytes from the deleterious detergent activity of bile salts (PubMed:21820390). Does not confer multidrug resistance (By similarity). {ECO:0000250\|UniProtKB:P21440, ECO:0000269\|PubMed:17523162, ECO:0000269\|PubMed:21820390, ECO:0000269\|PubMed:23468132, ECO:0000269\|PubMed:24045840, ECO:0000269\|PubMed:24594635, ECO:0000269\|PubMed:24723470, ECO:0000269\|PubMed:24806754, ECO:0000269\|PubMed:28012258, ECO:0000269\|PubMed:31873305, ECO:0000269\|PubMed:7957936, ECO:0000269\|PubMed:8898203, ECO:0000269\|PubMed:9366571}. | *Physiology* | $MD_{MD}$ |
| P32238 | *CCKAR CCKRA* | FUNCTION: Receptor for cholecystokinin. Mediates pancreatic growth and enzyme secretion, smooth muscle contraction of the gall bladder and stomach. Has a 1000-fold higher affinity for CCK rather than for gastrin. It modulates feeding and dopamine-induced behavior in the central and peripheral nervous system. This receptor mediates its action by association with G proteins that activate a phosphatidylinositol-calcium second messenger system. | *Physiology* | $WTD_{ML}$ |
| Q14916 | *SLC17A1 NPT1* | FUNCTION: Important for the resorption of phosphate by the kidney (PubMed:7826357). May be involved in actively transporting phosphate into cells via Na(+) cotransport in the renal brush border membrane (PubMed:7826357). Plays a role in urate transport in the kidney (PubMed:27906618, PubMed:25252215). {ECO:0000269\|PubMed:25252215, ECO:0000269\|PubMed:27906618, ECO:0000269\|PubMed:7826357}. | *Physiology* | $WTD_{ML}$ |

| | | | | |
|---|---|---|---|---|
| Q8IUK8 | *CBLN2 UNQ1892/PRO4338* | FUNCTION: Acts as a synaptic organizer in specific subsets of neurons in the brain (By similarity). Essential for long-term maintenance but not establishment of excitatory synapses (By similarity). {ECO:0000250\|UniProtKB:Q8BGU2}. | *Physiology* | $MD_{BTD}$ |
| Q8NB66 | *UNC13C* | FUNCTION: May play a role in vesicle maturation during exocytosis as a target of the diacylglycerol second messenger pathway. May be involved in the regulation of synaptic transmission at parallel fiber - Purkinje cell synapses (By similarity). {ECO:0000250}. | *Physiology* | $MD_{MD}$ |
| Q92954 | *PRG4 MSF SZP* | FUNCTION: Plays a role in boundary lubrication within articulating joints. Prevents protein deposition onto cartilage from synovial fluid by controlling adhesion-dependent synovial growth and inhibiting the adhesion of synovial cells to the cartilage surface.; FUNCTION: Isoform F plays a role as a growth factor acting on the primitive cells of both hematopoietic and endothelial cell lineages. | *Physiology* | $MD_{MD}$ |
| Q9H195 | *MUC3B* | FUNCTION: Major glycoprotein component of a variety of mucus gels. Thought to provide a protective, lubricating barrier against particles and infectious agents at mucosal surfaces (By similarity). {ECO:0000250}. | *Physiology* | $WTD_{ML}$ |
| Q9UBS5 | *GABBR1 GPRC3A* | FUNCTION: Component of a heterodimeric G-protein coupled receptor for GABA, formed by GABBR1 and GABBR2 (PubMed:9872316, PubMed:9872744, PubMed:15617512, PubMed:18165688, PubMed:22660477, PubMed:24305054). Within the heterodimeric GABA receptor, only GABBR1 seems to bind agonists, while GABBR2 mediates coupling to G proteins (PubMed:18165688). Ligand binding causes a conformation change that triggers signaling via guanine nucleotide-binding proteins (G proteins) and modulates the activity of down-stream effectors, such as adenylate cyclase (PubMed:10906333, PubMed:10773016, PubMed:10075644, PubMed:9872744, PubMed:24305054). Signaling inhibits adenylate cyclase, stimulates phospholipase A2, activates potassium channels, inactivates voltage-dependent calcium-channels and modulates inositol phospholipid hydrolysis (PubMed:10075644). Calcium is required for high affinity binding to GABA (By similarity). Plays a critical role in the fine-tuning of inhibitory synaptic transmission (PubMed:9844003). Pre-synaptic GABA receptor inhibits neurotransmitter release by down-regulating high-voltage activated calcium channels, whereas postsynaptic GABA receptor decreases neuronal excitability by activating a prominent inwardly rectifying potassium (Kir) conductance that underlies the late inhibitory postsynaptic potentials (PubMed:9844003, PubMed:9872316, PubMed:10075644, PubMed:9872744, PubMed:22660477). Not only implicated in synaptic inhibition but also in hippocampal long-term potentiation, slow wave sleep, muscle relaxation and antinociception (Probable). Activated by (-)-baclofen, cgp27492 and blocked by phaclofen (PubMed:9844003, PubMed:9872316, PubMed:24305054). {ECO:0000250\|UniProtKB:Q9Z0U4, ECO:0000269\|PubMed:10075644, ECO:0000269\|PubMed:10773016, ECO:0000269\|PubMed:10906333, ECO:0000269\|PubMed:15617512, ECO:0000269\|PubMed:18165688, ECO:0000269\|PubMed:22660477, ECO:0000269\|PubMed:24305054, ECO:0000269\|PubMed:9844003, ECO:0000269\|PubMed:9872316, ECO:0000269\|PubMed:9872744, ECO:0000305}.; FUNCTION: Isoform 1E may regulate the formation of functional GABBR1/GABBR2 heterodimers by competing for GABBR2 binding. This could explain the observation that certain small molecule ligands exhibit differential affinity for central versus peripheral sites. | *Physiology* | $WTD_{ML}$ |
| Q9BT56 | *SPX C12orf39* | FUNCTION: Plays a role as a central modulator of cardiovascular and renal function and nociception. Also plays a role in energy metabolism and storage. Inhibits adrenocortical cell proliferation with minor stimulation on corticosteroid release (By similarity). {ECO:0000250}.; FUNCTION: [Spexin-1]: Acts as a ligand for galanin receptors GALR2 and GALR3 (PubMed:17284679, PubMed:24517231). Intracerebroventricular administration of the peptide induces an increase in arterial blood pressure, a decrease in both heart rate and renal excretion and delayed natriuresis. Intraventricular administration of the peptide induces antinociceptive activity. Also induces contraction of muscarinic-like stomach smooth muscles. Intraperitoneal administration of the peptide induces a reduction in food consumption and body weight. Inhibits long chain fatty acid uptake into adipocytes (By similarity). {ECO:0000250, ECO:0000269\|PubMed:17284679, ECO:0000269\|PubMed:24517231}.; FUNCTION: [Spexin-2]: Intracerebroventricular administration of the peptide induces a decrease in heart rate, but no change in arterial pressure, and an increase in urine flow rate. Intraventricular administration of the peptide induces antinociceptive activity (By similarity). {ECO:0000250}. | *Physiology, metabolism* | $MD_{MD}$ |
| P07686 | *HEXB HCC7* | FUNCTION: Hydrolyzes the non-reducing end N-acetyl-D-hexosamine and/or sulfated N-acetyl-D-hexosamine of glycoconjugates, such as the oligosaccharide moieties from proteins and neutral glycolipids, or from certain mucopolysaccharides (PubMed:11707436, PubMed:9694901, PubMed:8672428, PubMed:8123671). The isozyme B does not hydrolyze each of these substrates, however hydrolyzes efficiently neutral oligosaccharide (PubMed:11707436). Only the isozyme A is responsible for the degradation of GM2 gangliosides in the presence of GM2A (PubMed:9694901, PubMed:8672428, PubMed:8123671). During fertilization is responsible, at least in part, for the zona block to polyspermy. Present in the cortical granules of non-activated oocytes, is exocytosed during the cortical reaction in response to oocyte activation and inactivates the sperm galactosyltransferase-binding site, accounting for the block in sperm binding to the zona pellucida (By similarity). {ECO:0000250\|UniProtKB:P20060, ECO:0000269\|PubMed:11707436, ECO:0000269\|PubMed:8123671, ECO:0000269\|PubMed:8672428, ECO:0000269\|PubMed:9694901}. | *Reproduction* | $MD_{MD}$ |
| P48378 | *RFX2* | FUNCTION: Transcription factor that acts as a key regulator of spermatogenesis. Acts by regulating expression of genes required for the haploid phase during spermiogenesis, such as genes required for cilium assembly and function (By similarity). Recognizes and binds the X-box, a regulatory motif with DNA sequence 5'-GTNRCC(0-3N)RGYAAC-3' present on promoters (PubMed:10330134). Probably activates transcription of the testis-specific histone gene H1-6 (By similarity). {ECO:0000250\|UniProtKB:P48379, ECO:0000269\|PubMed:10330134}. | *Reproduction* | $WTD_{ML}$ $MD_{MD}$ |
| Q07617 | *SPAG1* | FUNCTION: May play a role in the cytoplasmic assembly of the ciliary dynein arms (By similarity). May play a role in fertilization. Binds GTP and has GTPase activity. {ECO:0000250, ECO:0000269\|PubMed:11517287, ECO:0000269\|PubMed:1299558}. | *Reproduction* | $WTD_{ML}$ |
| Q5BJF6 | *ODF2* | FUNCTION: Seems to be a major component of sperm tail outer dense fibers (ODF). ODFs are filamentous structures located on the outside of the axoneme in the midpiece and principal piece of the mammalian sperm tail and may help to maintain the passive elastic structures and elastic recoil of the sperm tail. May have a modulating influence on sperm motility. Functions as a general scaffold protein that is specifically localized at the distal/subdistal appendages of mother centrioles. Component of the centrosome matrix required for the localization of PLK1 and NIN to the centrosomes. Required for the formation and/or maintenance of normal CETN1 assembly. {ECO:0000269\|PubMed:16966375}. | *Reproduction* | $MD_{MD}$ |
| Q6GV28 | *TMEM225 PMP22CD* | FUNCTION: Probably inhibits protein phosphatase 1 (PP1) in sperm via binding to catalytic subunit PPP1CC. {ECO:0000250\|UniProtKB:Q9D9S2}. | *Reproduction* | $WTD_{ML}$ |
| Q6HA08 | *ASTL* | FUNCTION: Oocyte-specific oolemmal receptor involved in sperm and egg adhesion and fertilization. Plays a role in the polyspermy inhibition. Probably acts as a protease for the post-fertilization cleavage of ZP2. Cleaves the sperm-binding ZP2 at the surface of the zona pellucida after fertilization and cortical granule exocytosis, rendering the zona pellucida unable to support further sperm binding. {ECO:0000250\|UniProtKB:Q6HA09}. | *Reproduction* | $WTD_{ML}$ |
| Q8N6M8 | *IQCF1* | FUNCTION: Involved in sperm capacitation and acrosome reaction. {ECO:0000250\|UniProtKB:Q9D9K8}. | *Reproduction* | $MD_{BTD}$ |

| | | | | |
|---|---|---|---|---|
| Q9UBK7 | *RABL2A* | FUNCTION: Plays an essential role in male fertility, sperm intra-flagellar transport, and tail assembly. Binds, in a GTP-regulated manner, to a specific set of effector proteins including key proteins involved in cilia development and function and delivers them into the growing sperm tail. {ECO:0000250\|UniProtKB:E9Q9D5}. | *Reproduction* | WTD$_{ML}$ |
| Q9UKJ8 | *ADAM21* | FUNCTION: May be involved in sperm maturation and/or fertilization. May also be involved in epithelia functions associated with establishing and maintaining gradients of ions or nutrients. | *Reproduction* | WTD$_{ML}$ |
| O43323 | *DHH* | FUNCTION: [Desert hedgehog protein]: The C-terminal part of the desert hedgehog protein precursor displays an autoproteolysis and a cholesterol transferase activity (By similarity). Both activities result in the cleavage of the full-length protein into two parts (N-product and C-product) followed by the covalent attachment of a cholesterol moiety to the C-terminal of the newly generated N-product (By similarity). Both activities occur in the reticulum endoplasmic (By similarity). Functions in cell-cell mediated juxtacrine signaling (PubMed:24342078). Promotes endothelium integrity (PubMed:33063110). Binds to PTCH1 receptor, which functions in association with smoothened (SMO), to activate the transcription of target genes in endothelial cells (PubMed:33063110). In Schwann cells, controls the development of the peripheral nerve sheath and the transition of mesenchymal cells to form the epithelium-like structure of the perineurial tube (By similarity). {ECO:0000250\|UniProtKB:Q61488, ECO:0000250\|UniProtKB:Q62226, ECO:0000269\|PubMed:24342078, ECO:0000269\|PubMed:33063110}.; FUNCTION: [Desert hedgehog protein N-product]: The dually lipidated desert hedgehog protein N-product is essential for a variety of patterning events during development (By similarity). Binds to the patched (PTCH1) receptor, which functions in association with smoothened (SMO), to activate the transcription of target genes (PubMed:11472839, PubMed:33063110). Required for normal testis development and spermatogenesis, namely for the formation of adult-type Leydig cells and normal development of peritubular cells and seminiferous tubules (By similarity). Activates primary cilia signaling on neighboring valve interstitial cells through a paracrine mechanism (By similarity). May induce motor neurons in lateral neural tube and may have a polarizing activity (PubMed:11472839). Prevents the desert hedgehog protein precursor binding to PTCH1 (PubMed:33063110). {ECO:0000250\|UniProtKB:Q15465, ECO:0000250\|UniProtKB:Q61488, ECO:0000250\|UniProtKB:Q62226, ECO:0000269\|PubMed:11472839, ECO:0000269\|PubMed:33063110}. | *Reproduction development* | WTD$_{ML}$ |
| O75752 | *B3GALNT1 B3GALT3 UNQ531/ PRO1074* | FUNCTION: Transfers N-acetylgalactosamine onto globotriaosylceramide (PubMed:10993897). Plays a critical role in preimplantation stage embryonic development (By similarity). {ECO:0000250\|UniProtKB:Q920V1, ECO:0000269\|PubMed:10993897}. | *Reproduction development* | MD$_{BTD}$ |
| P0CW00 | *TSPY8* | FUNCTION: May be involved in sperm differentiation and proliferation. {ECO:0000250\|UniProtKB:Q01534}. | *Reproduction development* | MD$_{MD}$ |
| P83110 | *HTRA3 PRSP* | FUNCTION: Serine protease that cleaves beta-casein/CSN2 as well as several extracellular matrix (ECM) proteoglycans such as decorin/DCN, biglycan/BGN and fibronectin/FN1. Inhibits signaling mediated by TGF-beta family proteins possibly indirectly by degradation of these ECM proteoglycans (By similarity). May act as a tumor suppressor. Negatively regulates, in vitro, trophoblast invasion during placental development and may be involved in the development of the placenta in vivo. May also have a role in ovarian development, granulosa cell differentiation and luteinization (PubMed:21321049, PubMed:22229724). {ECO:0000250\|UniProtKB:Q9D236, ECO:0000269\|PubMed:21321049, ECO:0000269\|PubMed:22229724}. | *Reproduction development* | WTD$_{KEY}$ |
| Q9NX45 | *SOHLH2 TEB1* | FUNCTION: Transcription regulator of both male and female germline differentiation. Suppresses genes involved in spermatogonial stem cells maintenance, and induces genes important for spermatogonial differentiation. Coordinates oocyte differentiation without affecting meiosis I (By similarity). {ECO:0000250\|UniProtKB:Q6IUP1, ECO:0000250\|UniProtKB:Q9D489}. | *Reproduction development* | MD$_{BTD}$ |
| O14804 | *TAAR5 PNR* | FUNCTION: Olfactory receptor specific for trimethylamine, a trace amine. Also activated at lower level by dimethylethylamine. Trimethylamine is a bacterial metabolite found in some animal odors, and to humans it is a repulsive odor associated with bad breath and spoiled food. This receptor is probably mediated by the G(s)-class of G-proteins which activate adenylate cyclase. {ECO:0000269\|PubMed:23393561}. | *Sensory perception* | MD$_{MD}$ |
| Q13002 | *GRIK2 GLUR6* | FUNCTION: Ionotropic glutamate receptor. L-glutamate acts as an excitatory neurotransmitter at many synapses in the central nervous system. Binding of the excitatory neurotransmitter L-glutamate induces a conformation change, leading to the opening of the cation channel, and thereby converts the chemical signal to an electrical impulse. The receptor then desensitizes rapidly and enters a transient inactive state, characterized by the presence of bound agonist (PubMed:28180184). Modulates cell surface expression of NETO2 (By similarity). {ECO:0000250\|UniProtKB:P39087, ECO:0000269\|PubMed:28180184}.; FUNCTION: Independent of its ionotropic glutamate receptor activity, acts as a thermoreceptor conferring sensitivity to cold temperatures (PubMed:31474366). Functions in dorsal root ganglion neurons (By similarity). {ECO:0000250\|UniProtKB:P39087, ECO:0000269\|PubMed:31474366}. | *Sensory perception* | MD$_{MD}$ |
| Q8NGL1 | *OR5D18* | FUNCTION: Odorant receptor. {ECO:0000305}. | *Sensory perception* | WTD$_{ML}$ MD$_{MD}$ |
| Q96RI8 | *TAAR6 TA4 TAR4 TRAR4* | FUNCTION: Orphan receptor. Could be a receptor for trace amines. Trace amines are biogenic amines present in very low levels in mammalian tissues. Although some trace amines have clearly defined roles as neurotransmitters in invertebrates, the extent to which they function as true neurotransmitters in vertebrates has remained speculative. Trace amines are likely to be involved in a variety of physiological functions that have yet to be fully understood. | *Sensory perception* | MD$_{MD}$ |
| Q9H343 | *OR51I1* | FUNCTION: Odorant receptor. {ECO:0000305}. | *Sensory perception* | MD$_{MD}$ |
| Q7Z2W7 | *TRPM8 LTRPC6 TRPP8* | FUNCTION: Receptor-activated non-selective cation channel involved in detection of sensations such as coolness, by being activated by cold temperature below 25 degrees Celsius. Activated by icilin, eucalyptol, menthol, cold and modulation of intracellular pH. Involved in menthol sensation. Permeable for monovalent cations sodium, potassium, and cesium and divalent cation calcium. Temperature sensing is tightly linked to voltage-dependent gating. Activated upon depolarization, changes in temperature resulting in graded shifts of its voltage-dependent activation curves. The chemical agonist menthol functions as a gating modifier, shifting activation curves towards physiological membrane potentials. Temperature sensitivity arises from a tenfold difference in the activation energies associated with voltage-dependent opening and closing. In prostate cancer cells, shows strong inward rectification and high calcium selectivity in contrast to its behavior in normal cells which is characterized by outward rectification and poor cationic selectivity. Plays a role in prostate cancer cell migration (PubMed:25559186). Isoform 2 and isoform 3 negatively regulate menthol- and cold-induced channel activity by stabilizing the closed state of the channel. {ECO:0000269\|PubMed:15306801, ECO:0000269\|PubMed:16174775, ECO:0000269\|PubMed:22128173, ECO:0000269\|PubMed:25559186}. | *Sensory perception, physiology* | WTD$_{ML}$ |

| | | | | |
|---|---|---|---|---|
| A6NP61 | *ZAR1L* | FUNCTION: mRNA-binding protein required for maternal mRNA storage, translation and degradation during oocyte maturation (By similarity). Probably promotes formation of some phase-separated membraneless compartment that stores maternal mRNAs in oocytes: acts by undergoing liquid-liquid phase separation upon binding to maternal mRNAs (By similarity). Binds to the 3'-UTR of maternal mRNAs, inhibiting their translation (By similarity). {ECO:0000250\|UniProtKB:C3VD30, ECO:0000250\|UniProtKB:Q80SU3}. | *Other* | $MD_{MD}$ |
| A7E2V4 | *ZSWIM8 KIAA0913* | FUNCTION: Substrate recognition component of a SCF-like E3 ubiquitin-protein ligase complex that promotes target-directed microRNA degradation (TDMD), a process that mediates degradation of microRNAs (miRNAs) (PubMed:33184234, PubMed:33184237). The SCF-like E3 ubiquitin-protein ligase complex acts by catalyzing ubiquitination and subsequent degradation of AGO proteins (AGO1, AGO2, AGO3 and/or AGO4), thereby exposing miRNAs for degradation (PubMed:33184234, PubMed:33184237). Specifically recognizes and binds AGO proteins when they are engaged with a TDMD target (PubMed:33184234). May also act as a regulator of axon guidance: specifically recognizes misfolded ROBO3 and promotes its ubiquitination and subsequent degradation (PubMed:24012004). {ECO:0000269\|PubMed:24012004, ECO:0000269\|PubMed:33184234, ECO:0000269\|PubMed:33184237}. | *Other* | $MD_{MD}$ |
| O00764 | *PDXK C21orf124 C21orf97 PKH PNK PRED79* | FUNCTION: Catalyzes the phosphorylation of the dietary vitamin B6 vitamers pyridoxal (PL), pyridoxine (PN) and pyridoxamine (PM) to form pyridoxal 5'-phosphate (PLP), pyridoxine 5'-phosphate (PNP) and pyridoxamine 5'-phosphate (PMP), respectively (PubMed:9099727, PubMed:10987144, PubMed:17766369, PubMed:19351586, PubMed:31187503) (Probable). PLP is the active form of vitamin B6, and acts as a cofactor for over 140 different enzymatic reactions. {ECO:0000269\|PubMed:10987144, ECO:0000269\|PubMed:17766369, ECO:0000269\|PubMed:19351586, ECO:0000269\|PubMed:31187503, ECO:0000269\|PubMed:9099727, ECO:0000305}. | *Other* | $MD_{MD}$ |
| O14686 | *KMT2D ALR MLL2 MLL4* | FUNCTION: Histone methyltransferase that catalyzes methyl group transfer from S-adenosyl-L-methionine to the epsilon-amino group of 'Lys-4' of histone H3 (H3K4) (PubMed:25561738). Part of chromatin remodeling machinery predominantly forms H3K4me1 methylation marks at active chromatin sites where transcription and DNA repair take place (PubMed:25561738, PubMed:17500065). Acts as a coactivator for estrogen receptor by being recruited by ESR1, thereby activating transcription (PubMed:16603732). {ECO:0000269\|PubMed:16603732, ECO:0000269\|PubMed:17500065, ECO:0000269\|PubMed:25561738}. | *Other* | $WTD_{ML}$ |
| O14709 | *ZNF197 ZKSCAN9 ZNF166* | FUNCTION: May be involved in transcriptional regulation. | *Other* | $WTD_{ML}$ |
| O14798 | *TNFRSF10C DCR1 LIT TRAILR3 TRID UNQ321/ PRO366* | FUNCTION: Receptor for the cytotoxic ligand TRAIL. Lacks a cytoplasmic death domain and hence is not capable of inducing apoptosis. May protect cells against TRAIL mediated apoptosis by competing with TRAIL-R1 and R2 for binding to the ligand. | *Other* | $MD_{MD}$ |
| O15427 | *SLC16A3 MCT3 MCT4* | FUNCTION: Proton-dependent transporter of monocarboxylates such as L-lactate and pyruvate (PubMed:11101640, PubMed:23935841, PubMed:31719150). Plays a predominant role in L-lactate efflux from highly glycolytic cells (By similarity). {ECO:0000250\|UniProtKB:O35910, ECO:0000269\|PubMed:11101640, ECO:0000269\|PubMed:23935841, ECO:0000269\|PubMed:31719150}. | *Other* | $MD_{MD}$ |
| O43150 | *ASAP2 DDEF2 KIAA0400* | FUNCTION: Activates the small GTPases ARF1, ARF5 and ARF6. Regulates the formation of post-Golgi vesicles and modulates constitutive secretion. Modulates phagocytosis mediated by Fc gamma receptor and ARF6. Modulates PXN recruitment to focal contacts and cell migration. {ECO:0000269\|PubMed:10022920, ECO:0000269\|PubMed:10749932, ECO:0000269\|PubMed:11304556}. | *Other* | $WTD_{ML}$ |
| O43818 | *RRP9 RNU3IP2 U355K* | FUNCTION: Component of a nucleolar small nuclear ribonucleoprotein particle (snoRNP) thought to participate in the processing and modification of pre-ribosomal RNA (pre-rRNA) (PubMed:26867678). Part of the small subunit (SSU) processome, first precursor of the small eukaryotic ribosomal subunit. During the assembly of the SSU processome in the nucleolus, many ribosome biogenesis factors, an RNA chaperone and ribosomal proteins associate with the nascent pre-rRNA and work in concert to generate RNA folding, modifications, rearrangements and cleavage as well as targeted degradation of pre-ribosomal RNA by the RNA exosome (PubMed:34516797). {ECO:0000269\|PubMed:26867678, ECO:0000269\|PubMed:34516797}. | *Other* | $MD_{BTD}$ |
| O60343 | *TBC1D4 AS160 KIAA0603* | FUNCTION: May act as a GTPase-activating protein for RAB2A, RAB8A, RAB10 and RAB14. Isoform 2 promotes insulin-induced glucose transporter SLC2A4/GLUT4 translocation at the plasma membrane, thus increasing glucose uptake. {ECO:0000269\|PubMed:15971998, ECO:0000269\|PubMed:18771725, ECO:0000269\|PubMed:22908308}. | *Other* | $WTD_{ML}$ |
| O75564 | *JRK JH8* | FUNCTION: May bind DNA. {ECO:0000250}. | *Other* | $MD_{MD}$ |
| O75691 | *UTP20 DRIM* | FUNCTION: Part of the small subunit (SSU) processome, first precursor of the small eukaryotic ribosomal subunit. During the assembly of the SSU processome in the nucleolus, many ribosome biogenesis factors, an RNA chaperone and ribosomal proteins associate with the nascent pre-rRNA and work in concert to generate RNA folding, modifications, rearrangements and cleavage as well as targeted degradation of pre-ribosomal RNA by the RNA exosome. Involved in 18S pre-rRNA processing. Associates with U3 snoRNA. {ECO:0000269\|PubMed:17498821, ECO:0000269\|PubMed:34516797}. | *Other* | $WTD_{ML}$ |
| O75818 | *RPP40 RNASEP1* | FUNCTION: Component of ribonuclease P, a ribonucleoprotein complex that generates mature tRNA molecules by cleaving their 5'-ends (PubMed:9630247, PubMed:30454648). Also a component of the MRP ribonuclease complex, which cleaves pre-rRNA sequences (PubMed:28115465). {ECO:0000269\|PubMed:28115465, ECO:0000269\|PubMed:30454648, ECO:0000269\|PubMed:9630247}. | *Other* | $WTD_{ML}$ |
| O95391 | *SLU7* | FUNCTION: Required for pre-mRNA splicing as component of the spliceosome (PubMed:10197984, PubMed:28502770, PubMed:30705154). Participates in the second catalytic step of pre-mRNA splicing, when the free hydroxyl group of exon I attacks the 3'-splice site to generate spliced mRNA and the excised lariat intron. Required for holding exon 1 properly in the spliceosome and for correct AG identification when more than one possible AG exists in 3'-splicing site region. May be involved in the activation of proximal AG. Probably also involved in alternative splicing regulation. {ECO:0000269\|PubMed:10197984, ECO:0000269\|PubMed:10647016, ECO:0000269\|PubMed:12764196, ECO:0000269\|PubMed:15181151, ECO:0000269\|PubMed:15728250, ECO:0000269\|PubMed:28502770, ECO:0000269\|PubMed:30705154}. | *Other* | $MD_{MD}$ |
| P05060 | *CHGB SCG1* | FUNCTION: Secretogranin-1 is a neuroendocrine secretory granule protein, which may be the precursor for other biologically active peptides. | *Other* | $MD_{BTD}$ |
| P05091 | *ALDH2 ALDM* | FUNCTION: Required for clearance of cellular formaldehyde, a cytotoxic and carcinogenic metabolite that induces DNA damage. {ECO:0000269\|PubMed:33355142}. | *Other* | $WTD_{ML}$ |
| P05160 | *F13B* | FUNCTION: The B chain of factor XIII is not catalytically active, but is thought to stabilize the A subunits and regulate the rate of transglutaminase formation by thrombin. {ECO:0000303\|PubMed:21742792, ECO:0000303\|PubMed:3021194}. | *Other* | $MD_{MD}$ |

| | | | | |
|---|---|---|---|---|
| P07602 | *PSAP* *GLBA* *SAP1* | FUNCTION: Saposin-A and saposin-C stimulate the hydrolysis of glucosylceramide by beta-glucosylceramidase (EC 3.2.1.45) and galactosylceramide by beta-galactosylceramidase (EC 3.2.1.46). Saposin-C apparently acts by combining with the enzyme and acidic lipid to form an activated complex, rather than by solubilizing the substrate.; FUNCTION: Saposin-B stimulates the hydrolysis of galacto-cerebroside sulfate by arylsulfatase A (EC 3.1.6.8), GM1 gangliosides by beta-galactosidase (EC 3.2.1.23) and globotriaosylceramide by alpha-galactosidase A (EC 3.2.1.22). Saposin-B forms a solubilizing complex with the substrates of the sphingolipid hydrolases.; FUNCTION: Saposin-D is a specific sphingomyelin phosphodiesterase activator (EC 3.1.4.12).; FUNCTION: [Prosaposin]: Behaves as a myelinotrophic and neurotrophic factor, these effects are mediated by its G-protein-coupled receptors, GPR37 and GPR37L1, undergoing ligand-mediated internalization followed by ERK phosphorylation signaling. {ECO:0000250\|UniProtKB:Q61207, ECO:0000269\|PubMed:10383054}.; FUNCTION: Saposins are specific low-molecular mass non-enzymic proteins, they participate in the lysosomal degradation of sphingolipids, which takes place by the sequential action of specific hydrolases. | *Other* | $MD_{BTD}$ |
| P08237 | *PFKM* *PFKX* | FUNCTION: Catalyzes the phosphorylation of D-fructose 6-phosphate to fructose 1,6-bisphosphate by ATP, the first committing step of glycolysis. | *Other* | $WTD_{ML}$ |
| P12270 | *TPR* | FUNCTION: Component of the nuclear pore complex (NPC), a complex required for the trafficking across the nuclear envelope. Functions as a scaffolding element in the nuclear phase of the NPC essential for normal nucleocytoplasmic transport of proteins and mRNAs, plays a role in the establishment of nuclear-peripheral chromatin compartmentalization in interphase, and in the mitotic spindle checkpoint signaling during mitosis. Involved in the quality control and retention of unspliced mRNAs in the nucleus; in association with NUP153, regulates the nuclear export of unspliced mRNA species bearing constitutive transport element (CTE) in a NXF1- and KHDRBS1-independent manner. Negatively regulates both the association of CTE-containing mRNA with large polyribosomes and translation initiation. Does not play any role in Rev response element (RRE)-mediated export of unspliced mRNAs. Implicated in nuclear export of mRNAs transcribed from heat shock gene promoters; associates both with chromatin in the HSP70 promoter and with mRNAs transcribed from this promoter under stress-induced conditions. Modulates the nucleocytoplasmic transport of activated MAPK1/ERK2 and huntingtin/HTT and may serve as a docking site for the XPO1/CRM1-mediated nuclear export complex. According to some authors, plays a limited role in the regulation of nuclear protein export (PubMed:22253824, PubMed:11952838). Also plays a role as a structural and functional element of the perinuclear chromatin distribution; involved in the formation and/or maintenance of NPC-associated perinuclear heterochromatin exclusion zones (HEZs). Finally, acts as a spatial regulator of the spindle-assembly checkpoint (SAC) response ensuring a timely and effective recruitment of spindle checkpoint proteins like MAD1L1 and MAD2L1 to unattached kinetochore during the metaphase-anaphase transition before chromosome congression. Its N-terminus is involved in activation of oncogenic kinases. {ECO:0000269\|PubMed:11952838, ECO:0000269\|PubMed:15654337, ECO:0000269\|PubMed:17897941, ECO:0000269\|PubMed:18794356, ECO:0000269\|PubMed:18981471, ECO:0000269\|PubMed:19273613, ECO:0000269\|PubMed:20133940, ECO:0000269\|PubMed:20407419, ECO:0000269\|PubMed:21613532, ECO:0000269\|PubMed:22253824, ECO:0000269\|PubMed:9864356}. | *Other* | $MD_{MD}$ |
| P17516 | *AKR1C4* *CHDR* | FUNCTION: Cytosolic aldo-keto reductase that catalyzes the NADH and NADPH-dependent reduction of ketosteroids to hydroxysteroids. Liver specific enzyme that acts as NAD(P)(H)-dependent 3-, 17- and 20-ketosteroid reductase on the steroid nucleus and side chain (PubMed:14672942, PubMed:10998348, PubMed:7650035, PubMed:1530633, PubMed:11158055, PubMed:10634139, PubMed:19218247). Displays the ability to catalyze both oxidation and reduction in vitro, but most probably acts as a reductase in vivo since the oxidase activity measured in vitro is inhibited by physiological concentration of NADPH (PubMed:14672942). Acts preferentially as a 3-alpha-hydroxysteroid dehydrogenase (HSD) with a subsidiary 3-beta-HSD activity (PubMed:14672942). Catalyzes efficiently the transformation of the potent androgen 5-alpha-dihydrotestosterone (5alpha-DHT or 17beta-hydroxy-5alpha-androstan-3-one) into the less active form, 5-alpha-androstan-3-alpha,17-beta-diol (3-alpha-diol) (PubMed:11158055, PubMed:10998348, PubMed:14672942). Catalyzes the reduction of estrone into 17beta-estradiol but with low efficiency (PubMed:14672942). Metabolizes a broad spectrum of natural and synthetic therapeutic steroid and plays an important role in metabolism of androgens, estrogens, progestereone and conjugated steroids (PubMed:10998348, PubMed:14672942, PubMed:19218247). Catalyzes the biotransformation of the pesticide chlordecone (kepone) to its corresponding alcohol leading to increased biliary excretion of the pesticide and concomitant reduction of its neurotoxicity since bile is the major excretory route (PubMed:2427522). {ECO:0000269\|PubMed:10634139, ECO:0000269\|PubMed:10998348, ECO:0000269\|PubMed:11158055, ECO:0000269\|PubMed:14672942, ECO:0000269\|PubMed:1530633, ECO:0000269\|PubMed:19218247, ECO:0000269\|PubMed:2427522, ECO:0000269\|PubMed:7650035}. | *Other* | $WTD_{ML}$ |
| P18887 | *XRCC1* | FUNCTION: Scaffold protein involved in DNA single-strand break repair by mediating the assembly of DNA break repair protein complexes (PubMed:11163244, PubMed:28002403). Negatively regulates ADP-ribosyltransferase activity of PARP1 during base-excision repair in order to prevent excessive PARP1 activity (PubMed:34102106, PubMed:34811483, PubMed:28002403). Recognizes and binds poly-ADP-ribose chains: specifically binds auto-poly-ADP-ribosylated PARP1, limiting its activity (PubMed:14500814, PubMed:34102106, PubMed:34811483). {ECO:0000269\|PubMed:11163244, ECO:0000269\|PubMed:14500814, ECO:0000269\|PubMed:28002403, ECO:0000269\|PubMed:34102106, ECO:0000269\|PubMed:34811483}. | *Other* | $MD_{MD}$ |
| P20226 | *TBP* *GTF2D1* *TF2D* *TFIID* | FUNCTION: The TFIID basal transcription factor complex plays a major role in the initiation of RNA polymerase II (Pol II)-dependent transcription (PubMed:33795473). TFIID recognizes and binds promoters with or without a TATA box via its subunit TBP, a TATA-box-binding protein, and promotes assembly of the pre-initiation complex (PIC) (PubMed:33795473, PubMed:27193682, PubMed:2194289, PubMed:2363050, PubMed:2374612). The TFIID complex consists of TBP and TBP-associated factors (TAFs), including TAF1, TAF2, TAF3, TAF4, TAF5, TAF6, TAF7, TAF8, TAF9, TAF10, TAF11, TAF12 and TAF13 (PubMed:33795473, PubMed:27007846). The TFIID complex structure can be divided into 3 modules TFIID-A, TFIID-B, and TFIID-C (PubMed:33795473). TBP forms the TFIID-A module together with TAF3 and TAF5 (PubMed:33795473). TBP is a general transcription factor that functions at the core of the TFIID complex (PubMed:33795473, PubMed:27193682, PubMed:2194289, PubMed:2363050, PubMed:2374612, PubMed:9836642). During assembly of the core PIC on the promoter, as part of TFIID, TBP binds to and also bends promoter DNA, irrespective of whether the promoter contains a TATA box (PubMed:33795473). Component of a BRF2-containing transcription factor complex that regulates transcription mediated by RNA polymerase III (PubMed:26638071). Component of the transcription factor SL1/TIF-IB complex, which is involved in the assembly of the PIC during RNA polymerase I-dependent transcription (PubMed:15970593). The rate of PIC formation probably is primarily dependent on the rate of association of SL1 with the rDNA promoter (PubMed:15970593). SL1 is involved in stabilization of nucleolar transcription factor 1/UBTF on rDNA (PubMed:15970593). {ECO:0000269\|PubMed:15970593, ECO:0000269\|PubMed:2194289, ECO:0000269\|PubMed:2363050, ECO:0000269\|PubMed:2374612, ECO:0000269\|PubMed:26638071, ECO:0000269\|PubMed:27007846, ECO:0000269\|PubMed:27193682, ECO:0000269\|PubMed:33795473, ECO:0000269\|PubMed:9836642}. | *Other* | $WTD_{ML}$ |

| | | | | |
|---|---|---|---|---|
| P24385 | *CCND1 BCL1 PRAD1* | FUNCTION: Regulatory component of the cyclin D1-CDK4 (DC) complex that phosphorylates and inhibits members of the retinoblastoma (RB) protein family including RB1 and regulates the cell-cycle during G(1)/S transition (PubMed:1833066, PubMed:1827756, PubMed:8114739, PubMed:8302605, PubMed:19412162, PubMed:33854235). Phosphorylation of RB1 allows dissociation of the transcription factor E2F from the RB/E2F complex and the subsequent transcription of E2F target genes which are responsible for the progression through the G(1) phase (PubMed:1833066, PubMed:1827756, PubMed:8114739, PubMed:8302605, PubMed:19412162). Hypophosphorylates RB1 in early G(1) phase (PubMed:1833066, PubMed:1827756, PubMed:8114739, PubMed:8302605, PubMed:19412162). Cyclin D-CDK4 complexes are major integrators of various mitogenenic and antimitogenic signals (PubMed:1833066, PubMed:1827756, PubMed:8302605, PubMed:19412162). Also a substrate for SMAD3, phosphorylating SMAD3 in a cell-cycle-dependent manner and repressing its transcriptional activity (PubMed:15241418). Component of the ternary complex, cyclin D1/CDK4/CDKN1B, required for nuclear translocation and activity of the cyclin D-CDK4 complex (PubMed:9106657). Exhibits transcriptional corepressor activity with INSM1 on the NEUROD1 and INS promoters in a cell cycle-independent manner (PubMed:16569215, PubMed:18417529). {ECO:0000269\|PubMed:15241418, ECO:0000269\|PubMed:16569215, ECO:0000269\|PubMed:1827756, ECO:0000269\|PubMed:1833066, ECO:0000269\|PubMed:18417529, ECO:0000269\|PubMed:19412162, ECO:0000269\|PubMed:33854235, ECO:0000269\|PubMed:8114739, ECO:0000269\|PubMed:8302605, ECO:0000269\|PubMed:9106657}. | *Other* | $WTD_{ML}$ |
| P28074 | *PSMB5 LMPX MB1 X* | FUNCTION: Component of the 20S core proteasome complex involved in the proteolytic degradation of most intracellular proteins. This complex plays numerous essential roles within the cell by associating with different regulatory particles. Associated with two 19S regulatory particles, forms the 26S proteasome and thus participates in the ATP-dependent degradation of ubiquitinated proteins. The 26S proteasome plays a key role in the maintenance of protein homeostasis by removing misfolded or damaged proteins that could impair cellular functions, and by removing proteins whose functions are no longer required. Associated with the PA200 or PA28, the 20S proteasome mediates ubiquitin-independent protein degradation. This type of proteolysis is required in several pathways including spermatogenesis (20S-PA200 complex) or generation of a subset of MHC class I-presented antigenic peptides (20S-PA28 complex). Within the 20S core complex, PSMB5 displays a chymotrypsin-like activity. {ECO:0000269\|PubMed:15244466, ECO:0000269\|PubMed:18502982, ECO:0000269\|PubMed:18565852, ECO:0000269\|PubMed:27176742, ECO:0000269\|PubMed:8610016}. | *Other* | $MD_{MD}$ |
| P30040 | *ERP29 C12orf8 ERP28* | FUNCTION: Does not seem to be a disulfide isomerase. Plays an important role in the processing of secretory proteins within the endoplasmic reticulum (ER), possibly by participating in the folding of proteins in the ER. | *Other* | $WTD_{ML}$ |
| P35052 | *GPC1* | FUNCTION: Cell surface proteoglycan that bears heparan sulfate. Binds, via the heparan sulfate side chains, alpha-4 (V) collagen and participates in Schwann cell myelination (By similarity). May act as a catalyst in increasing the rate of conversion of prion protein PRPN(C) to PRNP(Sc) via associating (via the heparan sulfate side chains) with both forms of PRPN, targeting them to lipid rafts and facilitating their interaction. Required for proper skeletal muscle differentiation by sequestering FGF2 in lipid rafts preventing its binding to receptors (FGFRs) and inhibiting the FGF-mediated signaling. {ECO:0000250, ECO:0000269\|PubMed:19936054, ECO:0000269\|PubMed:21642435}. | *Other* | $MD_{MD}$ |
| P35241 | *RDX* | FUNCTION: Probably plays a crucial role in the binding of the barbed end of actin filaments to the plasma membrane. | *Other* | $WTD_{ML}$ |
| P35813 | *PPM1A PPPM1A* | FUNCTION: Enzyme with a broad specificity. Negatively regulates TGF-beta signaling through dephosphorylating SMAD2 and SMAD3, resulting in their dissociation from SMAD4, nuclear export of the SMADs and termination of the TGF-beta-mediated signaling. Dephosphorylates PRKAA1 and PRKAA2. Plays an important role in the termination of TNF-alpha-mediated NF-kappa-B activation through dephosphorylating and inactivating IKBKB/IKKB. {ECO:0000269\|PubMed:16751101, ECO:0000269\|PubMed:18930133}. | *Other* | $WTD_{KEY}$ |
| P40616 | *ARL1* | FUNCTION: GTP-binding protein that recruits several effectors, such as golgins, arfaptins and Arf-GEFs to the trans-Golgi network, and modulates their functions at the Golgi complex (PubMed:9624189, PubMed:21239483, PubMed:27436755, PubMed:22679020, PubMed:27373159). Plays thereby a role in a wide range of fundamental cellular processes, including cell polarity, innate immunity, or protein secretion mediated by arfaptins, which were shown to play a role in maintaining insulin secretion from pancreatic beta cells (PubMed:22981988). {ECO:0000269\|PubMed:21239483, ECO:0000269\|PubMed:22679020, ECO:0000269\|PubMed:22981988, ECO:0000269\|PubMed:27373159, ECO:0000269\|PubMed:27436755, ECO:0000269\|PubMed:9624189}. | *Other* | $WTD_{ML}$ |
| P48595 | *SERPINB 10 PI10* | FUNCTION: Protease inhibitor that may play a role in the regulation of protease activities during hematopoiesis and apoptosis induced by TNF. May regulate protease activities in the cytoplasm and in the nucleus. {ECO:0000269\|PubMed:10871600, ECO:0000269\|PubMed:7592909}. | *Other* | $WTD_{ML}$ |
| P48730 | *CSNK1D HCKID* | FUNCTION: Essential serine/threonine-protein kinase that regulates diverse cellular growth and survival processes including Wnt signaling, DNA repair and circadian rhythms. It can phosphorylate a large number of proteins. Casein kinases are operationally defined by their preferential utilization of acidic proteins such as caseins as substrates. Phosphorylates connexin-43/GJA1, MAP1A, SNAPIN, MAPT/TAU, TOP2A, DCK, HIF1A, EIF6, p53/TP53, DVL2, DVL3, ESR1, AIB1/NCOA3, DNMT1, PKD2, YAP1, PER1 and PER2. Central component of the circadian clock. In balance with PP1, determines the circadian period length through the regulation of the speed and rhythmicity of PER1 and PER2 phosphorylation. Controls PER1 and PER2 nuclear transport and degradation. YAP1 phosphorylation promotes its SCF(beta-TRCP) E3 ubiquitin ligase-mediated ubiquitination and subsequent degradation. DNMT1 phosphorylation reduces its DNA-binding activity. Phosphorylation of ESR1 and AIB1/NCOA3 stimulates their activity and coactivation. Phosphorylation of DVL2 and DVL3 regulates WNT3A signaling pathway that controls neurite outgrowth. Phosphorylates NEDD9/HEF1 (By similarity). EIF6 phosphorylation promotes its nuclear export. Triggers down-regulation of dopamine receptors in the forebrain. Activates DCK in vitro by phosphorylation. TOP2A phosphorylation favors DNA cleavable complex formation. May regulate the formation of the mitotic spindle apparatus in extravillous trophoblast. Modulates connexin-43/GJA1 gap junction assembly by phosphorylation. Probably involved in lymphocyte physiology. Regulates fast synaptic transmission mediated by glutamate. {ECO:0000250\|UniProtKB:Q9DC28, ECO:0000269\|PubMed:10606744, ECO:0000269\|PubMed:12270943, ECO:0000269\|PubMed:14761950, ECO:0000269\|PubMed:16027726, ECO:0000269\|PubMed:17562708, ECO:0000269\|PubMed:17962809, ECO:0000269\|PubMed:19043076, ECO:0000269\|PubMed:20041275, ECO:0000269\|PubMed:20048001, ECO:0000269\|PubMed:20407760, ECO:0000269\|PubMed:20637175, ECO:0000269\|PubMed:20696890, ECO:0000269\|PubMed:20699359, ECO:0000269\|PubMed:21084295, ECO:0000269\|PubMed:21422228, ECO:0000269\|PubMed:23636092}. | *Other* | $MD_{MD}$ |
| P49368 | *CCT3 CCTG TRIC5* | FUNCTION: Component of the chaperonin-containing T-complex (TRiC), a molecular chaperone complex that assists the folding of proteins upon ATP hydrolysis (PubMed:25467444). The TRiC complex mediates the folding of WRAP53/TCAB1, thereby regulating telomere maintenance (PubMed:25467444). As part of the TRiC complex may play a role in the assembly of BBSome, a complex involved in ciliogenesis regulating transports vesicles to the cilia (PubMed:20080638). The TRiC complex plays a role in the folding of actin and tubulin (Probable). {ECO:0000269\|PubMed:20080638, ECO:0000269\|PubMed:25467444, ECO:0000305}. | *Other* | $MD_{MD}$ |

| | | | | |
|---|---|---|---|---|
| P51587 | *BRCA2 FACD FANCD1* | FUNCTION: Involved in double-strand break repair and/or homologous recombination. Binds RAD51 and potentiates recombinational DNA repair by promoting assembly of RAD51 onto single-stranded DNA (ssDNA). Acts by targeting RAD51 to ssDNA over double-stranded DNA, enabling RAD51 to displace replication protein-A (RPA) from ssDNA and stabilizing RAD51-ssDNA filaments by blocking ATP hydrolysis. Part of a PALB2-scaffolded HR complex containing RAD51C and which is thought to play a role in DNA repair by HR. May participate in S phase checkpoint activation. Binds selectively to ssDNA, and to ssDNA in tailed duplexes and replication fork structures. May play a role in the extension step after strand invasion at replication-dependent DNA double-strand breaks; together with PALB2 is involved in both POLH localization at collapsed replication forks and DNA polymerization activity. In concert with NPM1, regulates centrosome duplication. Interacts with the TREX-2 complex (transcription and export complex 2) subunits PCID2 and SEM1, and is required to prevent R-loop-associated DNA damage and thus transcription-associated genomic instability. Silencing of BRCA2 promotes R-loop accumulation at actively transcribed genes in replicating and non-replicating cells, suggesting that BRCA2 mediates the control of R-loop associated genomic instability, independently of its known role in homologous recombination (PubMed:24896180). {ECO:0000269\|PubMed:15115758, ECO:0000269\|PubMed:15199141, ECO:0000269\|PubMed:15671039, ECO:0000269\|PubMed:18317453, ECO:0000269\|PubMed:20729832, ECO:0000269\|PubMed:20729858, ECO:0000269\|PubMed:20729859, ECO:0000269\|PubMed:21084279, ECO:0000269\|PubMed:21719596, ECO:0000269\|PubMed:24485656, ECO:0000269\|PubMed:24896180}. | *Other* | $MD_{MD}$ |
| P52849 | *NDST2 HSST2* | FUNCTION: Essential bifunctional enzyme that catalyzes both the N-deacetylation and the N-sulfation of glucosamine (GlcNAc) of the glycosaminoglycan in heparan sulfate. Modifies the GlcNAc-GlcA disaccharide repeating sugar backbone to make N-sulfated heparosan, a prerequisite substrate for later modifications in heparin biosynthesis. Plays a role in determining the extent and pattern of sulfation of heparan sulfate. Required for the exosomal release of SDCBP, CD63 and syndecan (PubMed:22660413). {ECO:0000269\|PubMed:10758005, ECO:0000269\|PubMed:12634318, ECO:0000269\|PubMed:16343444, ECO:0000269\|PubMed:22660413}. | *Other* | $MD_{MD}$ |
| P53992 | *SEC24C KIAA0079* | FUNCTION: Component of the coat protein complex II (COPII) which promotes the formation of transport vesicles from the endoplasmic reticulum (ER). The coat has two main functions, the physical deformation of the endoplasmic reticulum membrane into vesicles and the selection of cargo molecules for their transport to the Golgi complex (PubMed:10214955, PubMed:17499046, PubMed:18843296, PubMed:20427317). Plays a central role in cargo selection within the COPII complex and together with SEC24D may have a different specificity compared to SEC24A and SEC24B (PubMed:17499046, PubMed:20427317, PubMed:18843296). May more specifically package GPI-anchored proteins through the cargo receptor TMED10 (PubMed:20427317). May also be specific for IxM motif-containing cargos like the SNAREs GOSR2 and STX5 (PubMed:18843296). {ECO:0000269\|PubMed:10214955, ECO:0000269\|PubMed:17499046, ECO:0000269\|PubMed:18843296, ECO:0000269\|PubMed:20427317}. | *Other* | $MD_{MD}$ |
| P53999 | *SUB1 PC4 RPO2TC1* | FUNCTION: General coactivator that functions cooperatively with TAFs and mediates functional interactions between upstream activators and the general transcriptional machinery. May be involved in stabilizing the multiprotein transcription complex. Binds single-stranded DNA. Also binds, in vitro, non-specifically to double-stranded DNA (ds DNA). {ECO:0000269\|PubMed:16605275, ECO:0000269\|PubMed:16689930, ECO:0000269\|PubMed:7628453, ECO:0000269\|PubMed:8062391, ECO:0000269\|PubMed:8062392, ECO:0000269\|PubMed:9360603, ECO:0000269\|PubMed:9482861}. | *Other* | $WTD_{KEY}$ |
| P54619 | *PRKAG1* | FUNCTION: AMP/ATP-binding subunit of AMP-activated protein kinase (AMPK), an energy sensor protein kinase that plays a key role in regulating cellular energy metabolism. In response to reduction of intracellular ATP levels, AMPK activates energy-producing pathways and inhibits energy-consuming processes: inhibits protein, carbohydrate and lipid biosynthesis, as well as cell growth and proliferation. AMPK acts via direct phosphorylation of metabolic enzymes, and by longer-term effects via phosphorylation of transcription regulators. Also acts as a regulator of cellular polarity by remodeling the actin cytoskeleton; probably by indirectly activating myosin. Gamma non-catalytic subunit mediates binding to AMP, ADP and ATP, leading to activate or inhibit AMPK: AMP-binding results in allosteric activation of alpha catalytic subunit (PRKAA1 or PRKAA2) both by inducing phosphorylation and preventing dephosphorylation of catalytic subunits. ADP also stimulates phosphorylation, without stimulating already phosphorylated catalytic subunit. ATP promotes dephosphorylation of catalytic subunit, rendering the AMPK enzyme inactive. {ECO:0000269\|PubMed:21680840}. | *Other* | $WTD_{ML}$ |
| P54840 | *GYS2* | FUNCTION: Transfers the glycosyl residue from UDP-Glc to the non-reducing end of alpha-1,4-glucan. | *Other* | $MD_{MD}$ |
| P58397 | *ADAMTS12 UNQ1918/PRO4389* | FUNCTION: Metalloprotease that may play a role in the degradation of COMP. Cleaves also alpha-2 macroglobulin and aggregan. Has anti-tumorigenic properties. {ECO:0000269\|PubMed:16611630, ECO:0000269\|PubMed:17895370, ECO:0000269\|PubMed:18485748}. | *Other* | $WTD_{KEY}$ |
| P61221 | *ABCE1 RLI RNASEL1 RNASELI RNS4I OK/SW-cl.40* | FUNCTION: Nucleoside-triphosphatase (NTPase) involved in ribosome recycling by mediating ribosome disassembly (PubMed:20122402, PubMed:21448132). Able to hydrolyze ATP, GTP, UTP and CTP (PubMed:20122402). Splits ribosomes into free 60S subunits and tRNA- and mRNA-bound 40S subunits (PubMed:20122402, PubMed:21448132). Acts either after canonical termination facilitated by release factors (ETF1/eRF1) or after recognition of stalled and vacant ribosomes by mRNA surveillance factors (PELO/Pelota) (PubMed:20122402, PubMed:21448132). Involved in the No-Go Decay (NGD) pathway: recruited to stalled ribosomes by the Pelota-HBS1L complex, and drives the disassembly of stalled ribosomes, followed by degradation of damaged mRNAs as part of the NGD pathway (PubMed:21448132). Also plays a role in quality control of translation of mitochondrial outer membrane-localized mRNA (PubMed:29861391). As part of the PINK1-regulated signaling, ubiquitinated by CNOT4 upon mitochondria damage; this modification generates polyubiquitin signals that recruit autophagy receptors to the mitochondrial outer membrane and initiate mitophagy (PubMed:29861391). RNASEL-specific protein inhibitor which antagonizes the binding of 2-5A (5'-phosphorylated 2',5'-linked oligoadenylates) to RNASEL (PubMed:9660177). Negative regulator of the anti-viral effect of the interferon-regulated 2-5A/RNASEL pathway (PubMed:9660177, PubMed:9847332, PubMed:11585831). {ECO:0000269\|PubMed:11585831, ECO:0000269\|PubMed:20122402, ECO:0000269\|PubMed:21448132, ECO:0000269\|PubMed:29861391, ECO:0000269\|PubMed:9660177, ECO:0000269\|PubMed:9847332}.; FUNCTION: (Microbial infection) May act as a chaperone for post-translational events during HIV-1 capsid assembly. {ECO:0000269\|PubMed:9847332}.; FUNCTION: (Microbial infection) Plays a role in the down-regulation of the 2-5A/RNASEL pathway during encephalomyocarditis virus (EMCV) and HIV-1 infections. {ECO:0000269\|PubMed:9660177}. | *Other* | $MD_{MD}$ |
| P68363 | *TUBA1B* | FUNCTION: Tubulin is the major constituent of microtubules, a cylinder consisting of laterally associated linear protofilaments composed of alpha- and beta-tubulin heterodimers (PubMed:34996871). Microtubules grow by the addition of GTP-tubulin dimers to the microtubule end, where a stabilizing cap forms (PubMed:34996871). Below the cap, tubulin dimers are in GDP-bound state, owing to GTPase activity of alpha-tubulin (PubMed:34996871). {ECO:0000269\|PubMed:34996871}. | *Other* | $WTD_{ML}$ |
| P83436 | *COG7 UNQ3082/PRO10013* | FUNCTION: Required for normal Golgi function. {ECO:0000269\|PubMed:11980916}. | *Other* | $WTD_{ML}$ |

| | | | | |
|---|---|---|---|---|
| P84074 | *HPCA BDR2* | FUNCTION: Calcium-binding protein that may play a role in the regulation of voltage-dependent calcium channels (PubMed:28398555). May also play a role in cyclic-nucleotide-mediated signaling through the regulation of adenylate and guanylate cyclases (By similarity). {ECO:0000250\|UniProtKB:P84076, ECO:0000269\|PubMed:28398555}. | *Other* | $\mathrm{WTD_{ML}}$ |
| P86397 | *HTD2* | FUNCTION: Mitochondrial 3-hydroxyacyl-thioester dehydratase, which may be involved in fatty acid biosynthesis. {ECO:0000269\|PubMed:17898086}. | *Other* | $\mathrm{WTD_{ML}}$ |
| Q05923 | *DUSP2 PAC1* | FUNCTION: Dephosphorylates both phosphorylated Thr and Tyr residues in MAPK1, and dephosphorylation of phosphotyrosine is slightly faster than that of phosphothreonine (PubMed:8107850). Can dephosphorylate MAPK1 (By similarity). {ECO:0000250\|UniProtKB:Q05922, ECO:0000269\|PubMed:8107850}. | *Other* | $\mathrm{WTD_{ML}}$ |
| Q13398 | *ZNF211* | FUNCTION: May be involved in transcriptional regulation. | *Other* | $\mathrm{WTD_{ML}}$ |
| Q13614 | *MTMR2 KIAA1073* | FUNCTION: Phosphatase that acts on lipids with a phosphoinositol headgroup. Has phosphatase activity towards phosphatidylinositol 3-phosphate and phosphatidylinositol 3,5-bisphosphate (PubMed:11733541, PubMed:12668758, PubMed:21372139, PubMed:14690594). Binds phosphatidylinositol 4-phosphate, phosphatidylinositol 5-phosphate, phosphatidylinositol 3,5-bisphosphate and phosphatidylinositol 3,4,5-trisphosphate (By similarity). Stabilizes SBF2/MTMR13 at the membranes (By similarity). Specifically in peripheral nerves, stabilizes SBF2/MTMR13 protein (By similarity). {ECO:0000250\|UniProtKB:Q9Z2D1, ECO:0000269\|PubMed:11733541, ECO:0000269\|PubMed:12668758, ECO:0000269\|PubMed:14690594, ECO:0000269\|PubMed:21372139}. | *Other* | $\mathrm{WTD_{ML}}$ $\mathrm{MD_{MD}}$ |
| Q14257 | *RCN2 ERC55* | FUNCTION: Not known. Binds calcium. | *Other* | $\mathrm{WTD_{ML}}$ |
| Q14CX7 | *NAA25 C12orf30 MDM20 NAP1* | FUNCTION: Non-catalytic subunit of the NatB complex which catalyzes acetylation of the N-terminal methionine residues of peptides beginning with Met-Asp, Met-Glu, Met-Asn and Met-Gln. May play a role in normal cell-cycle progression. {ECO:0000269\|PubMed:18570629}. | *Other* | $\mathrm{WTD_{ML}}$ |
| Q15393 | *SF3B3 KIAA0017 SAP130* | FUNCTION: Involved in pre-mRNA splicing as a component of the splicing factor SF3B complex, a constituent of the spliceosome (PubMed:10490618, PubMed:10882114, PubMed:27720643, PubMed:28781166). SF3B complex is required for 'A' complex assembly formed by the stable binding of U2 snRNP to the branchpoint sequence (BPS) in pre-mRNA. Sequence independent binding of SF3A/SF3B complex upstream of the branch site is essential, it may anchor U2 snRNP to the pre-mRNA (PubMed:12234937). May also be involved in the assembly of the 'E' complex (PubMed:10882114). As a component of the minor spliceosome, involved in the splicing of U12-type introns in pre-mRNAs (PubMed:15146077) (Probable). {ECO:0000269\|PubMed:10490618, ECO:0000269\|PubMed:10882114, ECO:0000269\|PubMed:12234937, ECO:0000269\|PubMed:15146077, ECO:0000269\|PubMed:27720643, ECO:0000269\|PubMed:28781166, ECO:0000305\|PubMed:33509932}. | *Other* | $\mathrm{WTD_{ML}}$ |
| Q16342 | *PDCD2 RP8 ZMYND7* | FUNCTION: May be a DNA-binding protein with a regulatory function. May play an important role in cell death and/or in regulation of cell proliferation. | *Other* | $\mathrm{WTD_{ML}}$ |
| Q16690 | *DUSP5 VH3* | FUNCTION: Dual specificity protein phosphatase; active with phosphotyrosine, phosphoserine and phosphothreonine residues. The highest relative activity is toward ERK1. {ECO:0000269\|PubMed:7961985}. | *Other* | $\mathrm{MD_{MD}}$ |
| Q17RD7 | *SYT16 STREP14 SYT14L SYT14R* | FUNCTION: May be involved in the trafficking and exocytosis of secretory vesicles in non-neuronal tissues. Is Ca(2+)-independent. | *Other* | $\mathrm{MD_{BTD}}$ |
| Q495W5 | *FUT11* | FUNCTION: [Isoform 1]: Has minor fucosyltransferase activity toward biantennary N-glycan acceptors. Does not fucosylate GlcNAc residue within type 2 lactosamine unit. {ECO:0000269\|PubMed:19088067}.; FUNCTION: [Isoform 2]: Has fucosyltransferase activity toward biantennary N-glycan acceptors. Does not fucosylate GlcNAc residue within type 2 lactosamine unit. {ECO:0000269\|PubMed:19088067}. | *Other* | $\mathrm{MD_{MD}}$ |
| Q53GS7 | *GLE1 GLE1L* | FUNCTION: Required for the export of mRNAs containing poly(A) tails from the nucleus into the cytoplasm. May be involved in the terminal step of the mRNA transport through the nuclear pore complex (NPC). {ECO:0000269\|PubMed:12668658, ECO:0000269\|PubMed:16000379, ECO:0000269\|PubMed:9618489}. | *Other* | $\mathrm{MD_{MD}}$ |
| Q5SGD2 | *PPM1L PP2CE* | FUNCTION: Acts as a suppressor of the SAPK signaling pathways by associating with and dephosphorylating MAP3K7/TAK1 and MAP3K5, and by attenuating the association between MAP3K7/TAK1 and MAP2K4 or MAP2K6. {ECO:0000269\|PubMed:17456047}. | *Other* | $\mathrm{MD_{BTD}}$ |
| Q5SWA1 | *PPP1R15 B* | FUNCTION: Maintains low levels of EIF2S1 phosphorylation in unstressed cells by promoting its dephosphorylation by PP1. {ECO:0000269\|PubMed:26159176, ECO:0000269\|PubMed:26307080}. | *Other* | $\mathrm{MD_{MD}}$ |
| Q5VVC0 | *SPO16 C1orf146 SCRE* | FUNCTION: Plays a key role in reinforcing the integrity of the central element of the synaptonemal complex (SC) thereby stabilizing SC, ensuring progression of meiotic prophase I in male and female germ cells (By similarity). Promotes homologous recombination and crossing-over in meiotic prophase I via its association with SHOC1 (By similarity). Required for the localization of TEX11 and MSH4 to recombination intermediates (By similarity). {ECO:0000250\|UniProtKB:Q3KQP7}. | *Other* | $\mathrm{WTD_{ML}}$ |
| Q641Q2 | *WASHC2 A FAM21A FAM21B* | FUNCTION: Acts at least in part as component of the WASH core complex whose assembly at the surface of endosomes inhibits WASH nucleation-promoting factor (NPF) activity in recruiting and activating the Arp2/3 complex to induce actin polymerization and is involved in the fission of tubules that serve as transport intermediates during endosome sorting. Mediates the recruitment of the WASH core complex to endosome membranes via binding to phospholipids and VPS35 of the retromer CSC. Mediates the recruitment of the F-actin-capping protein dimer to the WASH core complex probably promoting localized F-actin polymerization needed for vesicle scission. Via its C-terminus binds various phospholipids, most strongly phosphatidylinositol 4-phosphate (PtdIns-(4)P), phosphatidylinositol 5-phosphate (PtdIns-(5)P) and phosphatidylinositol 3,5-bisphosphate (PtdIns-(3,5)P2). Involved in the endosome-to-plasma membrane trafficking and recycling of SNX27-retromer-dependent cargo proteins, such as GLUT1. Required for the association of DNAJC13, ENTR1, ANKRD50 with retromer CSC subunit VPS35. Required for the endosomal recruitment of CCC complex subunits COMMD1 and CCDC93 as well as the retriever complex subunit VPS35L. {ECO:0000269\|PubMed:25355947, ECO:0000269\|PubMed:28892079}. | *Other* | $\mathrm{WTD_{ML}}$ |
| Q68DI1 | *ZNF776* | FUNCTION: May be involved in transcriptional regulation. {ECO:0000250}. | *Other* | $\mathrm{WTD_{KEY}}$ $\mathrm{WTD_{ML}}$ |
| Q6GYQ0 | *RALGAPA 1 GARNL1 KIAA0884 TULIP1* | FUNCTION: Catalytic subunit of the heterodimeric RalGAP1 complex which acts as a GTPase activator for the Ras-like small GTPases RALA and RALB. {ECO:0000250}. | *Other* | $\mathrm{WTD_{KEY}}$ |

| | | | | |
|---|---|---|---|---|
| Q6SJ93 | *FAM111B CANP* | FUNCTION: Serine protease. {ECO:0000250\|UniProtKB:Q96PZ2}. | *Other* | WTD$_{ML}$ |
| Q6UWI2 | *PARM1 UNQ1879/PRO4322* | FUNCTION: May regulate TLP1 expression and telomerase activity, thus enabling certain prostatic cells to resist apoptosis. {ECO:0000250}. | *Other* | WTD$_{ML}$ |
| Q6UWZ7 | *ABRAXAS1 ABRA1 CCDC98 FAM175A UNQ496/PRO1013* | FUNCTION: Involved in DNA damage response and double-strand break (DSB) repair. Component of the BRCA1-A complex, acting as a central scaffold protein that assembles the various components of the complex and mediates the recruitment of BRCA1. The BRCA1-A complex specifically recognizes 'Lys-63'-linked ubiquitinated histones H2A and H2AX at DNA lesion sites, leading to target the BRCA1-BARD1 heterodimer to sites of DNA damage at DSBs. This complex also possesses deubiquitinase activity that specifically removes 'Lys-63'-linked ubiquitin on histones H2A and H2AX. {ECO:0000269\|PubMed:17525340, ECO:0000269\|PubMed:17643121, ECO:0000269\|PubMed:17643122, ECO:0000269\|PubMed:18077395, ECO:0000269\|PubMed:19261748, ECO:0000269\|PubMed:22357538, ECO:0000269\|PubMed:26778126}. | *Other* | WTD$_{KEY}$ |
| Q6ZXV5 | *TMTC3* | FUNCTION: Transfers mannosyl residues to the hydroxyl group of serine or threonine residues. The 4 members of the TMTC family are O-mannosyl-transferases dedicated primarily to the cadherin superfamily, each member seems to have a distinct role in decorating the cadherin domains with O-linked mannose glycans at specific regions. Also acts as O-mannosyl-transferase on other proteins such as PDIA3 (PubMed:28973932). Involved in the positive regulation of proteasomal protein degradation in the endoplasmic reticulum (ER), and the control of ER stress response. {ECO:0000269\|PubMed:21603654, ECO:0000269\|PubMed:28973932}. | *Other* | WTD$_{KEY}$ |
| Q75T13 | *PGAP1 UNQ3024/PRO9822* | FUNCTION: Involved in inositol deacylation of GPI-anchored proteins. GPI inositol deacylation may important for efficient transport of GPI-anchored proteins from the endoplasmic reticulum to the Golgi (By similarity). {ECO:0000250}. | *Other* | MD$_{MD}$ |
| Q7L590 | *MCM10 PRO2249* | FUNCTION: Acts as a replication initiation factor that brings together the MCM2-7 helicase and the DNA polymerase alpha/primase complex in order to initiate DNA replication. Additionally, plays a role in preventing DNA damage during replication. Key effector of the RBBP6 and ZBTB38-mediated regulation of DNA-replication and common fragile sites stability; acts as a direct target of transcriptional repression by ZBTB38 (PubMed:24726359). {ECO:0000269\|PubMed:11095689, ECO:0000269\|PubMed:15136575, ECO:0000269\|PubMed:17699597, ECO:0000269\|PubMed:19608746, ECO:0000269\|PubMed:24726359, ECO:0000269\|PubMed:32865517}. | *Other* | WTD$_{ML}$ |
| Q7Z419 | *RNF144B IBRDC2 P53RFP* | FUNCTION: E3 ubiquitin-protein ligase which accepts ubiquitin from E2 ubiquitin-conjugating enzymes UBE2L3 and UBE2L6 in the form of a thioester and then directly transfers the ubiquitin to targeted substrates such as LCMT2, thereby promoting their degradation. Induces apoptosis via a p53/TP53-dependent but caspase-independent mechanism. However, its overexpression also produces a decrease of the ubiquitin-dependent stability of BAX, a pro-apoptotic protein, ultimately leading to protection of cell death; But, it is not an anti-apoptotic protein per se. {ECO:0000269\|PubMed:12853982, ECO:0000269\|PubMed:20300062}. | *Other* | MD$_{MD}$ |
| Q7Z5R6 | *APBB1IP PREL1 RARP1 RIAM* | FUNCTION: Appears to function in the signal transduction from Ras activation to actin cytoskeletal remodeling. Suppresses insulin-induced promoter activities through AP1 and SRE. Mediates Rap1-induced adhesion. {ECO:0000269\|PubMed:14530287, ECO:0000269\|PubMed:15469846}. | *Other* | WTD$_{KEY}$ |
| Q86XI2 | *NCAPG2 LUZP5* | FUNCTION: Regulatory subunit of the condensin-2 complex, a complex which establishes mitotic chromosome architecture and is involved in physical rigidity of the chromatid axis. {ECO:0000269\|PubMed:14532007, ECO:0000269\|PubMed:30609410}. | *Other* | MD$_{MD}$ |
| Q8IUB5 | *WFDC13 C20orf138 WAP13* | FUNCTION: Putative acid-stable proteinase inhibitor. {ECO:0000250}. | *Other* | MD$_{BTD}$ |
| Q8IXH6 | *TP53INP2 C20orf110 DOR PINH* | FUNCTION: Dual regulator of transcription and autophagy. Positively regulates autophagy and is required for autophagosome formation and processing. May act as a scaffold protein that recruits MAP1LC3A, GABARAP and GABARAPL2 and brings them to the autophagosome membrane by interacting with VMP1 where, in cooperation with the BECN1-PI3-kinase class III complex, they trigger autophagosome development. Acts as a transcriptional activator of THRA. {ECO:0000269\|PubMed:18030323, ECO:0000269\|PubMed:19056683, ECO:0000269\|PubMed:22470510}. | *Other* | WTD$_{ML}$ |
| Q8IXI2 | *RHOT1 ARHT1* | FUNCTION: Mitochondrial GTPase involved in mitochondrial trafficking (PubMed:12482879, PubMed:16630562, PubMed:22396657). Probably involved in control of anterograde transport of mitochondria and their subcellular distribution (PubMed:12482879, PubMed:16630562, PubMed:22396657). Promotes mitochondrial fission during high calcium conditions (PubMed:27716788). {ECO:0000269\|PubMed:12482879, ECO:0000269\|PubMed:16630562, ECO:0000269\|PubMed:22396657, ECO:0000269\|PubMed:27716788}. | *Other* | WTD$_{ML}$ |
| Q8IYS1 | *PM20D2 ACY1L2* | FUNCTION: Catalyzes the peptide bond hydrolysis in dipeptides having basic amino acids lysine, ornithine or arginine at C-terminus. Postulated to function in a metabolite repair mechanism by eliminating alternate dipeptide by-products formed during carnosine synthesis. {ECO:0000269\|PubMed:24891507}. | *Other* | MD$_{MD}$ |
| Q8N0Z2 | *ABRA* | FUNCTION: Acts as an activator of serum response factor (SRF)-dependent transcription possibly by inducing nuclear translocation of MKL1 or MKL2 and through a mechanism requiring Rho-actin signaling. {ECO:0000250\|UniProtKB:Q8BUZ1}. | *Other* | WTD$_{ML}$ |
| Q8N3J9 | *ZNF664 ZFOC1 ZNF176* | FUNCTION: May be involved in transcriptional regulation. | *Other* | MD$_{MD}$ |
| Q8N3P4 | *VPS8 KIAA0804* | FUNCTION: Plays a role in vesicle-mediated protein trafficking of the endocytic membrane transport pathway. Believed to act as a component of the putative CORVET endosomal tethering complexes which is proposed to be involved in the Rab5-to-Rab7 endosome conversion probably implicating MON1A/B, and via binding SNAREs and SNARE complexes to mediate tethering and docking events during SNARE-mediated membrane fusion. The CORVET complex is proposed to function as a Rab5 effector to mediate early endosome fusion probably in specific endosome subpopulations (PubMed:25266290). Functions predominantly in APPL1-containing endosomes (PubMed:25266290). {ECO:0000269\|PubMed:25266290, ECO:0000305\|PubMed:25266290}. | *Other* | MD$_{MD}$ |
| Q8N465 | *D2HGDH D2HGD* | FUNCTION: Catalyzes the oxidation of D-2-hydroxyglutarate (D-2-HG) to alpha-ketoglutarate (PubMed:15070399, PubMed:15609246, PubMed:16037974, PubMed:20020533, PubMed:33431826). Also catalyzes the oxidation of other D-2-hydroxyacids, such as D-malate (D-MAL) and D-lactate (D-LAC) (PubMed:33431826). Exhibits high activities towards D-2-HG and D-MAL but a very weak activity towards D-LAC (PubMed:33431826). {ECO:0000269\|PubMed:15070399, ECO:0000269\|PubMed:15609246, ECO:0000269\|PubMed:16037974, ECO:0000269\|PubMed:20020533, ECO:0000269\|PubMed:33431826}. | *Other* | WTD$_{ML}$ |

| | | | | |
|---|---|---|---|---|
| Q8N556 | *AFAP1*<br>*AFAP* | FUNCTION: Can cross-link actin filaments into both network and bundle structures (By similarity). May modulate changes in actin filament integrity and induce lamellipodia formation. May function as an adapter molecule that links other proteins, such as SRC and PKC to the actin cytoskeleton. Seems to play a role in the development and progression of prostate adenocarcinoma by regulating cell-matrix adhesions and migration in the cancer cells. {ECO:0000250, ECO:0000269\|PubMed:15485829}. | *Other* | $MD_{MD}$ |
| Q8N573 | *OXR1*<br>*Nbla003*<br>*07* | FUNCTION: May be involved in protection from oxidative damage. {ECO:0000269\|PubMed:11114193, ECO:0000269\|PubMed:15060142}. | *Other* | $WTD_{ML}$ |
| Q8NA72 | *POC5*<br>*C5orf37* | FUNCTION: Essential for the assembly of the distal half of centrioles, required for centriole elongation. {ECO:0000269\|PubMed:19349582}. | *Other* | $WTD_{ML}$<br>$MD_{MD}$ |
| Q8TDG4 | *HELQ*<br>*HEL308* | FUNCTION: Single-stranded 3'-5' DNA helicase that plays a key role in homology-driven double-strand break (DSB) repair (PubMed:11751861, PubMed:19995904, PubMed:21398521, PubMed:24005041, PubMed:24005565, PubMed:34316696, PubMed:34937945). Involved in different DSB repair mechanisms that are guided by annealing of extensive stretches of complementary bases at break ends, such as microhomology-mediated end-joining (MMEJ), single-strand annealing (SSA) or synthesis-dependent strand annealing (SDSA) (PubMed:34937945). Possesses both DNA unwinding and annealing activities (PubMed:34937945). Forms a complex with RAD51, stimulating HELQ DNA helicase activity and ability to unwing DNA (PubMed:34937945). Efficiently unwinds substrates containing 3' overhangs or a D-loop (PubMed:21398521, PubMed:34937945). In contrast, interaction with the replication protein A (RPA/RP-A) complex inhibits DNA unwinding by HELQ but strongly stimulates DNA strand annealing (PubMed:34937945). Triggers displacement of RPA from single-stranded DNA to facilitate annealing of complementary sequences (PubMed:34316696, PubMed:34937945). {ECO:0000269\|PubMed:11751861, ECO:0000269\|PubMed:19995904, ECO:0000269\|PubMed:21398521, ECO:0000269\|PubMed:24005041, ECO:0000269\|PubMed:24005565, ECO:0000269\|PubMed:34316696, ECO:0000269\|PubMed:34937945}. | *Other* | $WTD_{KEY}$ |
| Q8WWB<br>7 | *GLMP*<br>*C1orf85*<br>*PSEC003*<br>*0*<br>*UNQ2553*<br>*/PRO618*<br>*2* | FUNCTION: Required to protect lysosomal transporter MFSD1 from lysosomal proteolysis and for MFSD1 lysosomal localization. {ECO:0000250\|UniProtKB:Q9JHJ3}. | *Other* | $MD_{MD}$ |
| Q8WWQ<br>2 | *HPSE2*<br>*HPA2* | FUNCTION: Binds heparin and heparan sulfate with high affinity, but lacks heparanase activity. Inhibits HPSE, possibly by competing for its substrates (in vitro). {ECO:0000269\|PubMed:20576607}. | *Other* | $WTD_{ML}$ |
| Q8WXF0 | *SRSF12*<br>*SFRS13B*<br>*SFRS19*<br>*SRRP35* | FUNCTION: Splicing factor that seems to antagonize SR proteins in pre-mRNA splicing regulation. {ECO:0000269\|PubMed:11684676}. | *Other* | $MD_{MD}$ |
| Q8WYH8 | *ING5* | FUNCTION: Component of the HBO1 complex, which specifically mediates acetylation of histone H3 at 'Lys-14' (H3K14ac) and, to a lower extent, acetylation of histone H4 (PubMed:24065767). Component of the MOZ/MORF complex which has a histone H3 acetyltransferase activity (PubMed:16387653). Through chromatin acetylation it may regulate DNA replication and may function as a transcriptional coactivator (PubMed:12750254, PubMed:16387653). Inhibits cell growth, induces a delay in S-phase progression and enhances Fas-induced apoptosis in an INCA1-dependent manner (PubMed:21750715). {ECO:0000269\|PubMed:12750254, ECO:0000269\|PubMed:16387653, ECO:0000269\|PubMed:21750715, ECO:0000269\|PubMed:24065767}. | *Other* | $WTD_{ML}$ |
| Q969S9 | *GFM2*<br>*EFG2*<br>*MSTP027* | FUNCTION: Mitochondrial GTPase that mediates the disassembly of ribosomes from messenger RNA at the termination of mitochondrial protein biosynthesis. Acts in collaboration with MRRF. GTP hydrolysis follows the ribosome disassembly and probably occurs on the ribosome large subunit. Not involved in the GTP-dependent ribosomal translocation step during translation elongation. {ECO:0000255\|HAMAP-Rule:MF_03059, ECO:0000269\|PubMed:19716793}. | *Other* | $MD_{MD}$ |
| Q96A04 | *TSACC*<br>*C1orf182* | FUNCTION: Co-chaperone that facilitates HSP-mediated activation of TSSK6. {ECO:0000269\|PubMed:20829357}. | *Other* | $MD_{MD}$ |
| Q96A23 | *CPNE4* | FUNCTION: Probable calcium-dependent phospholipid-binding protein that may play a role in calcium-mediated intracellular processes. {ECO:0000250\|UniProtKB:Q99829}. | *Other* | $WTD_{ML}$ |
| Q96B42 | *TMEM18* | FUNCTION: Transcription repressor. Sequence-specific ssDNA and dsDNA binding protein, with preference for GCT end CTG repeats. Cell migration modulator which enhances the glioma-specific migration ability of neural stem cells (NSC) and neural precursor cells (NPC). {ECO:0000269\|PubMed:18559506, ECO:0000269\|PubMed:21980424}. | *Other* | $WTD_{ML}$<br>$MD_{MD}$ |
| Q96BM0 | *IFI27L1*<br>*FAM14B* | FUNCTION: Plays a role in the apoptotic process and has a pro-apoptotic activity. {ECO:0000269\|PubMed:27673746}. | *Other* | $WTD_{ML}$ |
| Q96D46 | *NMD3*<br>*CGI-07* | FUNCTION: Acts as an adapter for the XPO1/CRM1-mediated export of the 60S ribosomal subunit. {ECO:0000269\|PubMed:12724356, ECO:0000269\|PubMed:12773398}. | *Other* | $MD_{BTD}$ |
| Q96IF1 | *AJUBA*<br>*JUB* | FUNCTION: Adapter or scaffold protein which participates in the assembly of numerous protein complexes and is involved in several cellular processes such as cell fate determination, cytoskeletal organization, repression of gene transcription, mitosis, cell-cell adhesion, cell differentiation, proliferation and migration. Contributes to the linking and/or strengthening of epithelia cell-cell junctions in part by linking adhesive receptors to the actin cytoskeleton. May be involved in signal transduction from cell adhesion sites to the nucleus. Plays an important role in regulation of the kinase activity of AURKA for mitotic commitment. Also a component of the IL-1 signaling pathway modulating IL-1-induced NFKB1 activation by influencing the assembly and activity of the PRKCZ-SQSTM1-TRAF6 multiprotein signaling complex. Functions as an HDAC-dependent corepressor for a subset of GFI1 target genes. Acts as a transcriptional corepressor for SNAI1 and SNAI2/SLUG-dependent repression of E-cadherin transcription. Acts as a hypoxic regulator by bridging an association between the prolyl hydroxylases and VHL enabling efficient degradation of HIF1A. Positively regulates microRNA (miRNA)-mediated gene silencing. Negatively regulates the Hippo signaling pathway and antagonizes phosphorylation of YAP1. {ECO:0000269\|PubMed:12417594, ECO:0000269\|PubMed:13678582, ECO:0000269\|PubMed:15870274, ECO:0000269\|PubMed:16413547, ECO:0000269\|PubMed:17909014, ECO:0000269\|PubMed:18805794, ECO:0000269\|PubMed:20303269, ECO:0000269\|PubMed:20616046, ECO:0000269\|PubMed:22286099}. | *Other* | $MD_{MD}$ |
| Q96IY4 | *CPB2* | FUNCTION: Cleaves C-terminal arginine or lysine residues from biologically active peptides such as kinins or anaphylatoxins in the circulation thereby regulating their activities. Down-regulates fibrinolysis by removing C-terminal lysine residues from fibrin that has already been partially degraded by plasmin. {ECO:0000269\|PubMed:10574983}. | *Other* | $WTD_{ML}$ |

| | | | | |
|---|---|---|---|---|
| Q96JM3 | *CHAMP1 C13orf8 CAMP CHAMP KIAA1802 ZNF828* | FUNCTION: Required for proper alignment of chromosomes at metaphase and their accurate segregation during mitosis. Involved in the maintenance of spindle microtubules attachment to the kinetochore during sister chromatid biorientation. May recruit CENPE and CENPF to the kinetochore. {ECO:0000269\|PubMed:21063390}. | *Other* | $WTD_{ML}$ |
| Q96NW7 | *LRRC7 KIAA1365 LAP1* | FUNCTION: Required for normal synaptic spine architecture and function. Necessary for DISC1 and GRM5 localization to postsynaptic density complexes and for both N-methyl D-aspartate receptor-dependent and metabotropic glutamate receptor-dependent long term depression. {ECO:0000269\|PubMed:11729199}. | *Other* | $MD_{MD}$ |
| Q96QG7 | *MTMR9 C8orf9 MTMR8* | FUNCTION: Acts as an adapter for myotubularin-related phosphatases (PubMed:19038970, PubMed:22647598). Increases lipid phosphatase MTMR6 catalytic activity, specifically towards phosphatidylinositol 3,5-bisphosphate and MTMR6 binding affinity for phosphorylated phosphatidylinositols (PubMed:19038970, PubMed:22647598). Positively regulates lipid phosphatase MTMR7 catalytic activity (By similarity). Increases MTMR8 catalytic activity towards phosphatidylinositol 3-phosphate (PubMed:22647598). The formation of the MTMR6-MTMR9 complex, stabilizes both MTMR6 and MTMR9 protein levels (PubMed:19038970). Stabilizes MTMR8 protein levels (PubMed:22647598). Plays a role in the late stages of macropinocytosis possibly by regulating MTMR6-mediated dephosphorylation of phosphatidylinositol 3-phosphate in membrane ruffles (PubMed:24591580). Negatively regulates autophagy, in part via its association with MTMR8 (PubMed:22647598). Negatively regulates DNA damage-induced apoptosis, in part via its association with MTMR6 (PubMed:19038970, PubMed:22647598). Does not bind mono-, di- and tri-phosphorylated phosphatidylinositols, phosphatidic acid and phosphatidylserine (PubMed:19038970). {ECO:0000250\|UniProtKB:Q9Z2D0, ECO:0000269\|PubMed:19038970, ECO:0000269\|PubMed:22647598, ECO:0000269\|PubMed:24591580}. | *Other* | $WTD_{ML}$ |
| Q99547 | *MPHOSP H6 MPP6* | FUNCTION: RNA-binding protein that associates with the RNA exosome complex. Involved in the 3'-processing of the 7S pre-RNA to the mature 5.8S rRNA and play a role in recruiting the RNA exosome complex to pre-rRNA; this function may include C1D. {ECO:0000269\|PubMed:17412707, ECO:0000269\|PubMed:26166824}. | *Other* | $MD_{MD}$ |
| Q9BQE5 | *APOL2* | FUNCTION: May affect the movement of lipids in the cytoplasm or allow the binding of lipids to organelles. | *Other* | $WTD_{KEY}$ |
| Q9BTT4 | *MED10 L6 TRG17 TRG20* | FUNCTION: Component of the Mediator complex, a coactivator involved in the regulated transcription of nearly all RNA polymerase II-dependent genes. Mediator functions as a bridge to convey information from gene-specific regulatory proteins to the basal RNA polymerase II transcription machinery. Mediator is recruited to promoters by direct interactions with regulatory proteins and serves as a scaffold for the assembly of a functional preinitiation complex with RNA polymerase II and the general transcription factors. | *Other* | $WTD_{ML}$ |
| Q9BUT1 | *BDH2 DHRS6 SDR15C1 UNQ6308 /PRO209 33* | FUNCTION: NAD(H)-dependent dehydrogenase/reductase with a preference for cyclic substrates (PubMed:35150746) (By similarity). Catalyzes stereoselective conversion of 4-oxo-L-proline to cis-4-hydroxy-L-proline, likely a detoxification mechanism for ketoprolines (PubMed:35150746). Mediates the formation of 2,5-dihydroxybenzoate (2,5-DHBA), a siderophore that chelates free cytoplasmic iron and associates with LCN2, thereby regulating iron transport and homeostasis while protecting cells against free radical-induced oxidative stress. The iron-siderophore complex is imported into mitochondria, providing an iron source for mitochondrial metabolic processes in particular heme synthesis (By similarity). May act as a 3-hydroxybutyrate dehydrogenase (PubMed:16380372). {ECO:0000250\|UniProtKB:Q8JZV9, ECO:0000269\|PubMed:16380372, ECO:0000269\|PubMed:35150746}. | *Other* | $WTD_{ML}$ |
| Q9BXR6 | *CFHR5 CFHL5 FHR5* | FUNCTION: Involved in complement regulation. The dimerized forms have avidity for tissue-bound complement fragments and efficiently compete with the physiological complement inhibitor CFH. {ECO:0000269\|PubMed:23487775}. | *Other* | $MD_{MD}$ |
| Q9BY50 | *SEC11C SEC11L3 SPC21 SPCS4C* | FUNCTION: Catalytic component of the signal peptidase complex (SPC) which catalyzes the cleavage of N-terminal signal sequences from nascent proteins as they are translocated into the lumen of the endoplasmic reticulum (PubMed:34388369). Specifically cleaves N-terminal signal peptides that contain a hydrophobic alpha-helix (h-region) shorter than 18-20 amino acids (PubMed:34388369). {ECO:0000269\|PubMed:34388369}. | *Other* | $MD_{MD}$ |
| Q9BZH6 | *WDR11 BRWD2 KIAA1351 WDR15* | FUNCTION: Involved in the Hedgehog (Hh) signaling pathway, is essential for normal ciliogenesis (PubMed:29263200). Regulates the proteolytic processing of GLI3 and cooperates with the transcription factor EMX1 in the induction of downstream Hh pathway gene expression and gonadotropin-releasing hormone production (PubMed:29263200). WDR11 complex facilitates the tethering of Adaptor protein-1 complex (AP-1)-derived vesicles. WDR11 complex acts together with TBC1D23 to facilitate the golgin-mediated capture of vesicles generated using AP-1 (PubMed:29426865). {ECO:0000269\|PubMed:29263200, ECO:0000269\|PubMed:29426865}. | *Other* | $WTD_{KEY}$ |
| Q9H6Z4 | *RANBP3* | FUNCTION: Acts as a cofactor for XPO1/CRM1-mediated nuclear export, perhaps as export complex scaffolding protein. Bound to XPO1/CRM1, stabilizes the XPO1/CRM1-cargo interaction. In the absence of Ran-bound GTP prevents binding of XPO1/CRM1 to the nuclear pore complex. Binds to CHC1/RCC1 and increases the guanine nucleotide exchange activity of CHC1/RCC1. Recruits XPO1/CRM1 to CHC1/RCC1 in a Ran-dependent manner. Negative regulator of TGF-beta signaling through interaction with the R-SMAD proteins, SMAD2 and SMAD3, and mediating their nuclear export. {ECO:0000269\|PubMed:11425870, ECO:0000269\|PubMed:11571268, ECO:0000269\|PubMed:11932251, ECO:0000269\|PubMed:19289081, ECO:0000269\|PubMed:9637251}. | *Other* | $WTD_{ML}$ $MD_{BTD}$ |
| Q9H765 | *ASB8 PP14212* | FUNCTION: May be a substrate-recognition component of a SCF-like ECS (Elongin-Cullin-SOCS-box protein) E3 ubiquitin-protein ligase complex which mediates the ubiquitination and subsequent proteasomal degradation of target proteins. {ECO:0000250}. | *Other* | $WTD_{ML}$ |
| Q9H777 | *ELAC1 D29* | FUNCTION: Zinc phosphodiesterase, which displays some tRNA 3'-processing endonuclease activity (PubMed:12711671, PubMed:32075755). Specifically involved in tRNA repair: acts downstream of the ribosome-associated quality control (RQC) pathway by removing a 2',3'-cyclic phosphate from tRNAs following cleavage by ANKZF1 (PubMed:32075755). tRNAs are then processed by TRNT1 (PubMed:32075755). {ECO:0000269\|PubMed:12711671, ECO:0000269\|PubMed:32075755}. | *Other* | $WTD_{ML}$ |
| Q9H790 | *EXO5 C1orf176 DEM1* | FUNCTION: Single-stranded DNA (ssDNA) bidirectional exonuclease involved in DNA repair. Probably involved in DNA repair following ultraviolet (UV) irradiation and interstrand cross-links (ICLs) damage. Has both 5'-3' and 3'-5' exonuclease activities with a strong preference for 5'-ends. Acts as a sliding exonuclease that loads at ssDNA ends and then slides along the ssDNA prior to cutting; however the sliding and the 3'-5' exonuclease activities are abolished upon binding to the replication protein A (RPA) complex that enforces 5'-directionality activity. {ECO:0000269\|PubMed:23095756}. | *Other* | $WTD_{ML}$ $MD_{MD}$ |
| Q9H813 | *PACC1 C1orf75 TMEM20 6* | FUNCTION: Proton-activated chloride channel that mediates import of chloride ion in response to extracellular acidic pH (PubMed:31023925, PubMed:31318332). Involved in acidosis-induced cell death by mediating chloride influx and subsequent cell swelling (PubMed:31023925, PubMed:31318332). {ECO:0000269\|PubMed:31023925, ECO:0000269\|PubMed:31318332}. | *Other* | $MD_{MD}$ |
| Q9H9E3 | *COG4* | FUNCTION: Required for normal Golgi function (PubMed:19536132, PubMed:30290151). Plays a role in SNARE-pin assembly and Golgi-to-ER retrograde transport via its interaction with SCFD1 (PubMed:19536132). {ECO:0000269\|PubMed:19536132, ECO:0000269\|PubMed:30290151}. | *Other* | $WTD_{ML}$ |

| | | | | |
|---|---|---|---|---|
| Q9HC07 | *TMEM165 TPARL* | FUNCTION: May function as a calcium/proton transporter involved in calcium and in lysosomal pH homeostasis. Therefore, it may play an indirect role in protein glycosylation. {ECO:0000269\|PubMed:22683087, ECO:0000269\|PubMed:23569283}. | *Other* | $WTD_{ML}$ |
| Q9NP91 | *SLC6A20 SIT1 XT3 XTRP3* | FUNCTION: Mediates the Na(+)- and Cl(-)-dependent uptake of imino acids such as L-proline, N-methyl-L-proline and pipecolate as well as N-methylated amino acids (PubMed:15632147, PubMed:19033659, PubMed:33428810). Also transports glycine, regulates proline and glycine homeostasis in the brain playing a role in the modulation of NMDAR currents (PubMed:33428810). {ECO:0000269\|PubMed:15632147, ECO:0000269\|PubMed:19033659, ECO:0000269\|PubMed:33428810}. | *Other* | $MD_{MD}$ |
| Q9NQV7 | *PRDM9 PFM6* | FUNCTION: Histone methyltransferase that sequentially mono-, di-, and tri-methylates both 'Lys-4' (H3K4) and 'Lys-36' (H3K36) of histone H3 to produce respectively trimethylated 'Lys-4' (H3K4me3) and trimethylated 'Lys-36' (H3K36me3) histone H3 and plays a key role in meiotic prophase by determining hotspot localization thereby promoting meiotic recombination (PubMed:24634223, PubMed:24095733, PubMed:26833727, PubMed:27129774). Can also methylate all four core histones with H3 being the best substrate and the most highly modified (PubMed:24095733, PubMed:24634223, PubMed:26833727). Is also able, on one hand, to mono and di-methylate H4K20 and on other hand to trimethylate H3K9 with the di-methylated H3K9 as the best substrate (By similarity). During meiotic prophase, binds specific DNA sequences through its zinc finger domains thereby determining hotspot localization where it promotes local H3K4me3 and H3K36me3 enrichment on the same nucleosomes through its histone methyltransferase activity (PubMed:26833727). Thereby promotes double-stranded breaks (DSB) formation, at this subset of PRDM9-binding sites, that initiates meiotic recombination for the proper meiotic progression (By similarity). During meiotic progression hotspot-bound PRDM9 interacts with several complexes; in early leptonema binds CDYL and EHMT2 followed by EWSR1 and CXXC1 by the end of leptonema. EWSR1 joins PRDM9 with the chromosomal axis through REC8 (By similarity). In this way, controls the DSB repair pathway, pairing of homologous chromosomes and sex body formation (By similarity). Moreover plays a central role in the transcriptional activation of genes during early meiotic prophase thanks to H3K4me3 and H3K36me3 enrichment that represents a specific tag for epigenetic transcriptional activation (By similarity). In addition performs automethylation (By similarity). Acetylation and phosphorylation of histone H3 attenuate or prevent histone H3 methylation (By similarity). {ECO:0000250\|UniProtKB:Q96EQ9, ECO:0000269\|PubMed:24095733, ECO:0000269\|PubMed:24634223, ECO:0000269\|PubMed:26833727}. | *Other* | $WTD_{ML}$ |
| Q9NR71 | *ASAH2 HNAC1* | FUNCTION: Plasma membrane ceramidase that hydrolyzes sphingolipid ceramides into sphingosine and free fatty acids at neutral pH (PubMed:10781606, PubMed:16229686, PubMed:26190575). Ceramides, sphingosine, and its phosphorylated form sphingosine-1-phosphate are bioactive lipids that mediate cellular signaling pathways regulating several biological processes including cell proliferation, apoptosis and differentiation (PubMed:15946935, PubMed:19345744, PubMed:24798654). Also catalyzes the reverse reaction allowing the synthesis of ceramides from fatty acids and sphingosine (PubMed:11278489, PubMed:17475390). Together with sphingomyelinase, participates in the production of sphingosine and sphingosine-1-phosphate from the degradation of sphingomyelin, a sphingolipid enriched in the plasma membrane of cells (PubMed:16061940). Also participates in the hydrolysis of ceramides from the extracellular milieu allowing the production of sphingosine-1-phosphate inside and outside cells (By similarity). This is the case for instance with the digestion of dietary sphingolipids in the intestinal tract (By similarity). {ECO:0000250\|UniProtKB:Q9JHE3, ECO:0000269\|PubMed:10781606, ECO:0000269\|PubMed:11278489, ECO:0000269\|PubMed:15946935, ECO:0000269\|PubMed:16061940, ECO:0000269\|PubMed:16229686, ECO:0000269\|PubMed:17475390, ECO:0000269\|PubMed:19345744, ECO:0000269\|PubMed:24798654, ECO:0000269\|PubMed:26190575}. | *Other* | $WTD_{ML}$ |
| Q9NTJ5 | *SACM1L KIAA0851 SAC1* | FUNCTION: Phosphoinositide phosphatase which catalyzes the hydrolysis of phosphatidylinositol 4-phosphate (PtdIns(4)P) (PubMed:24209621, PubMed:27044890, PubMed:29461204, PubMed:30659099). Can also catalyze the hydrolysis of phosphatidylinositol 3-phosphate (PtdIns(3)P) and has low activity towards phosphatidylinositol-3,5-bisphosphate (PtdIns(3,5)P2) (By similarity). Shows a very robust PtdIns(4)P phosphatase activity when it binds PtdIns(4)P in a 'cis' configuration in the cellular environment, with much less activity seen when it binds PtdIns(4)P in 'trans' configuration (PubMed:29461204, PubMed:24209621, PubMed:30659099). PtdIns(4)P phosphatase activity (when it binds PtdIns(4)P in 'trans' configuration) is enhanced in the presence of PLEKHA3 (PubMed:30659099). {ECO:0000250\|UniProtKB:Q9ES21, ECO:0000269\|PubMed:24209621, ECO:0000269\|PubMed:27044890, ECO:0000269\|PubMed:29461204, ECO:0000269\|PubMed:30659099}. | *Other* | $MD_{MD}$ |
| Q9NUD9 | *PIGV* | FUNCTION: Alpha-1,6-mannosyltransferase involved in glycosylphosphatidylinositol-anchor biosynthesis. Transfers the second mannose to the glycosylphosphatidylinositol during GPI precursor assembly. {ECO:0000269\|PubMed:15623507, ECO:0000269\|PubMed:15720390}. | *Other* | $WTD_{ML}$ |
| Q9NV58 | *RNF19A RNF19* | FUNCTION: E3 ubiquitin-protein ligase which accepts ubiquitin from E2 ubiquitin-conjugating enzymes UBE2L3 and UBE2L6 in the form of a thioester and then directly transfers the ubiquitin to targeted substrates, such as SNCAIP or CASR. Specifically ubiquitinates pathogenic SOD1 variants, which leads to their proteasomal degradation and to neuronal protection. {ECO:0000269\|PubMed:11237715, ECO:0000269\|PubMed:12145308, ECO:0000269\|PubMed:12750386, ECO:0000269\|PubMed:15456787, ECO:0000269\|PubMed:16513638}. | *Other* | $WTD_{ML}$ |
| Q9NXL6 | *SIDT1* | FUNCTION: In vitro binds long double-stranded RNA (dsRNA) (500 and 700 base pairs), but not dsRNA shorter than 300 bp. Not involved in RNA autophagy, a process in which RNA is directly imported into lysosomes in an ATP-dependent manner, and degraded. {ECO:0000250\|UniProtKB:Q6AXF6}. | *Other* | $WTD_{ML}$ |
| Q9NYQ8 | *FAT2 CDHF8 KIAA0811 MEGF1* | FUNCTION: Involved in the regulation of cell migration (PubMed:18534823). May be involved in mediating the organization of the parallel fibers of granule cells during cerebellar development (By similarity). {ECO:0000250\|UniProtKB:O88277, ECO:0000269\|PubMed:18534823}. | *Other* | $WTD_{ML}$ |
| Q9UGI9 | *PRKAG3 AMPKG3* | FUNCTION: AMP/ATP-binding subunit of AMP-activated protein kinase (AMPK), an energy sensor protein kinase that plays a key role in regulating cellular energy metabolism. In response to reduction of intracellular ATP levels, AMPK activates energy-producing pathways and inhibits energy-consuming processes: inhibits protein, carbohydrate and lipid biosynthesis, as well as cell growth and proliferation. AMPK acts via direct phosphorylation of metabolic enzymes, and by longer-term effects via phosphorylation of transcription regulators. AMPK also acts as a regulator of cellular polarity by remodeling the actin cytoskeleton; probably by indirectly activating myosin. The AMPK gamma3 subunit is a non-catalytic subunit with a regulatory role in muscle energy metabolism (PubMed:17878938). It mediates binding to AMP, ADP and ATP, leading to AMPK activation or inhibition: AMP-binding results in allosteric activation of alpha catalytic subunit (PRKAA1 or PRKAA2) both by inducing phosphorylation and preventing dephosphorylation of catalytic subunits. ADP also stimulates phosphorylation, without stimulating already phosphorylated catalytic subunit. ATP promotes dephosphorylation of catalytic subunit, rendering the AMPK enzyme inactive. {ECO:0000269\|PubMed:14722619, ECO:0000269\|PubMed:17878938}. | *Other* | $WTD_{ML}$ |
| Q9UGP8 | *SEC63 SEC63L* | FUNCTION: Mediates cotranslational and post-translational transport of certain precursor polypeptides across endoplasmic reticulum (ER) (PubMed:22375059, PubMed:29719251). Proposed to play an auxiliary role in recognition of precursors with short and apolar signal peptides. May cooperate with SEC62 and HSPA5/BiP to facilitate targeting of small presecretory proteins into the SEC61 channel-forming translocon complex, triggering channel opening for polypeptide translocation to the ER lumen (PubMed:29719251). Required for efficient PKD1/Polycystin-1 biogenesis and trafficking to the plasma membrane of the primary cilia (By similarity). {ECO:0000250\|UniProtKB:Q8VHE0, ECO:0000269\|PubMed:22375059, ECO:0000269\|PubMed:29719251}. | *Other* | $WTD_{ML}$ |

| | | | | |
|---|---|---|---|---|
| Q9UHC7 | *MKRN1 RNF61* | FUNCTION: E3 ubiquitin ligase catalyzing the covalent attachment of ubiquitin moieties onto substrate proteins. These substrates include FILIP1, p53/TP53, CDKN1A and TERT. Keeps cells alive by suppressing p53/TP53 under normal conditions, but stimulates apoptosis by repressing CDKN1A under stress conditions. Acts as a negative regulator of telomerase. Has negative and positive effects on RNA polymerase II-dependent transcription. {ECO:0000269\|PubMed:16785614, ECO:0000269\|PubMed:19536131}. | *Other* | $WTD_{ML}$ |
| Q9UHK6 | *AMACR* | FUNCTION: Catalyzes the interconversion of (R)- and (S)-stereoisomers of alpha-methyl-branched-chain fatty acyl-CoA esters (PubMed:7649182, PubMed:10655068, PubMed:11060359). Acts only on coenzyme A thioesters, not on free fatty acids, and accepts as substrates a wide range of alpha-methylacyl-CoAs, including pristanoyl-CoA, trihydroxycoprostanoyl-CoA (an intermediate in bile acid synthesis), and arylpropionic acids like the anti-inflammatory drug ibuprofen (2-(4-isobutylphenyl)propionic acid) but neither 3-methyl-branched nor linear-chain acyl-CoAs (PubMed:7649182, PubMed:10655068, PubMed:11060359). {ECO:0000269\|PubMed:10655068, ECO:0000269\|PubMed:11060359, ECO:0000269\|PubMed:7649182}. | *Other* | $MD_{MD}$ |
| Q9UJA5 | *TRMT6 KIAA1153 TRM6 CGI-09* | FUNCTION: Substrate-binding subunit of tRNA (adenine-N(1)-)-methyltransferase, which catalyzes the formation of N(1)-methyladenine at position 58 (m1A58) in initiator methionyl-tRNA (PubMed:16043508). Together with the TRMT61A catalytic subunit, part of a mRNA N(1)-methyltransferase complex that mediates methylation of adenosine residues at the N(1) position of a small subset of mRNAs: N(1) methylation takes place in tRNA T-loop-like structures of mRNAs and is only present at low stoichiometries (PubMed:29107537, PubMed:29072297). {ECO:0000269\|PubMed:16043508, ECO:0000269\|PubMed:29072297, ECO:0000269\|PubMed:29107537}. | *Other* | $MD_{BTD}$ |
| Q9UKN1 | *MUC12 MUC11* | FUNCTION: Involved in epithelial cell protection, adhesion modulation, and signaling. May be involved in epithelial cell growth regulation. Stimulated by both cytokine TNF-alpha and TGF-beta in intestinal epithelium. {ECO:0000269\|PubMed:17058067}. | *Other* | $WTD_{ML}$ |
| Q9UMX9 | *SLC45A2 AIM1 MATP* | FUNCTION: Proton-associated glucose and sucrose transporter (By similarity). May be able to transport also fructose (By similarity). Expressed at a late melanosome maturation stage where functions as proton/glucose exporter which increase lumenal pH by decreasing glycolysis (PubMed:32966160, PubMed:35469906). Regulates melanogenesis by maintaining melanosome neutralization that is initially initiated by transient OCA2 and required for a proper function of the tyrosinase TYR (PubMed:32966160, PubMed:35469906). {ECO:0000250\|UniProtKB:P58355, ECO:0000269\|PubMed:18563784, ECO:0000269\|PubMed:18683857, ECO:0000269\|PubMed:32966160, ECO:0000269\|PubMed:35469906}. | *Other* | $MD_{MD}$ |
| Q9Y2C5 | *SLC17A4* | FUNCTION: Acts as a membrane potential-dependent organic anion transporter, the transport requires a low concentration of chloride ions (PubMed:22460716). Mediates chloride-dependent transport of urate (PubMed:22460716). Mediates sodium-independent high affinity transport of thyroid hormones including L-thyroxine (T4) and 3,3',5-triiodo-L-thyronine (T3) (PubMed:30367059, PubMed:34937426). Can actively transport inorganic phosphate into cells via Na(+) cotransport (PubMed:22460716). {ECO:0000269\|PubMed:22460716, ECO:0000269\|PubMed:30367059, ECO:0000269\|PubMed:34937426}. | *Other* | $WTD_{ML}$ |
| Q9Y2W2 | *WBP11 NPWBP SIPP1 SNP70* | FUNCTION: Activates pre-mRNA splicing. May inhibit PP1 phosphatase activity. {ECO:0000269\|PubMed:10593949, ECO:0000269\|PubMed:11375989, ECO:0000269\|PubMed:14640981}. | *Other* | $WTD_{ML}$ |
| Q9Y3E0 | *GOLT1B GCT2 GOT1A CGI-141 HDCMA3 9P UNQ432/ PRO793* | FUNCTION: May be involved in fusion of ER-derived transport vesicles with the Golgi complex. | *Other* | $MD_{MD}$ |
| Q9Y6F1 | *PARP3 ADPRT3 ADPRTL3* | FUNCTION: Mono-ADP-ribosyltransferase that mediates mono-ADP-ribosylation of target proteins and plays a key role in the response to DNA damage (PubMed:16924674, PubMed:20064938, PubMed:21211721, PubMed:21270334, PubMed:25043379, PubMed:24598253, PubMed:28447610, PubMed:19354255, PubMed:23742272). Mediates mono-ADP-ribosylation of glutamate, aspartate or lysine residues on target proteins (PubMed:20064938, PubMed:25043379). In contrast to PARP1 and PARP2, it is not able to mediate poly-ADP-ribosylation (PubMed:25043379). Involved in DNA repair by mediating mono-ADP-ribosylation of a limited number of acceptor proteins involved in chromatin architecture and in DNA metabolism, such as histone H2B, XRCC5 and XRCC6 (PubMed:16924674, PubMed:24598253). ADP-ribosylation follows DNA damage and appears as an obligatory step in a detection/signaling pathway leading to the reparation of DNA strand breaks (PubMed:16924674, PubMed:21211721, PubMed:21270334). Involved in single-strand break repair by catalyzing mono-ADP-ribosylation of histone H2B on 'Glu-2' (H2BE2ADPr) of nucleosomes containing nicked DNA (PubMed:27530147). Cooperates with the XRCC5-XRCC6 (Ku80-Ku70) heterodimer to limit end-resection thereby promoting accurate NHEJ (PubMed:24598253). Suppresses G-quadruplex (G4) structures in response to DNA damage (PubMed:28447610). Associates with a number of DNA repair factors and is involved in the response to exogenous and endogenous DNA strand breaks (PubMed:16924674, PubMed:21211721, PubMed:21270334). Together with APLF, promotes the retention of the LIG4-XRCC4 complex on chromatin and accelerate DNA ligation during non-homologous end-joining (NHEJ) (PubMed:21211721). May link the DNA damage surveillance network to the mitotic fidelity checkpoint (PubMed:16924674). Acts as a negative regulator of immunoglobulin class switch recombination, probably by controlling the level of AICDA /AID on the chromatin (By similarity). In addition to proteins, also able to ADP-ribosylate DNA: mediates DNA mono-ADP-ribosylation of DNA strand break termini via covalent addition of a single ADP-ribose moiety to a 5'- or 3'-terminal phosphate residues in DNA containing multiple strand breaks (PubMed:29361132, PubMed:29520010). {ECO:0000250\|UniProtKB:Q3ULW8, ECO:0000269\|PubMed:16924674, ECO:0000269\|PubMed:19354255, ECO:0000269\|PubMed:20064938, ECO:0000269\|PubMed:21211721, ECO:0000269\|PubMed:21270334, ECO:0000269\|PubMed:23742272, ECO:0000269\|PubMed:24598253, ECO:0000269\|PubMed:25043379, ECO:0000269\|PubMed:27530147, ECO:0000269\|PubMed:28447610, ECO:0000269\|PubMed:29361132, ECO:0000269\|PubMed:29520010}. | *Other* | $MD_{BTD}$ |
| Q9Y6R4 | *MAP3K4 KIAA0213 MAPKKK 4 MEKK4 MTK1* | FUNCTION: Component of a protein kinase signal transduction cascade. Activates the CSBP2, P38 and JNK MAPK pathways, but not the ERK pathway. Specifically phosphorylates and activates MAP2K4 and MAP2K6. {ECO:0000269\|PubMed:12052864, ECO:0000269\|PubMed:9305639}. | *Other* | $WTD_{ML}$ |
| A3QJZ7 | *PRAMEF2 7* | | | $MD_{MD}$ |

| | | |
|---|---|---|
| A6NFZ4 | *FAM24A* | $MD_{BTD}$ |
| A6NHY2 | *ANKDD1B* | $WTD_{ML}$<br>$MD_{MD}$ |
| H3BQB6 | *STMND1* | $WTD_{ML}$ |
| P08243 | *ASNS*<br>*TS11* | $MD_{BTD}$ |
| P15086 | *CPB1*<br>*CPB*<br>*PCPB* | $MD_{BTD}$ |
| P15088 | *CPA3* | $MD_{BTD}$ |
| Q14094 | *CCNI* | $MD_{MD}$ |
| Q15434 | *RBMS2*<br>*SCR3* | $MD_{MD}$ |
| Q68CR7 | *LRRC66* | $WTD_{ML}$ |
| Q7Z2Y8 | *GVINP1*<br>*GVIN1*<br>*VLIG1* | $MD_{BTD}$<br>$WTD_{KEY}$ |
| Q8IWF9 | *CCDC83*<br>*HSD9* | $MD_{MD}$ |
| Q8IXL9 | *IQCF2* | $MD_{BTD}$ |
| Q8IZP2 | *ST13P4*<br>*FAM10A4* | $MD_{MD}$ |
| Q8N7B9 | *EFCAB3* | $WTD_{ML}$<br>$MD_{MD}$<br>MDBTD |
| Q8N7C4 | *TMEM217*<br>*C6orf128* | $MD_{MD}$ |
| Q8N865 | *C7orf31* | $MD_{BTD}$ |
| Q8NCL8 | *TMEM116* | $WTD_{ML}$ |
| Q8TBF8 | *FAM81A* | $WTD_{ML}$ |
| Q8TEQ0 | *SNX29*<br>*RUNDC2A* | $WTD_{ML}$ |
| Q969K7 | *TMEM54*<br>*BCLP*<br>*CAC1* | $WTD_{ML}$ |
| Q96BP2 | *CHCHD1*<br>*C10orf34*<br>*MRPS37* | $MD_{MD}$ |
| Q9H425 | *C1orf198* | $MD_{MD}$ |
| Q9H972 | *C14orf93* | $MD_{MD}$ |
| Q9P2S6 | *ANKMY1*<br>*TSAL1*<br>*ZMYND13* | $MD_{MD}$ |
| Q9UKR5 | *ERG28*<br>*C14orf1*<br>*AD-011*<br>*HSPC288*<br>*x0006* | $WTD_{KEY}$ |
| Q9Y3D5 | *MRPS18C*<br>*CGI-134* | $WTD_{KEY}$ |
| Q9Y6X4 | *FAM169A*<br>*KIAA0888* | $MD_{BTD}$ |
